# A comprehensive identification protocol for postcranial bones of Southeast Asian Ruminantia. Part 2: Morphological criteria for small and medium-sized species

**DOI:** 10.64898/2026.10.23.761334

**Authors:** Supalak Mithong, Sirikanya Chantasri, Julien Claude, Papavee Seesod, Corentin Bochaton

## Abstract

The taxonomic identification of postcranial elements from small and medium-sized ruminants represents a significant challenge in zooarchaeological and Late Quaternary paleontological studies, particularly in biodiversity-rich regions such as continental Southeast Asia. This study presents a comprehensive osteological reference framework for the small and medium-sized Ruminantia of this region, based on a comparative sample of 228 specimens representing 14 species across three families (Bovidae, Cervidae, and Moschidae). Osteological characters were first described on a small initial specimen set and subsequently scored across the full sample. Although a small number of character states were found to be unique to specific taxa, most proved highly variable, limiting their individual identification potential. To address this, a Random Forest and a Naïve Bayes classifier were applied to assess the discriminatory power of character combinations across four taxonomic ranks (Family, Subfamily, Genus, and Species) on independent bones. Both approaches performed similarly, achieving mean identification accuracies of 87–86% at the family level, 69% at the subfamily level, 50% at the genus level, and 35–33% at the species level. The Naïve Bayes models were further deployed as an interactive web application, allowing users to perform bone identification based on any combination of observable character states from a single anatomical element. Our results highlight the limits of morphological identification of postcranial elements in this taxonomic and biogeographic context, while demonstrating that a transparent, reproducible, and statistically grounded approach can still represent a significant step forward for zooarchaeological and Late Quaternary paleontological studies in Southeast Asia and beyond.

## Introduction

In recent decades, archaeological research into past interactions between human populations and animals has increased markedly, paralleling the development of zooarchaeology into a mature research field. Numerous published studies now address a wide range of questions, including human subsistence strategies and techniques, economic organization, mobility patterns of human groups, seasonal exploitation of natural resources, food preferences and taboos, and cooking practices (Reitz & Wing, 2008). As a result, zooarchaeology has become a well-established discipline encompassing studies from diverse periods, regions, and animal groups. Yet despite this diversity of research topics, most zooarchaeological studies ultimately rely on a basic and fundamental operation: the taxonomic identification of bone and dental remains recovered from archaeological deposits. This task most often depends on bone and tooth morphology and on direct comparison between modern and archaeological specimens. Such an approach can, in some cases, lack robustness, show limited reproducibility, and prove difficult or impossible to apply in complex contexts. Indeed, it is mainly viable in regions and periods where only a limited range of taxa is expected in the archaeological record, thereby restricting both the number of comparative modern specimens required and the likelihood of encountering morphologically similar taxa. However, although zooarchaeology initially developed largely in the Northern Hemisphere—where wild vertebrate biodiversity is relatively limited—the field is increasingly expanding into historically underexplored regions, including tropical areas. This shift introduces new challenges for zooarchaeologists, who now work in contexts characterized by high modern biodiversity, a feature that is also reflected in the taxonomic diversity of archaeological faunal assemblages. These new challenges also expose the current limitations of traditional zooarchaeological bone identification approaches, which often lack clear, standardized, and reproducible protocols. Indeed, the morphological identifications of animal remains from archaeological sites implicitly assume that archaeological specimens closely conform to their sampled modern counterparts (LeFebvre & Sharpe, 2018), and that taxonomic nomenclature is accurate and kept up to date by the curator of the collections owning the reference skeletons. In addition, factors such as the availability and quality of comparative collections, analyst experience and individual practice, and an implicit understanding of “standard zooarchaeological procedures” (Reitz and Wing, 2008) strongly influence identification accuracy and reliability, ultimately rendering the identification process poorly reproducible. Analyst experience and inter-observer variability in specimen identification are among the greatest challenges in zooarchaeological practice (Wolverton, 2013).

Addressing these issues requires access to comprehensive comparative skeletal or invertebrate collections, as well as extensive training, practice, mentorship, and the development of clear and reproducible identification protocols (Driver, 2011; Wolverton, 2013; Lyman, 2019). Biomolecular methods—such as collagen peptide mass fingerprinting (ZooMS) and ancient DNA analyses—have recently been applied to the taxonomic identification of archaeological bone remains to bypass the shortcomings of the morphological identifications. Although these approaches often provide more reliable and precise identifications, they are generally more costly and time-consuming than morphological analyses and are constrained by poor DNA or collagen preservation (Buckley et al., 2010; Rizzi et al., 2012; Buckley, 2018). Another way to improve morphology-based bone identification is the development of visually diagnostic osteological criteria tested on multiple modern specimens (Bochaton et al., 2016; Hanot & Bochaton, 2018; Zhang et al., 2022), or the use of osteological characters described with the same level of precision commonly applied in paleontological studies. Such approaches overcome most of the drawbacks of the traditional comparative approach and would be particularly relevant in areas of complex biodiversity, such as the tropical regions.

Yet, such regions have been largely neglected regarding the development of reproducible identification methods. For instance, in continental Southeast Asia, a very rich area in terms of both modern biodiversity and archaeological record, both biomolecular (Wannajuk et al., 2013; Siripan et al., 2019) and morphological approaches (Wang et al., 2020; Rodrigo & Manamendra-Arachchi, 2022) remain poorly developed. Although several paleontological studies have been conducted on reptile species, some of which focus on archaeological assemblages (Bochaton et al., 2019, 2023, 2025; Claude et al., 2019), mammals, which represent the bulk of large vertebrate assemblages in the region, are still poorly investigated from a zooarchaeological point of view. Except for a few rare cases (Suraprasit et al., 2016), most mammal studies rely primarily on cranial and dental remains (Meijaard & Groves, 2004; Wattanapituksakul et al., 2018; Yudha et al., 2019; Zhang & Tong, 2024), elements that are often scarce in archaeological contexts. This situation represents a major limitation for the development of zooarchaeological and taphonomic research in Southeast Asia, where high vertebrate biodiversity combined with the general lack of comprehensive modern comparative skeletal collections makes robust and precise identification of large mammal postcranial remains—often a substantial component of archaeological assemblages—particularly challenging.

To directly address this limitation, we developed a clear and detailed identification protocol for the appendicular skeleton of all wild ruminant species from continental Southeast Asia, based on osteometric data and osteological characters. We focused on Ruminantia (mainly Bovidae and Cervidae) because they are strongly represented in most prehistoric deposits of the region and were consistently hunted by past hunter-gatherer groups. We developed the identification protocol across three complementary studies: one dealing with the osteometry of modern species from continental Southeast Asia (Seesod et al., 2026), and two focusing on their morphological characteristics—one centered on large-sized species (Chantasri et al., 2026), and the other on small and medium-sized species. The present paper corresponds to the latter study. It focuses on fourteen small and medium-sized ruminant species occurring in continental Southeast Asia to propose a comprehensive identification protocol for their appendicular skeleton based on reliability-tested osteomorphological criteria.

## Material and Methods

### Investigated skeletal specimens

The Ruminantia species included in this study were selected based on their documented occurrence in continental Southeast Asia (Lekagul & McNeely, 1988; Francis, 2019) or in recent paleontological studies (Suraprasit et al., 2016). All the small and medium-sized genera currently present in the region were sampled, except *Pseudoryx nghetinhensis* Dung, Giao, Chinh, Tuoc, Arctander & MacKinnon, 1993, for which no complete skeleton is available in museum collections. In a few cases, closely related species from outside the region were included to increase sample size at the generic level (e.g., *Capricornis crispus*, *Muntiacus reevesi*). The present sample includes most Ruminantia species occurring in Southeast Asia but excludes the very small *Tragulus* species, which are easily identifiable on the basis of their size alone (Seesod et al., 2026). It also excludes large-sized species that were investigated in a separate study. These “large-sized” species have been defined on the basis of their skeletal measurements and correspond to the genera *Bos*, *Bubalus*, and *Budorcas* (Seesod et al., 2026). As their size was found to rarely overlap with that of other taxa in the region, it is generally easy to exclude the attribution of their bones to smaller Southeast Asian ruminant species (Seesod et al., 2026). We thus take this opportunity to exclude them from the present study and investigate them separately, in order to facilitate the identification of osteological characters allowing the distinction of the different species. Regarding the *Muntiacus* species included in our sample, three of the five species currently present in continental Southeast Asia (Francis, 2019) were described within the past 30 years: *M. gongshanensis* in 1990 (Shi-lai et al., 1990), *M. truongsonensis* in 1998 (Giao et al., 1998), and *M. putaoensis* in 1999 (Amato et al., 1999). In the historical collections we examined, specimens were generally assigned to *M. muntjak* for Southeast Asian material and *M. reevesi* for Chinese material. We chose to retain these attributions, while acknowledging that each may encompass multiple species; nonetheless, they still provide reliable information at the genus level. Regarding serows and gorals, recent molecular studies (Mori et al., 2019; Li et al., 2020) have shown that *Naemorhedus griseus* (Milne-Edwards, 1871) is conspecific with *Naemorhedus goral* (Hardwicke, 1825), and that *Capricornis milneedwardsii* David, 1869 is a synonym of *Capricornis sumatraensis* (Bechstein, 1799) (Mori et al., 2019). Accordingly, the museum specimens we examined were attributed to these species. No skeletal specimens of *Capricornis rubidus* (David, 1869) or *Naemorhedus baileyi* Pocock, 1914—both valid species—were present in the collections visited. Regarding the genus *Moschus*, we considered the type species of the genus (*Moschus moschiferus*), as it was the only one available during the initial definition of the characters. This name was used to refer to all the species of the genus *Moschus* in older taxonomies, and the monophyly of this genus has never been questioned (Groves & Grubb, 2011). As with *Muntiacus*, these specimens might correspond to several species of the genus.

The investigated specimen sample corresponds to a total of 228 specimens representing 14 species (or species groups). Our datasets include the following taxa: *Axis axis* (N=27); *Axis porcinus* (N=12); *Capricornis crispus* (N=4); *Capricornis sumatraensis* (N=13); *Rucervus eldii* (N=15); *Cervus nippon* (N=21); *Rucervus schomburgki* (N=1); *Rusa unicolor* (N=23); *Elaphodus cephalophus* (N=12); *Moschus moschiferus* (N=19); *Muntiacus muntjak* (N=32); *Muntiacus reevesi* (N=20); *Naemorhedus goral* (N=14); and *Pseudois nayaur* (N=15). These specimens were obtained from five museum collections in France, Belgium, Germany, and the United States (see full details in Supplementary Material 01; Mithong et al., 2026).

### Osteological character definition and recording protocol

In the initial step of the study, one specimen of each species was compared in an attempt to define meaningful osteological differences between the various taxa. The specimens used in this first step are the following: *Axis axis* (MNHN-ZM-AC-1861-45); *Axis porcinus* (MNHN-ZM-AC-1908-531); *Capricornis crispus* (MNHN-ZM-AC-1877-574); *Capricornis sumatraensis* (MNHN-ZM-AC-1901-1350); *Rucervus eldii* (MNHN-ZM-AC-1979-80); *Cervus nippon* (MNHN-ZM-AC-1904-234); *Rucervus schomburgki* (ZMB 16032); *Rusa unicolor* (MNHN-ZM-AC-1878-13); *Elaphodus cephalophus* (USNM 256107); *Moschus moschiferus* (MNHN-ZM-AC-2010-658); *Muntiacus muntjak* (MNHN-ZM-AC-1875-693); *Muntiacus reevesi* (MNHN-ZM-AC-1979-36); *Naemorhedus goral* (MNHN-ZM-AC-1963-320); and *Pseudois nayaur* (MNHN-ZM-AC-1996-2097). These characters follow the anatomical nomenclature of Barone (1986) and were illustrated using 3D models of the bones observed in the initial specimen sample to which the following specimens have been added: *Axis porcinus* (MNHN-ZM-AC-1971-72, and ZSM 1969-63), *Capricornis sumatraensis* (MNHN-ZM-AC-1937-314, and ZSM 1908-402), *Elaphodus cephalophus* (MNHN-ZM-AC-1896-689, and ZMB 70004), *Moschus moschiferus* (ZSM 1961-71), *Muntiacus muntjak* (MNHN-ZM-AC-9545), *Naemorhedus goral* (MNHN-ZM-AC-1962-153, and ZMB Not numbered), *Pseudois nayaur* (MNHN-ZM-AC-1985-055), and *Rusa unicolor* (MNHN-ZM-AC-2013-28). These 3D models were produced using an EinScan SP V2 scanner and the Agisoft PhotoScan Professional Edition software. All the 3D models are provided as Supplementary Material (Supplementary Material 07; Mithong et al., 2026).

This initial set of osteological characters was then recorded across the full sample in order to assess their reliability and relevance. The resulting dataset (Supplementary Material 02, Supplementary Material 03; Mithong et al., 2026) was then analyzed following the protocol described below.

### Analytical protocol of the characters

The initial dataset underwent a preliminary cleaning step before analysis. During this step, we removed all character states that were very rare (occurring in fewer than 10% of the specimens of every species in which they were observed), and all ambiguous character states (which represented 2.7% of the recorded character states). This cleaning step was performed manually on the original scoring table prior to the analyses described below; the resulting dataset (Supplementary Material 02; Mithong et al., 2026) thus already reflects these exclusions. The remaining characters and character states were then evaluated by identifying those unique to a given taxon, as well as by using the maximum frequency of each character state within a taxon. This approach provided a general indication of their stability and relevance for taxonomic identification. However, due to the variability of the recorded characters among the different taxa, we had to conduct more elaborate analyses to assess their performance in identifying isolated bones from the subfossil record.

To estimate the discriminatory power of each skeletal element and character combination, a Random Forest classifier was applied to binary-encoded (dummy-variable) character states. Because subfossil skeletal elements are most commonly recovered in isolation, each anatomical part was analyzed independently. For each element, 1000 independent cross-validation iterations were performed for every possible character subset of size k, ranging from 1 to the maximum number of characters available for that element, as most present a different number of characters. Each iteration was based on a stratified random split retaining 20% of specimens as a test set, with the constraint that at least two taxa must be represented in the training set. Classification accuracy was evaluated across all four taxonomic ranks (Family, Subfamily, Genus, and Species). For all the analyses at the genus rank, *Cervus*, *Rucervus*, and *Rusa* were all considered as representing a single genus and referred to *Cervus* (*sensu lato*) to avoid duplicating at the genus rank the analyses already conducted at the species rank, these three genera together comprising only four species in our dataset.

Because the deployed identification application must accept any user-defined combination of characters as input — a scenario incompatible with the fixed feature space of a single pre-trained Random Forest, unlike the fixed-subset training/testing scheme used in the benchmark above — a weighted multinomial Naive Bayes framework was also implemented as the inference engine for each skeletal element and taxonomic rank, since it can natively incorporate any subset of the available evidence at prediction time. Its performance was evaluated under the same parameters as the Random Forest approach. For this latter approach, likelihood estimates were derived from the observed frequencies of each character state within each taxon and the discriminatory power of each character (estimated by their information gain). In both the RF and NB approaches, for each identification query, analyses are conducted independently for each taxonomic rank (Family, Subfamily, Genus, and Species). While this approach may occasionally produce discrepancies between probability estimates at different ranks, it ensures that the attribution probabilities at each rank are as unbiased and internally consistent as possible given the available reference data. This approach has been preserved in the web application integrating the NB approach (see below).

All the analyses were performed in the software R v. 4.6.0 (R Core Team, 2026) (https://cran.r-project.org), with the packages dplyr (Wickham, François, et al., 2026), stringr (Wickham, 2025), ranger (Wright et al., 2026), readr (Wickham, Hester, et al., 2026), and tidyverse (Wickham et al., 2019). The NB approach was conducted with a homemade function. Coding assistance was provided by the generative AI model Claude (Anthropic), but all code was reviewed and validated by the authors prior to use. All the code necessary to reproduce these analyses is provided as supplementary information (Supplementary Material 04, Mithong et al., 2026).

The weighted multinomial Naïve Bayes models were exported as static parameter sets and deployed within a self-contained, browser-based identification HTML application (Bochaton et al., 2026). The prediction algorithm was reimplemented in JavaScript from the original R code to reproduce the R models’ probability estimates exactly. Every reference specimen was submitted to the deployed models for each anatomical element it is scored for. For each element, we tested every number of characters that can be entered at once, from a single character up to the maximum available on that bone, and for each of these we drew up to 100 distinct combinations of the character states recorded, to reproduce the range of situations a user may face (Supplementary Material 04, Mithong et al., 2026). The probabilities returned were then compared with whether the attribution was correct. Reliability rises steeply with the probability returned, and exceeds 90% from 80% upwards at the family, subfamily and genus ranks, and from 95% upwards at the species rank. The application applies these two thresholds and highlights the lowest rank that reaches its own; where none does, no attribution is highlighted. Instructions on how to use this application and how to combine the results with those of the previous osteometric study (Seesod et al., 2026) are provided in the Discussion section.

## Results

### Character descriptions

In this part, we describe and figure all the osteological features observed and included in the subsequent analyses. Unless otherwise stated, the figured specimens are the following: *Axis axis* (MNHN-ZM-AC-1861-45); *Axis porcinus* (MNHN-ZM-AC-1971-72); *Capricornis sumatraensis* (MNHN-ZM-AC-1901-1350); *Rucervus eldii* (MNHN-ZM-AC-1979-80); *Cervus nippon* (MNHN-ZM-AC-1904-234); *Rucervus schomburgki* (ZMB 16032); *Rusa unicolor* (MNHN-ZM-AC-1878-13); *Elaphodus cephalophus* (ZMB 70004); *Moschus moschiferus* (MNHN-ZM-AC-2010-658); *Muntiacus muntjak* (MNHN-ZM-AC-1875-693); *Muntiacus reevesi* (MNHN-ZM-AC-1979-36); *Naemorhedus goral* (MNHN-ZM-AC-1963-320); and *Pseudois nayaur* (MNHN-ZM-AC-1996-2097). The figured specimens are depicted with the sole intent of figuring characters, and the reader should keep in mind that these characters are, in nearly all cases, not unique to the figured species.

### Characters of the scapula

1. **Shape of supraspinous fossa margin in lateral view. (Fig. 1)**

A. It is of trapezoid shape with a pointed anterior margin.
B. It is inclined ventrally with a straight anterior edge.
C. It presents a dorsal expansion with a rounded anterior edge.
D. It is of trapezoid shape with a prominent dorsal expansion and a reduced ventral part.
2. **Shape of dorsal margin in lateral view. (Fig. 1)**

A. It is inclined, with the cranial angle being higher than the caudal angle.
B. It is straight.
C. It is concave.
D. It is inclined and slightly concave, with the cranial angle being lower than the caudal angle.
E. It is curved and inclined posteriorly.
3. **Shape of the lateral margin of the scapular spine in anterior view. (Fig. 2)**

A. It is regularly curved.
B. It is slightly expanded in the middle with a straight ventral margin and a strongly inclined dorsal area.
4. **Shape of the ventral margin of the scapular spine in anterior view. (Fig. 2)**

A. It is straight, perpendicular to the scapular collar, and the extremity of the spine is also perpendicular to the scapular collar.
B. It is slightly concave, and the extremity of the spine is perpendicular to the scapular collar.
C. It is straight, forms an acute angle with the scapular collar, and the extremity of the spine is turned up in posterior direction.
D. It is strongly concave, and the extremity of the spine is perpendicular to the scapular collar.
5. **Morphology of scapular collar in anterior view. (Fig. 2)**

A. It is short.
B. It is moderately elongated.
C. It is strongly elongated.
6. **Shape of the coracoid process in ventral view. (Fig. 3)**

A. It is wide, poorly developed, and weakly expanded in medial direction.
B. It is narrow and moderately expanded in medial direction.
C. It is elongated, well-developed in anterior direction, and expanded in medial direction.
D. It is wide, well-developed, and weakly expanded in medial direction.
E. It is short, weakly developed, and blunt.
7. **Shape of the glenoid cavity ventral margin in anterolateral view. (Fig. 4)**

A. It is slightly curved.
B. It presents an anterior angle and a straight posterior area.
C. It is sinusoidal.
D. It presents an anterior angle and a depressed posterior area.
8. **Occurrence of a bone extension on the anterodorsal extremity of the bone in lateral view. (Fig. 1)**

A. Present.
B. Absent. Characters of the humerus
9. **Occurrence of a supratrochlear foramen in anterior view. (Fig. 5)**

A. The supratrochlear foramen is present.
B. The supratrochlear foramen is absent.
10. **Development of the deltoid tuberosity in anterior view. (Fig. 5)**

A. It is well developed.
B. It is weakly developed.
C. It is moderately developed.
11. **Shape of the major tubercle of the humerus in posterior view. (Fig. 6)**

A. It is narrow with a rounded tip pointing in medial direction, and there is a notch on its lateral margin.
B. It is narrow with a pointed tip pointing in dorsomedial direction, and there is a notch on its lateral margin.
C. It is wide with a rounded tip and there is a notch on the lateral margin.
D. It is wide, and there is no notch on the lateral margin.
E. It is wide and well extended in dorsal direction with a rounded tip and a notch on its lateral margin.
12. **Shape of humerus diaphysis in medial view. (Fig. 7)**

A. The posterior and anterior margins of the diaphysis are strongly curved.
B. The posterior and anterior margins of the diaphysis are moderately curved.
C. The posterior margin of the diaphysis is weakly curved, and the anterior margin is straight.
13. **Morphology of minor tubercle in dorsal view. (Fig. 8)**

A. The dorsal margin is thin and sharp.
B. The dorsal margin is broad and blunt, and the medial margin is expanded in posteromedial direction.
C. The dorsal margin is broad and blunt, and the medial part is well extended in medial direction.
D. The dorsal margin is broad, extended in posterior direction, and of rectangular shape.
E. The dorsal margin is thin and strongly extended in dorsal and posterior directions.
14. **The shape of the medial epicondyle in ventral view. (Fig. 9)**

A. It is poorly developed, poorly individualized, slightly extended in lateral direction, and presents a curved medial margin.
B. It is moderately developed, poorly individualized, slightly extended in lateral direction, and presents an oblique medial margin.
C. It is moderately developed, moderately individualized, slightly extended in posterolateral direction, and presents an oblique medial margin.
D. It is well-developed, well-individualized, well extended and strongly inclined in posterolateral direction, and presents a curved medial margin. Characters of the radius
15. **Shape of the radius diaphysis in anterior view. (Fig. 10)**

A. It is narrow.
B. It is broad.
16. **Morphology of the styloid radial process in anterior view. (Fig. 10)**

A. It is weakly developed and blunt.
B. It is pointed and extended in ventral direction.
17. **Shape of medial articular surface in ventral view. (Fig. 11)**

A. It is wide with a posteromedial extension.
B. It is narrow with a blunt posteromedial margin.
C. It is wide, extended in medial direction, and has a pointed posterior margin.
D. It is blunt, with a depressed medial margin.
E. It is wide, of triangular shape, with a pointed posterior margin.
18. **Morphology of the groove of the extensor carpi radialis in ventral view. (Fig. 11)**

A. It is strongly depressed.
B. It is moderately depressed.
C. It is weakly depressed.
D. It is domed (not figured).
19. **Shape of the lateral margin of the proximal articular surface of the radius in dorsal view. (Fig. 12)**

A. It presents a non-articular lateral extension.
B. It is lacking a lateral extension.
20. **Occurrence of a concavity in the dorsal medial margin of the diaphysis in anterior view. (Fig. 13)**

A. Present.
B. Absent.
21. **Shape of coronoid process in anterior view. (Fig. 13)**

A. It is well developed and pointed.
B. It is weakly developed. Characters of the ulna
22. **Shape of the posterior margin of the olecranon tuberosity in medial view. (Fig. 14)**

A. Straight.
B. Projected in posterior direction.
23. **Shape of the olecranon process in medial view. (Fig. 14)**

A. It is narrow and short.
B. It is wide and moderately elongated.
C. It is narrow and elongated.
D. It is wide and short.
24. **Shape of anterodorsal margin of olecranon process in medial view. (Fig. 14)**

A. It forms a right angle.
B. It is curved.
C. It is beak-shaped.
25. **Shape of the summit of the olecranon process in medial view. (Fig. 14)**

A. It presents a triangular dorsal projection.
B. It is blunt.
C. It presents a moderately developed dorsal projection with a rounded dorsal margin. Characters of the scaphoid bone
26. **Shape of the posterodorsal area in lateral view. (Fig. 15)**

A. It is moderately extended in the posterior direction and presents a pointed process expanding in dorsal direction.
B. The whole area is extended posteriorly and is slightly pointed in dorsal direction.
C. The whole area presents a blunt posterodorsal margin and is moderately extended in posterior direction.
D. It is weakly extended in the posterior direction and presents a blunt posterodorsal margin.
E. It is not extended in the posterior direction and presents a pointed dorsal margin.
27. **The shape of the posteroventral articular surface in medial view. (Fig. 16)**

A. The facet is oriented in posterodorsal direction and is projected medially.
B. The facet is oriented in medial direction and not projected.
28. **Shape of scaphoid in posterior view (Fig. 17)**

A. The posterior face of the bone is flat, and its medial margin is strongly concave at mid-length. It also presents a well-marked dorsal expansion.
B. The area is flat, thin, with a concave lateral margin, and extends in the lateroventral direction.
C. The area is oval in shape with an inflated lateral margin.
D. The area is oval-shaped with a strong medioventral projection.
E. The posterior face of the bone is flat. The medial margin is strongly concave at mid-length. The dorsal region shows minimal dorsal expansion, while the medial area features a pointed medial projection.
F. The area is oval in shape without any strong dorsal or medial projection.
29. **Occurrence of a fossa on the dorsal margin of the bone in lateral view. (Fig. 15)**

A. Present.
B. Absent.
30. **Shape of scaphoid in ventral view (Fig. 18)**

A. It is triangular and narrow with a pointed posterior area strongly extended in posteromedial direction.
B. It is triangular and wide with a pointed posterior area strongly extended in posteromedial direction.
C. It is rectangular with a postero-medial area moderately extended in posteromedial direction and no posterolateral extension.
D. It is rectangular, with a postero-medial area slightly extended in posteromedial direction, and the lateral margin presents a well-developed posterolateral expansion.
E. It is rectangular, with a postero-medial area slightly extended in posteromedial direction, and the lateral margin presents a small posterolateral expansion.
F. It is nearly square shape, wide, and with very limited posteromedial and posterolateral expansions.
31. **Shape of the ventral margin of the bone in lateral view. (Fig. 15)**

A. The anteroventral area is more or less pointed and extended in ventral direction.
B. It has a blunt anteroventral area. Characters of the lunatum bone
32. **Shape of dorsal margin in lateral view. (Fig. 19)**

A. It is concave with raised anterior and posterior margins.
B. It is straight and inclined.
C. It is concave with a high and pointed anterior area and no raised posterior margin.
33. **The shape of the antero-dorsal area of the dorsal margin of the bone in lateral view. (Fig. 19)**

A. It is slightly pointed and weakly extended in dorsal direction.
B. It forms a prominent spine.
C. It is blunt.
34. **Shape of posterior margin in lateral view. (Fig. 19)**

A. It is triangular and well extended in posterior direction.
B. It is nearly flat and weakly extended in posterior direction.
35. **Morphology of the laterodorsal projection on the lateral margin of the bone in anterior view. (Fig. 20)**

A. It is well developed.
B. It is weakly developed.
36. **Morphology of the lateral margin of the bone in anterior view. (Fig. 20)**

A. Incurved in its ventral area.
B. Straight.
37. **Morphology of the medial margin of the bone in anterior view. (Fig. 20)**

A. It presents a deep depression at mid-length.
B. It presents a slight depression in its ventral half.
C. It is uniformly curved. Characters of the pyramidal bone
38. **Shape of the anterior articular surface with the lunatum bone in lateral view. (Fig. 21)**

A. It is depressed.
B. It is nearly flat.
39. **Shape of dorsal margin of the bone in medial view. (Fig. 22)**

A. It is curved.
B. It is pointed.
C. It is depressed.
40. **Morphology of the ventral process in lateral view. (Fig. 21)**

A. It is long with a blunt distal tip.
B. It is moderately long with a blunt distal tip.
C. It is long and has a pointed distal tip.
D. It is long with a pointed distal tip and presents a small pointed anterior expansion. Characters of the pisiform bone
41. **Shape of the pisiform in ventral view. (Fig. 23)**

A. The posterior area is widened and bent in medial direction.
B. It is thin, extended posteriorly, and moderately bent.
C. It is wide, moderately elongated, and weakly bent.
42. **Shape of dorsal outline of the bone in lateral view. (Fig. 24)**

A. It is domed and well extended in dorsal direction.
B. It is curved and moderately extended in dorsal direction.
C. It is flattened. Characters of the magnum bone
43. **Shape of posterior margin in dorsal view. (Fig. 25)**

A. It is strongly depressed.
B. It is weakly depressed.
C. It is flat.
D. It is moderately depressed.
44. **Morphology of the bone in dorsal view. (Fig. 25)**

A. It presents a medial ridge.
B. It has no medial ridge.
45. **Shape of the lateroventral projection in posterior view. (Fig. 26)**

A. It is weakly developed.
B. It is well developed and pointed in the lateroventral direction.
C. It is absent.
46. **Shape of the dorsomedial projection in posterior view. (Fig. 26)**

A. It is well developed and oriented in dorsal direction.
B. It is moderately developed and oriented in dorsal direction.
C. It is well developed and oriented in dorsomedial direction.
47. **Shape of the articular surface in ventral view. (Fig. 27)**

A. Its anterolateral margin is angular.
B. Its anterolateral margin is truncated.
C. Its anterolateral margin is curved.
48. **Morphology of the posterolateral projection in ventral view. (Fig. 27)**

A. It is well developed.
B. It is poorly developed or absent.
49. **Shape of magnum bone in medial view. (Fig. 28)**

A. It has an L shape with a well developed posterodorsal expansion.
B. It is nearly flat with a weakly developed posterodorsal expansion. Characters of the unciform
50. **Morphology of the anterior articular surface in medial view. (Fig. 29)**

A. It is well-expanded on the dorsal area of the bone.
B. It is slightly expanded on the dorsal area of the bone.
C. It is restricted to the posterior area of the bone and does not develop on the dorsal and ventral areas of the bone.
D. It expands only on the posterior area and on the whole ventral area margin of the bone.
51. **Shape of the anterior margin of the dorsal articular surface in dorsal view. (Fig. 30)**

A. It is depressed with an anteromedial projection.
B. It is strongly stepped with a flat medial portion.
C. It is nearly flat.
D. It presents a small expansion in anterolateral direction.
E. It is inclined in posterolateral direction.
52. **Morphology of the lateral margin of the bone in ventral view. (Fig. 31)**

A. It is angular with a well-developed lateral expansion.
B. It is curved with a flat posterior area.
C. It is homogeneously curved. Characters of the metacarpus
53. **Morphology of the sagittal ridges in posterior view. (Fig. 32)**

A. The ridges are strongly extended in dorsal direction above the limits of the articular surface.
B. The ridges are moderately extended in dorsal direction above the level of the articular surface.
C. The ridges do not extend above the level of the articulations.
54. **Shape of the lateral and medial distal margins of the bone in posterior view. (Fig. 32)**

A. The two margins bear moderately developed projections.
B. The two margins are nearly flat and parallel.
C. The two margins are slightly curved but do not present individualized lateral projections.
D. It has well-developed lateral projections.
55. **Morphology of the distal epiphysis in medial view. (Fig. 33)**

A. The width of the bone at the level of the sagittal ridge is similar to that at the level of the lateral extremity of the lateral articular eminence, and the distal margin of the sagittal ridge is sub-circular.
B. The width of the bone at the level of the sagittal ridge is clearly greater than that at the level of the lateral extremity of the lateral articular eminence, and the distal margin of the sagittal ridge is sub-circular.
C. The width of the bone at the level of the sagittal ridge is clearly greater than that at the level of the lateral extremity of the lateral articular eminence, and the distal margin of the sagittal ridge extends in ventral direction.
56. **Shape of the anterior outline of the proximal articular surface in dorsal view. (Fig. 34)**

A. It presents an anterior expansion, giving it a more rectangular shape.
B. It is rounded. Characters of the coxal
57. **Shape of the ischiatic tuberosity in lateral view. (Fig. 35)**

A. It is well-extended in posterodorsal direction with a triangular posterior margin.
B. It is extended in posterior direction and has a regularly curved posterior margin.
C. It is extended in posterior direction and has a beak-like tip.
58. **Shape of sciatic spine in dorsolateral view. (Fig. 36)**

A. It is flat.
B. It is curved.
59. **Morphology of the body of the ilium in dorsolateral view. (Fig. 36)**

A. It is elongated.
B. It is short and robust.
60. **Morphology of the ischiatic tuberosity in ventral view. (Fig. 37)**

A. It has blunt lateral and ventral tips that are poorly individualized.
B. The lateral tip and ventral tips are pointed and well-individualized.
61. **Morphology of the iliopubic eminence in ventral view. (Fig. 38)**

A. It is well developed, inflated, and well individualized.
B. It is moderately developed, thin, and not fully individualized.
C. It is weakly developed and weakly individualized. Characters of the femur
62. **Angle formed by the dorsal edge of the bone between the head of the femur and the greater trochanter in anterior view. (Fig. 39)**

A. It forms a clear acute angle.
B. It forms a clear obtuse angle.
C. It forms a nearly straight angle.
63. **Shape of the greater trochanter in anterior view. (Fig. 39)**

A. It is triangular, high, and its lateral margin is straight.
B. It is wide, short, and its lateral margin is curved.
C. It is wide, short, and presents a blunt tip on its lateral margin.
D. It is thin, short, and its lateral margin is curved.
E. It is triangular, short, and its lateral margin is straight.
F. It is thin, high, and its lateral margin is curved.
64. **Shape of the articular area of the head of the femur in dorsal view. (Fig. 40)**

A. It is short and rounded.
B. It is elongated and extended in medial direction.
65. **Morphology of the posterior margin of the neck of the femur in dorsal view. (Fig. 40)**

A. It presents a U-shape, short, and deep concavity.
B. It presents a long and regular concavity of moderate depth.
C. It presents a short concavity forming an indentation of moderate depth.
D. It presents a short and deep V-shape concavity.
66. **Anterior extension of the medial bulge of the anterior margin of the neck of the femur in dorsal view. (Fig. 40)**

A. It is weakly developed.
B. It is moderately developed.
C. It is well developed.
67. **Morphology of the distal part of the femur in medial view. (Fig. 41)**

A. The distal outline is long and strongly incurved, and the medial surface of the distal epiphysis is smooth and bears a crest running from the medial condyle to the diaphysis.
B. The distal outline is long and slightly curved with a depression located just anterior to the medial condyle.
C. The distal outline is short and moderately incurved.
68. **Shape of the anterior margin of the trochlea in ventral view. (Fig. 42)**

A. The medial lip of the trochlea is well extended in anterior direction, far beyond the lateral lip.
B. The medial lip of the trochlea is moderately extended in anterior direction, slightly beyond the lateral lip.
C. The medial lip of the trochlea is weakly extended in the anterior direction and is nearly at the same level as the lateral lip. Characters of the patella
69. **Thickness of the bone in medial view. (Fig. 43)**

A. The whole bone is inflated and very thick.
B. The anterodorsal area is thin and only slightly thicker than the ventral area of the bone.
C. The anterodorsal area is moderately inflated, around three times thicker than the ventral half.
70. **Morphology of the articular surface in medial view. (Fig. 43)**

A. It is nearly straight with a slight indentation in its middle part.
B. It is slightly and uniformly depressed.
C. It is strongly depressed.
71. **Shape of the ventral area in medial view. (Fig. 43)**

A. It is enlarged and short.
B. It is elongated.
72. **Shape of medial margin in anterior view. (Fig. 44)**

A. It presents a well-developed broad eminence with a blunt margin in its dorsal area.
B. It presents a moderately developed expansion in its dorsal area.
C. It is straight and oblique, giving the bone a triangular shape.
D. It lacks a dorsal enlargement, and the bone is oblong.
73. **Shape of anterodorsal margin of the anterior tuberosity in anterior view. (Fig. 44)**

A. It presents a broad crest running from its central area to its dorsal margin.
B. There is no dorsal crest.
C. It presents a broad and short crest restricted to the dorsal area.
D. It presents a thin and short crest restricted to the dorsal area.
74. **Shape of the dorsal margin of the articular surface in posterior view. (Fig. 45)**

A. It is curved or pointed.
B. It is flat.
75. **Morphology of the dorsomedial fossa in dorsal view. (Fig. 46)**

A. It is deep.
B. It is shallow.
C. It is absent. Characters of the tibia
76. **Shape of tibial tuberosity in dorsal view. (Fig. 47)**

A. It is concave.
B. It is weakly incurved or flat.
77. **Morphology of the lateral condyle in dorsal view. (Fig. 47)**

A. It is moderately extended in lateral direction.
B. It is well extended in lateral direction.
78. **Occurrence of an anterior projection on the lateral condyle in dorsal view. (Fig. 47)**

A. Yes.
B. No.
79. **Shape of the posterior intercondylar area in dorsal view. (Fig. 47)**

A. It is weakly extended in posterior direction.
B. It is well extended in posterior direction.
80. **Shape of tibial crest in lateral view. (Fig. 48)**

A. It is weakly developed.
B. It is well developed and presents a ventral indentation.
81. **Shape of anterior tubercle in ventral view. (Fig. 49)**

A. It is oval.
B. It is oval with two posterior projections.
C. It is oblong with two posterior projections.
D. It is oblong without posterior projections.
E. It has a crescent shape.
F. It is semicircular with two small posterior projections.
82. **Shape of the lateral tibial cochlea in ventral view. (Fig. 49)**

A. It is well extended in posterolateral direction in comparison to the medial tibial cochlea.
B. It is moderately extended in posterolateral direction in comparison to the medial tibial cochlea.
C. The lateral tibial cochlea is at the same level as the medial tibial cochlea. Characters of the malleolar bone
83. **Shape of the dorsal tip in medial view. (Fig. 50)**

A. It is narrow and well extended in dorsal direction with a pointed tip.
B. It is moderately extended with a pointed tip.
C. It is moderately extended with a blunt tip.
D. It is short.
84. **Shape of the ventral tip in medial view. (Fig. 50)**

A. It is well extended in ventral direction.
B. It is moderately extended in ventral direction.
C. It is weakly extended in ventral direction.
85. **Shape of the posterior margin in lateral view. (Fig. 51)**

A. It has a weak depression in the middle part.
B. It is curved.
C. It is pointed.
D. It is straight.
86. **Shape of the ventral margin in lateral view. (Fig. 51)**

A. It is curved.
B. It is pointed with a flat anteroventral margin.
C. It is pointed with a concave anteroventral margin.
D. It bears an anterior projection expanding in anteroventral direction. Characters of the calcaneus
87. **Shape of contact area with the posterior part of the astragalus in anterior view. (Fig. 52)**

A. It forms a right angle with the anterior branch of the calcaneus, and the fibular trochlea is located in posterior position.
B. The contact area with the anterior branch of the calcaneus is deeply curved and the fibular trochlea is located in anterior position.
C. The contact area with the anterior branch of the calcaneus is slightly curved, and the fibular trochlea is located in posterior position.
88. **Shape of the malleolar bone articular surface in anterior view. (Fig. 53)**

A. It is elongated with well-developed posterior and anterior facets.
B. It is elongated with a ventromedial projection.
C. It is short, with a weakly developed anterior part, and well individualized.
D. It is short with a ventromedial projection.
E. It is elongated, wide, and presents a posteromedial depression connected to the posterior articular facet. Characters of the astragalus
89. **Shape of the proximal trochlea groove in medial view. (Fig. 54)**

A. It is wide, deep, and its dorsal margin is flattened.
B. It is deep, and its dorsal margin is curved.
C. It is shallow and poorly delimited.
90. **Occurrence of an anterior indentation in the ventral margin in medial view. (Fig. 54)**

A. Present.
B. Absent.
91. **Morphology of posterior trochlea in medial view. (Fig. 54)**

A. Its posteroventral tip is strongly projected in ventral direction.
B. Its posteroventral tip is weakly projected in ventral direction.
92. **Shape of the anteroventral groove between the proximal trochlea and the plantar trochlea in posterior view. (Fig. 55)**

A. It is wide.
B. It is narrow.
C. It is wide in its medial area and narrow in its lateral area.
93. **Morphology of the lateral part of the proximal trochlea in posterior view. (Fig. 55)**

A. It has a pointed lateral tip.
B. It has a blunt lateral margin. Characters of the large cuneiform
94. **Morphology of the posterolateral margin of the articular surface in dorsal view. (Fig. 56)**

A. It is blunt.
B. It presents a wide, rounded posterolateral projection.
C. It presents a short triangular posterolateral projection.
95. **Shape of the dorsal and ventral margins in medial view. (Fig. 57)**

A. They are nearly parallel and flat.
B. They are incurved.
C. The dorsal margin is convex, and the ventral margin is incurved.
96. **Fusions with other bones. (Fig. 57)**

A. The large cuneiform is fused with the small cuneiform.
B. It is not fused with any bone. Characters of the navicular-cuboid
97. **The large cuneiform is fused with the navicular-cuboid. (Fig. 58)**

A. Yes.
B. No.
98. **Shape of the lateral articular surface in ventral view. (Fig. 59)**

A. It is triangular with a straight or slightly convex lateral margin.
B. It is elongated in posterolateral direction with a concave lateral margin.
C. It is triangular with a concave lateral margin.
D. It is triangular in shape, concave on its anterolateral margin, and presents a posterior constriction.
E. It has a concave lateral margin and a small and well-marked depression on its posterolateral margin.
99. **Shape of posterior articular surface in ventral view. (Fig. 59)**

A. It is triangular and well developed.
B. It is elongated and moderately developed.
C. It is triangular and moderately developed.
D. It is oval and weakly developed.
100. **Morphology of the posteromedial projection in ventral view. (Fig. 59)**

A. It is well-developed with a blunt posterior extremity.
B. It is moderately developed with a blunt posterior extremity.
C. It is weakly developed with a flat posterior margin.
D. It is very well-developed but narrow with a pointed posterior extremity. Characters of the metatarsus
101. **Shape of the ventromedial projection in dorsal view. (Fig. 60)**

A. It is wide, blunt, and well extended in posterior direction.
B. It is pointed and well extended in posterior direction.
C. It is pointed and moderately extended in posterior direction.
D. It is pointed and does not extend posteriorly beyond the posterior articular surface.
E. It is absent or weakly developed.
102. **Shape of the posterior articular surface in dorsal view. (Fig. 60)**

A. It is wide and occupies most of the posterior area of the bone.
B. It is oval, short, and located in a central position on the posterior area of the bone.
C. It is oblong and located in a central position on the posterior area of the bone.
D. It is oblong and located on the lateral side of the bone.
103. **Overall shape of proximal area. (Fig. 60)**

A. Rounded.
B. Triangular or rectangular.
104. **Morphology of the sagittal ridge in posterior view. (Fig. 61)**

A. It is long and extends nearly to the lateral edge of the bone.
B. It is well-extended above the dorsal limit of the condyle.
C. It is short and weakly extended above the dorsal limit of the condyle.
105. **Shape of the lateral and medial margins in posterior view. (Fig. 61)**

A. It has small lateral projections just above the dorsal margins of the condyles.
B. The margins are flat and lack lateral projections.
C. The margins are slightly curved and lack lateral projections.
D. The margins are well-curved and lack lateral projections.
106. **Shape of the distal metatarsal canal. (Fig. 62)**

A. The canal reaches the ventral limit of the bone.
B. The canal closes before reaching the ventral area of the bone.
107. **Morphology of the medial condyle in medial view. (Fig. 63)**

A. It is slightly smaller than the lateral condyle and is circular.
B. It is clearly smaller than the lateral condyle and is circular.
C. It is clearly smaller than the lateral condyle, and presents a reduced ventral area. Characters of the axial large sesamoid
108. **Shape of the bone in lateral view. (Fig. 64)**

A. It is triangular with a posteroventral area expanded in the posteroventral direction.
B. It is rectangular.
C. It is semicircular.
D. It is triangular with a pointed dorsal margin.
109. **Shape of ventral margin in lateral view. (Fig. 64)**

A. It has a deep and wide indentation.
B. It is slightly concave or with a very small and shallow indentation.
C. It is flat or convex. Characters of the first phalanx
110. **Shape of medial margin in posterior view. (Fig. 65)**

A. It is straight.
B. It is depressed in the middle.
C. It is straight with a dorsomedial projection.
111. **Morphology of the condyles in posterior view. (Fig. 65)**

A. The lateral condyle is extended dorsally above the level of the medial condyle.
B. Both the lateral and medial condyles extend dorsally at the same level.
C. The medial condyle is extended dorsally above the level of the lateral condyle.
112. **Shape of the distal articular surface in ventral view. (Fig. 66)**

A. Its anterior margin is curved.
B. The anterior margin is flat.
113. **Shape of the first phalanx in medial view. (Fig. 67)**

A. It is wide and triangular.
B. It is rectangular, moderately enlarged, and has a straight palmar face.
C. It is rectangular and narrow.
D. It is rectangular, moderately enlarged, and has a concave palmar face.
114. **Shape of the medial distal articular facet in medial view (Fig. 67)**

A. It is wide in comparison to the width of the body of the phalanx.
B. It is reduced in comparison to the width of the body of the phalanx.
115. **Morphology of the phalanx in anterior view. (Fig. 68)**

A. The whole phalanx is thin.
B. The whole phalanx is wide, the lateral condyle expands in lateral direction, and the medial condyle is more extended in ventral direction than the lateral condyle.
C. The whole phalanx is wide, the lateral condyle does not expand in lateral direction, and the medial condyle is more extended in ventral direction than the lateral condyle. Characters of the second phalanx
116. **Shape of the condyle surface in anterior view. (Fig. 69)**

A. It is moderately extended on the dorsal area of the phalanx.
B. It is well-extended on the dorsal area of the phalanx, and its extension has an angular dorsal margin.
C. It is well-extended on the dorsal area of the phalanx, and its extension has a pointed dorsal margin.
117. **Shape of the insertion tubercle in dorsal view. (Fig. 70)**

A. It is well developed with a pointed extremity.
B. It is moderately developed with a pointed extremity.
C. It is moderately developed and blunt.
118. **Shape of the proximal articular surface in dorsal view. (Fig. 70)**

A. It is rounded.
B. It is narrow.
C. It is wide and triangular.
119. **Shape of the second phalanx in axial view. (Fig. 71)**

A. It is wide.
B. It is narrow.
120. **Morphology of the proximal articular surface in axial view. (Fig. 71)**

A. The posteroaxial angle of the proximal articular surface strongly extends dorsally above the level of the anterior angle, and the posterior axial area of the proximal part is strongly enlarged.
B. The anterior and posteroaxial angles of the proximal articular surface are of similar dorsal extensions, and the posterior axial area of the proximal part is moderately enlarged.
C. The anterior angle of the proximal articular surface strongly extends dorsally above the level of the posteroaxial angle, and the posterior axial area of the proximal part is moderately enlarged.
121. **Shape of distal condyle in axial view. (Fig. 71)**

A. It is well-delimited, curved, short, and not extended on either the dorsal or palmar surface of the bone.
B. It is well-delimited, broad, curved, and well-extended on the palmar surface of the bone.
C. It is well-delimited, curved, and extended on the dorsal area of the bone only.
D. It is well-delimited, slightly flattened, curved, and extended on the palmar surface of the bone only.
E. It is curved, moderately delimited, and extended on the palmar surface of the bone only.
122. **Morphology of the insertion tubercle in abaxial view. (Fig. 72)**

A. It is triangular, more or less pointed and well-extended in posterodorsal direction.
B. It is short and not extended in any direction.
C. It is broad, angular, and weakly extended in posterodorsal direction.
D. It is triangular, and moderately oriented in dorsal direction.
123. **Shape of condyle in abaxial view. (Fig. 72)**

A. It is broad, centered on the axis of the anterodorsal margin of the phalanx, and with a subcircular anterior margin.
B. It is small, and its anterior margin has a slightly subtriangular shape.
C. It is broad with its centre slightly displaced in ventral direction and with an anterior margin of a truncated triangular shape.
D. It is small with its centre slightly displaced in ventral direction and with an anterior margin of circular shape.
124. **Shape of the distal articular surface in posterior view. (Fig. 73)**

A. It is narrow, and the lateral articular surface extends on the posterior area beyond the level of the medial articular surface.
B. It is narrow, and the lateral articular surface is less extended on the posterior area of the bone than the medial articular surface.
C. It is wide, the dorsolateral and dorsomedial margins of the condyle are compressed, and the lateral and medial articular surfaces are of similar dorsal development.
D. It is wide, and the lateral articular surface extends on the posterior area beyond the level of the medial articular surface.
125. **Ventral extension of the extremities of the distal articular surface in posterior view. (Fig. 73)**

A. The ventral extremities of the condyles are similarly extended in ventral direction.
B. The ventrolateral area of the condyle is pointing in ventral direction and is more extended than the ventromedial area.
126. **Shape of the distal articular surface in ventral view. (Fig. 74)**

A. It is triangular without a notch on its anterolateral margin.
B. It is rectangular.
C. It is triangular and bears a notch on its anterolateral margin. Characters of the third phalanx
127. **Shape of solar face in ventral view. (Fig. 75)**

A. It is narrow, elongated, and its medial margin is curved.
B. It is triangular with straight lateral and medial margins.
C. It is oblong with a pointed anterior tip.
D. It is semicircular, with a curved lateral margin and a straight medial margin, and a blunt anterior tip.
E. It is semicircular, with a curved lateral margin and a straight medial margin, and a pointed anterior tip.
F. It is oblong with a blunt anterior tip.
128. **Morphology of the extensor process in posterior view. (Fig. 76)**

A. It is well extended in dorsal direction.
B. It is moderately extended in dorsal direction.
C. It is weakly extended in dorsal direction.
129. **Shape of medial margin in posterior view. (Fig. 76)**

A. It is convex.
B. It is uniformly concave.
C. It is sinusoidal.
130. **Shape of the palmar process in lateral view. (Fig. 77)**

A. It is moderately extended in posterior direction.
B. It is strongly extended in posterior direction.
C. It is weakly extended in posterior direction.
131. **Shape of the apex in lateral view. (Fig. 77)**

A. It is pointed.
B. It is truncated.
132. **Morphology of the solar face in lateral view. (Fig. 77)**

A. It is straight or slightly concave.
B. It is straight.
133. **Shape of the extensor process in the lateral view. (Fig. 77)**

A. It is short and truncated.
B. It is pointed.

### II) Variability of the character states across the taxa

The frequency of the recorded characters across different taxonomic ranks (Supplementary Material 05; Mithong et al., 2026) shows that, despite our attempts to record characters unique to each species, most character states are highly variable and occur in several taxa at variable frequencies (**Figure 78**). Indeed, among our 133 characters, only one (131) is never polymorphic at the species level. The other characters present polymorphism with at least 1 species (8 characters), 2 species (4 characters), 3 species (6 characters), 4 species (4 characters), 5 species (5 characters), six species (6 characters), 7 species (10 characters), 8 species (10 characters), 9 species (13 characters), 10 species (17 characters), 11 species (22 characters), 12 species (15 characters), and 13 species (12 characters). This extremely high level of polymorphism impacting all postcranial elements strongly blurs the taxonomic and phylogenetic signal of our dataset.

**Figure 1.**
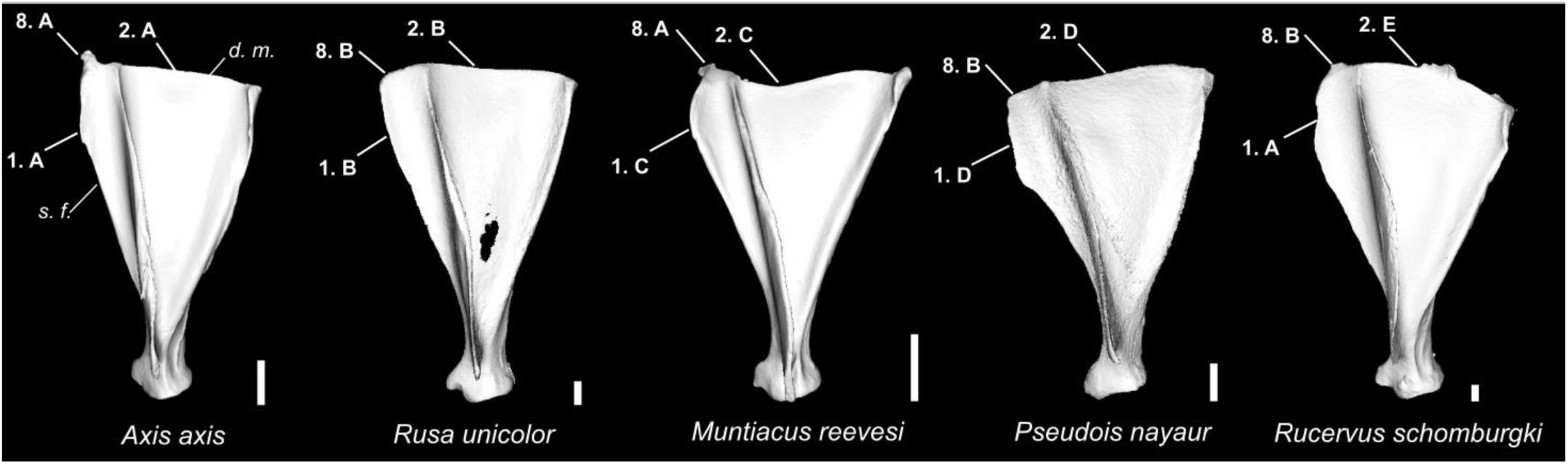
Character states visible on the lateral face of the left scapula (characters 1, 2, and 8). Abbreviations: s. f.: supraspinous fossa; d. m.: dorsal margin. Scale bar = 2 cm.

**Figure 2.**
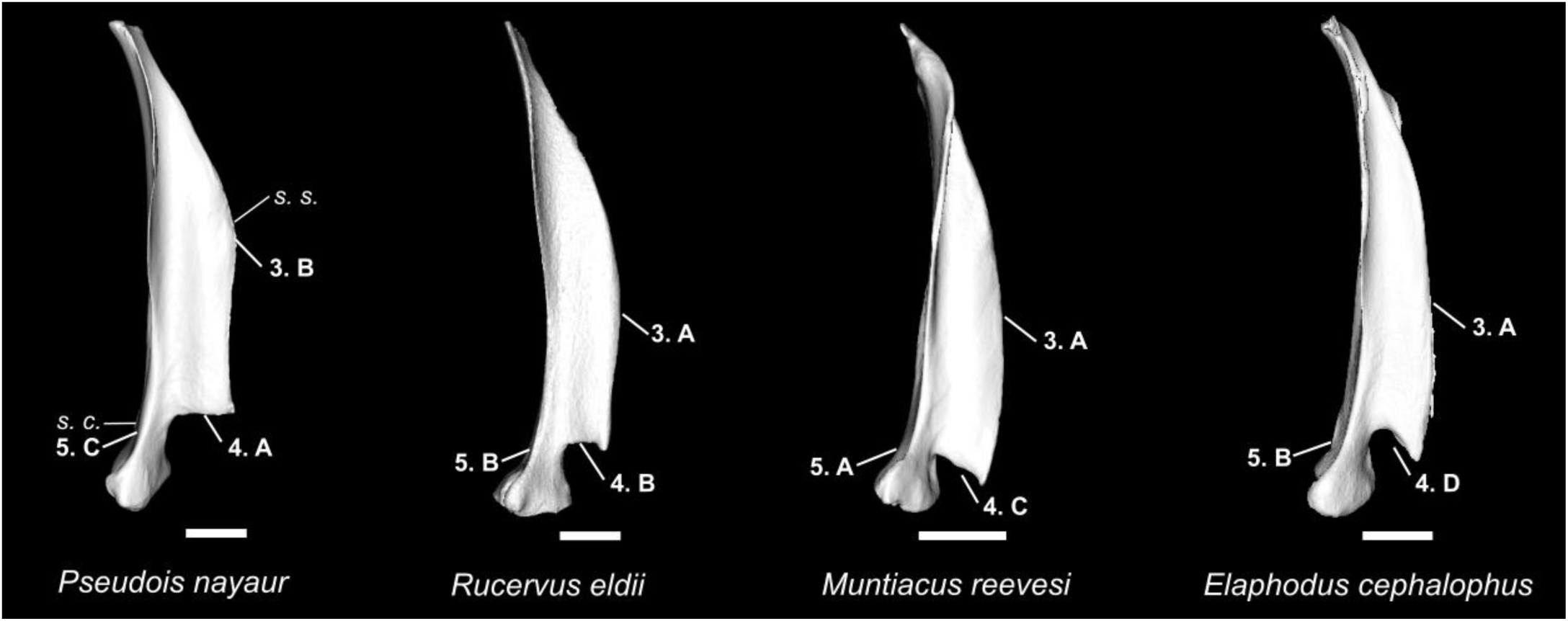
Character states visible on the anterior face of the left scapula (characters 3, 4, and 5). Abbreviations: s. s.: scapular spine; s. c.: scapular collar. Scale bar = 2 cm.

**Figure 3.**
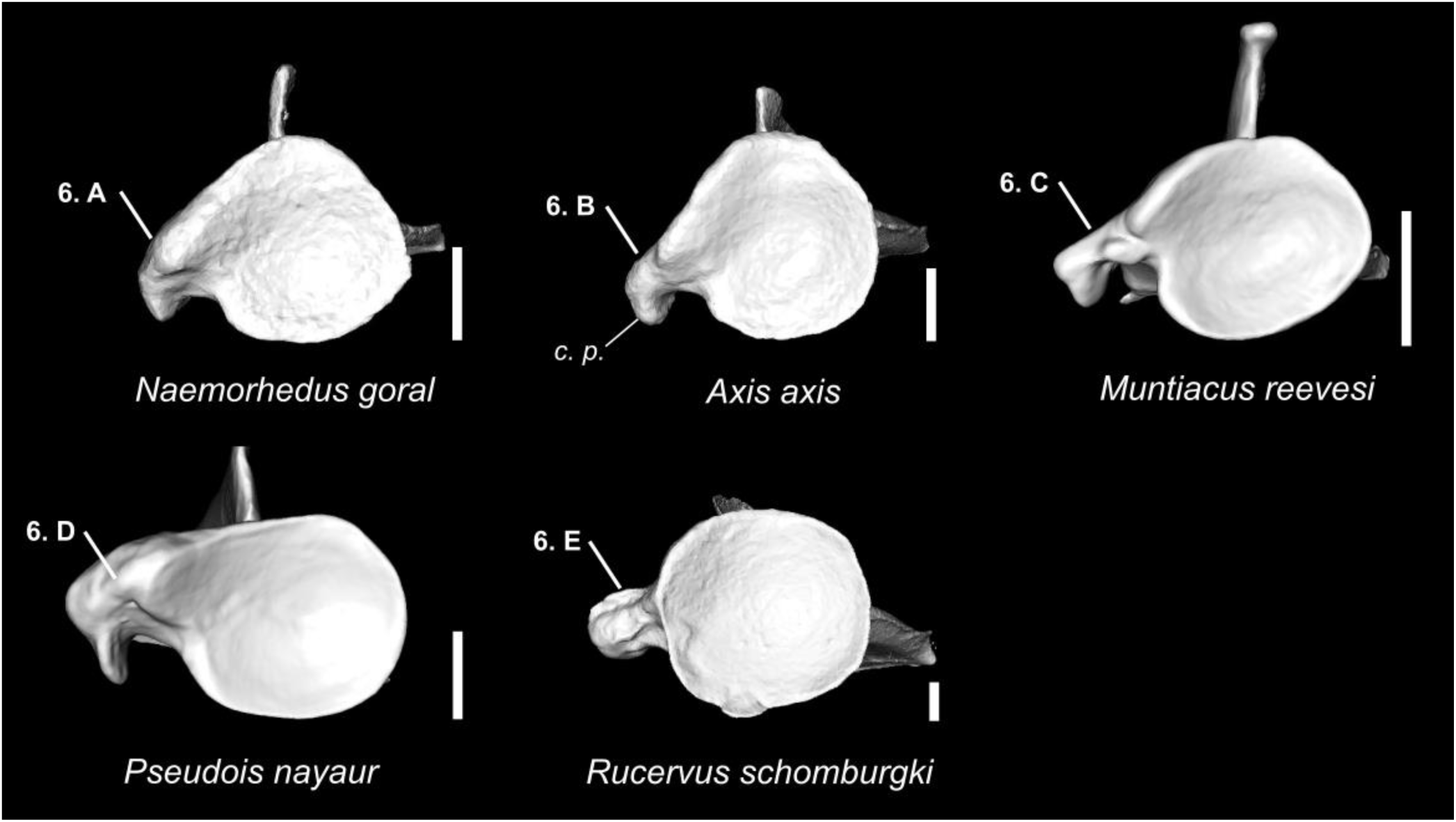
Character states visible on the ventral face of the left scapula (character 6). Abbreviations: c. p.: coracoid process. Scale bar = 1 cm.

**Figure 4.**
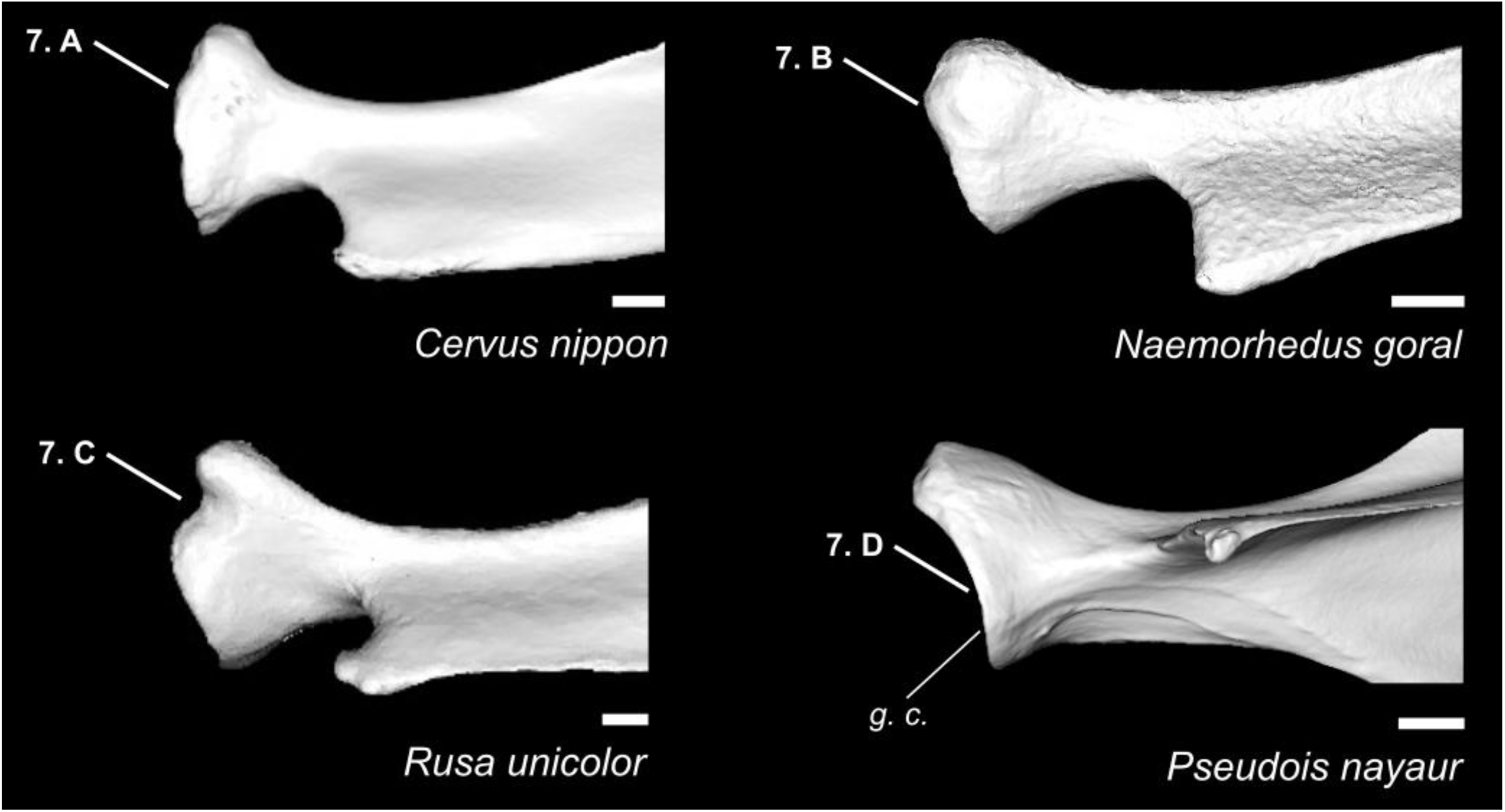
Character states visible on the anterolateral face of the left scapula (character 7). Abbreviations: g. c.: glenoid cavity. Scale bar = 1 cm.

**Figure 5.**
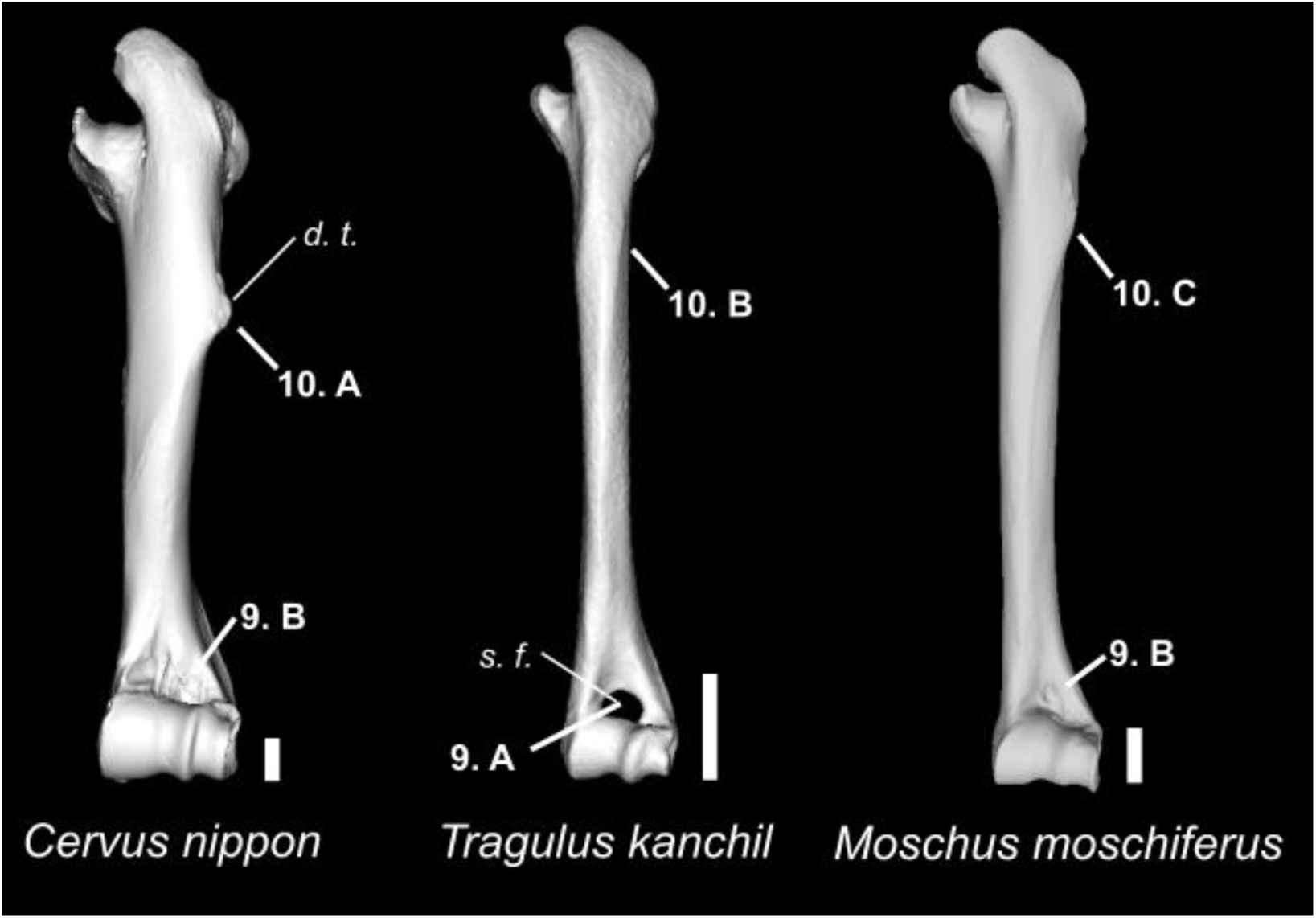
Character states visible on the anterior face of the left humerus (characters 9 and 10). Abbreviations: s. f.: supratrochlear foramen; d. t.: deltoid tuberosity. *Tragulus kanchil* is figured here because none of the specimens scanned for the species included in this study displayed these character states. Scale bar = 1 cm.

**Figure 6.**
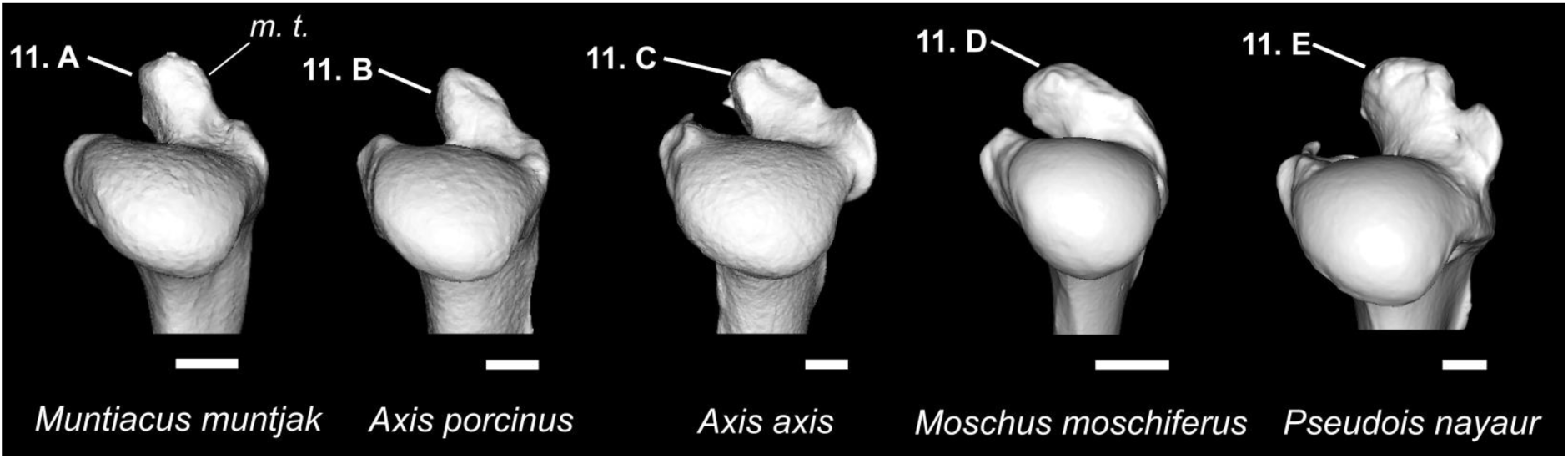
Character states visible on the posterior face of the right humerus (character 11). Abbreviations: m. t.: major tubercle. Scale bar = 1 cm.

**Figure 7.**
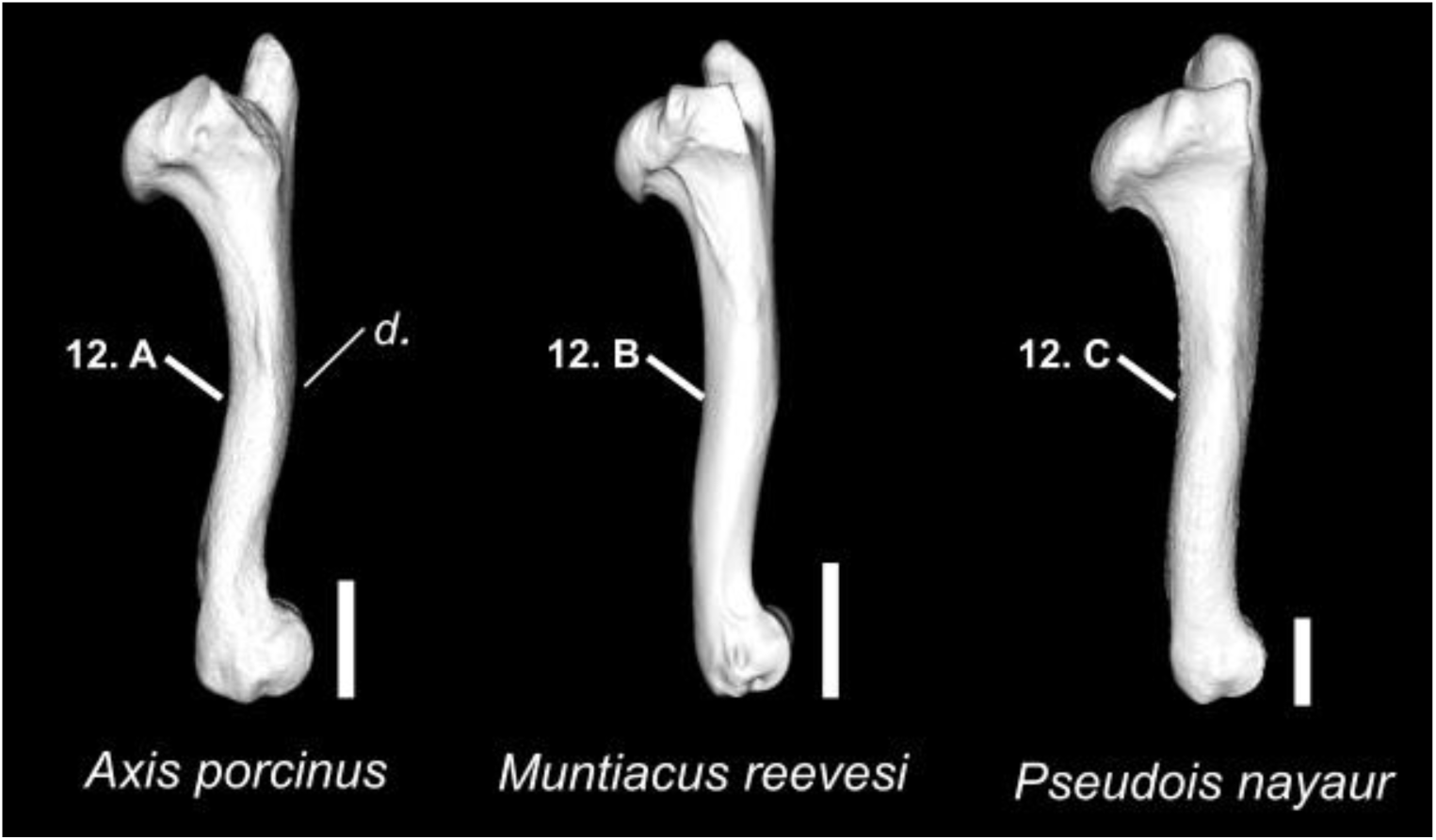
Character states visible on the medial face of the left humerus (character 12). Abbreviations: d.: diaphysis. Scale bar = 2 cm.

**Figure 8.**
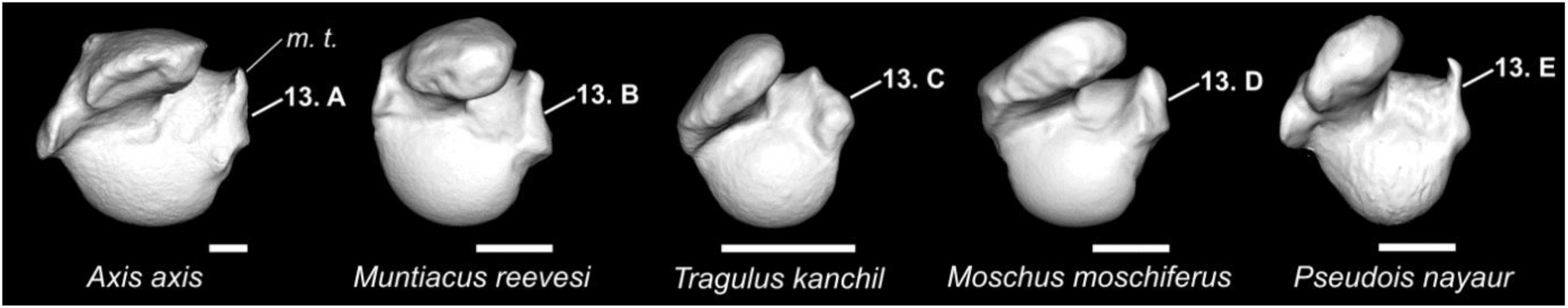
Character states visible on the dorsal face of the left humerus (character 13). *Tragulus kanchil* is figured here because none of the specimens scanned for the species included in this study displayed this character state. Abbreviations: m. t.: minor tubercle. Scale bar = 1 cm.

**Figure 9.**
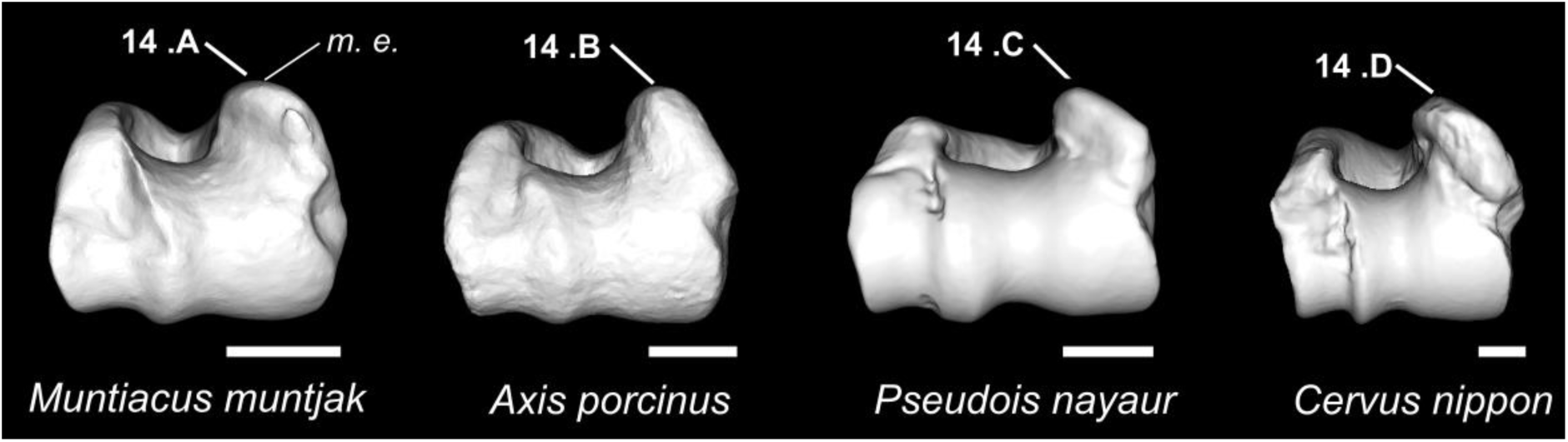
Character states visible on the ventral face of the left humerus (character 14). Abbreviations: m. e.: medial epicondyle. Scale bar = 1 cm.

**Figure 10.**
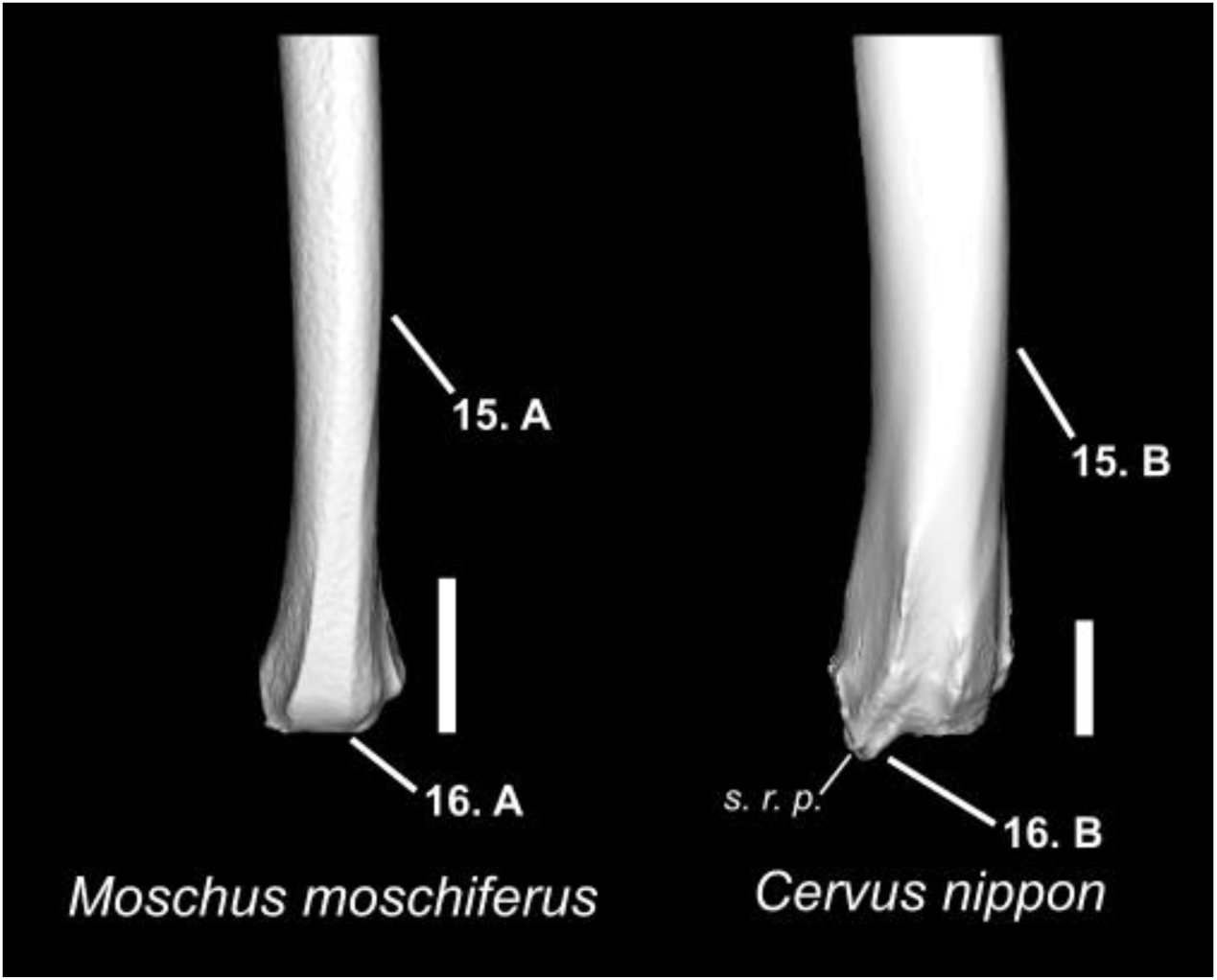
Character states visible on the anterior face of the left radius (characters 15 and 16). Abbreviations: s. r. p.: styloid radial process. Scale bar = 2 cm.

**Figure 11.**
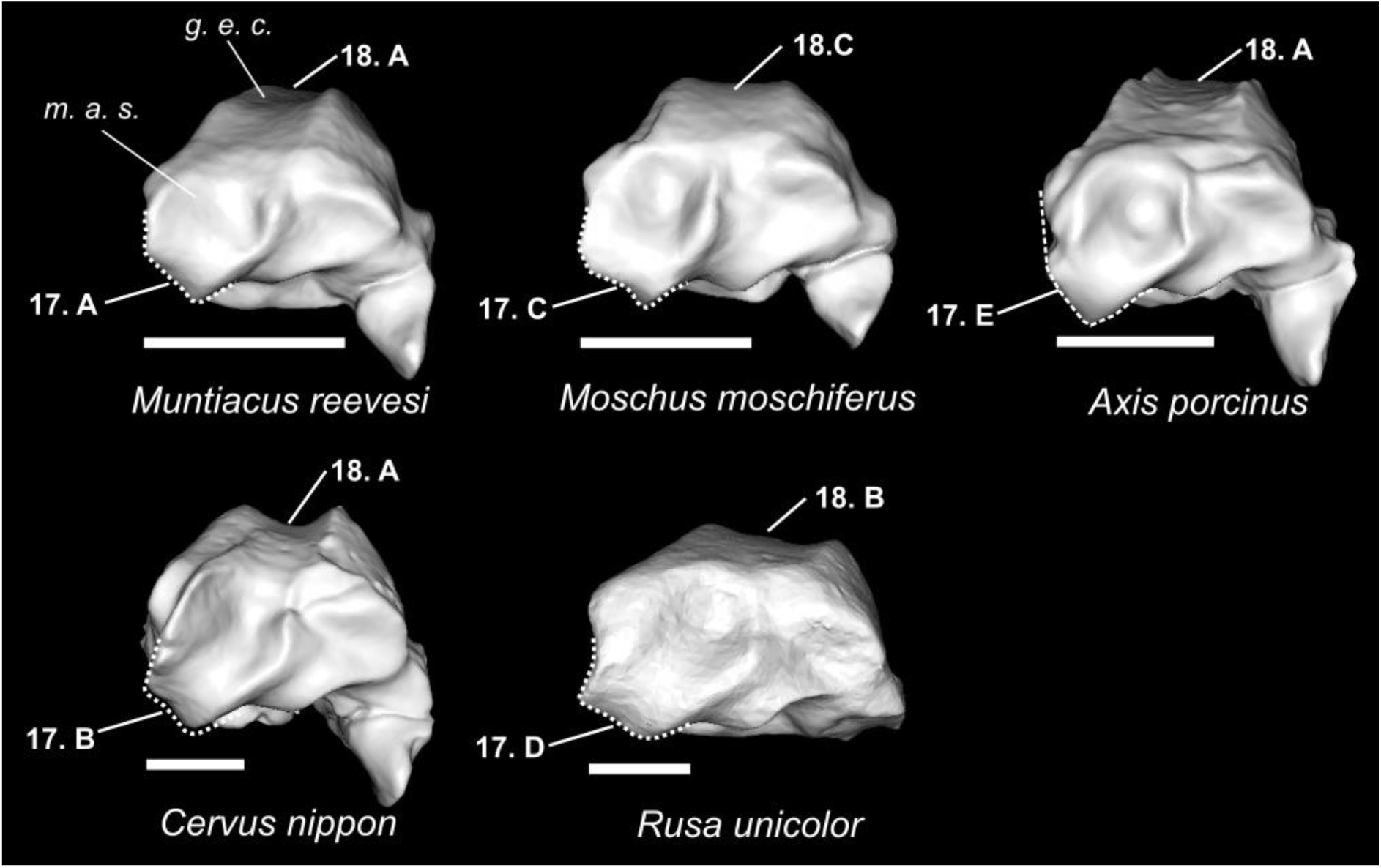
Character states visible on the ventral face of the right radius (characters 17 and 18). Abbreviations: m. a. s.: medial articular surface; g. e. c.: groove of the extensor carpi radialis. Scale bar = 1 cm.

**Figure 12.**
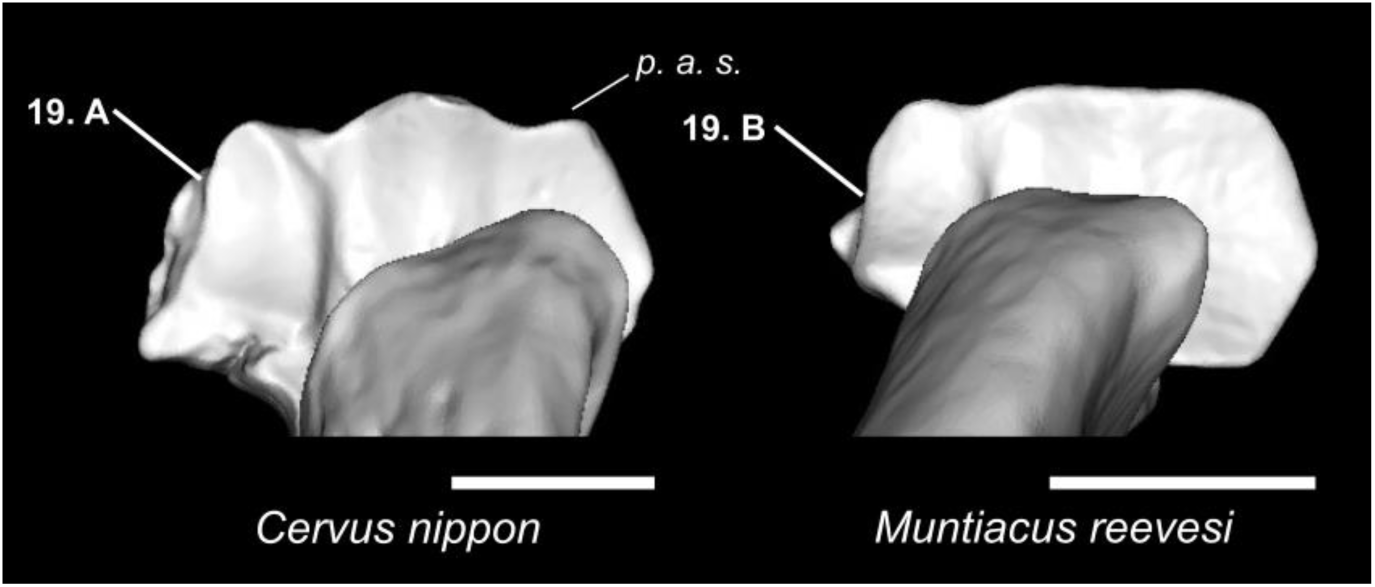
Character states visible on the dorsal face of the left radius (character 19). Abbreviations: p. a. s.: proximal articular surface. Scale bar = 1 cm.

**Figure 13.**
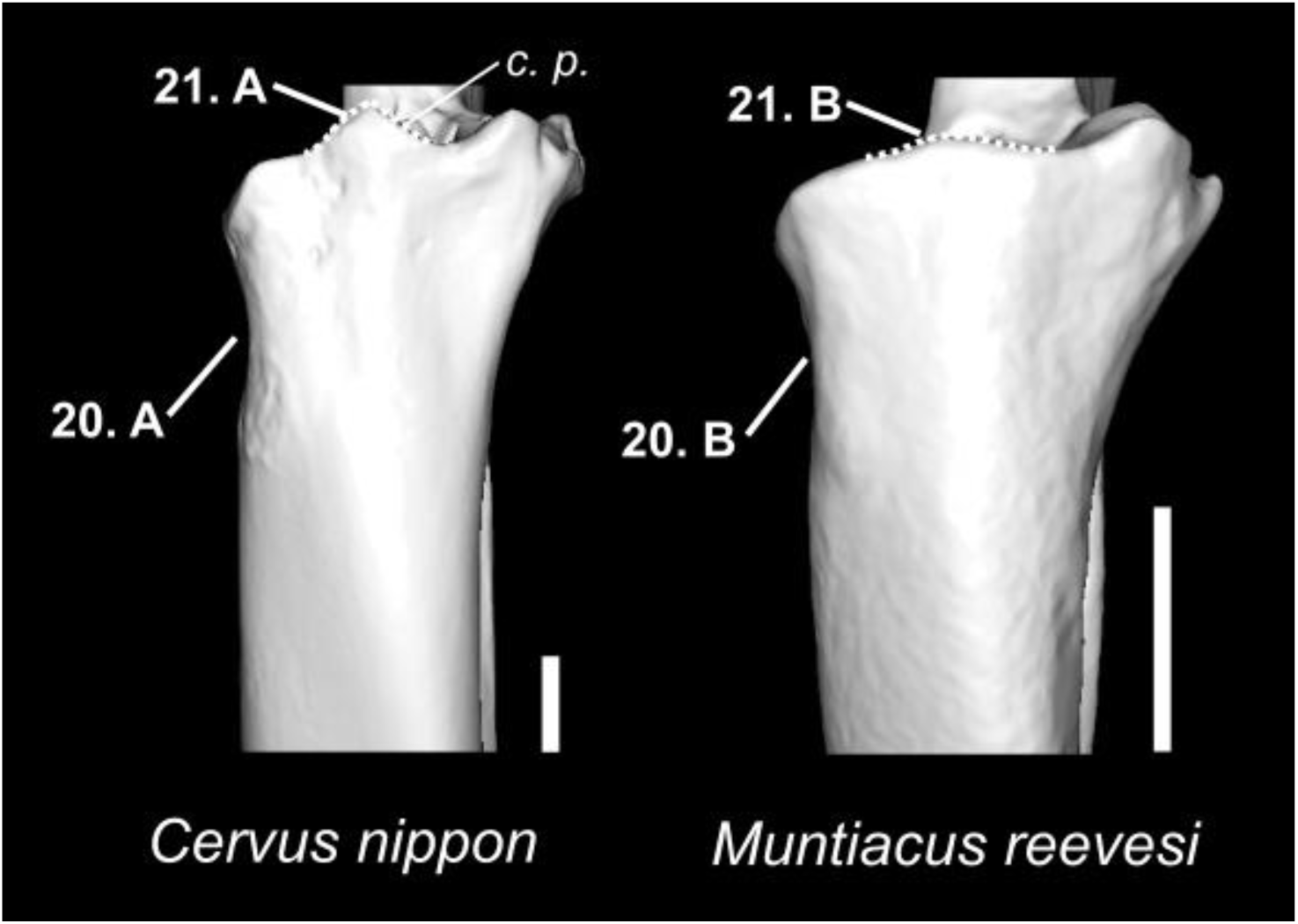
Character states visible on the anterior face of the left radius (characters 20 and 21). Abbreviations: c. p.: coronoid process. Scale bar = 1 cm.

**Figure 14.**
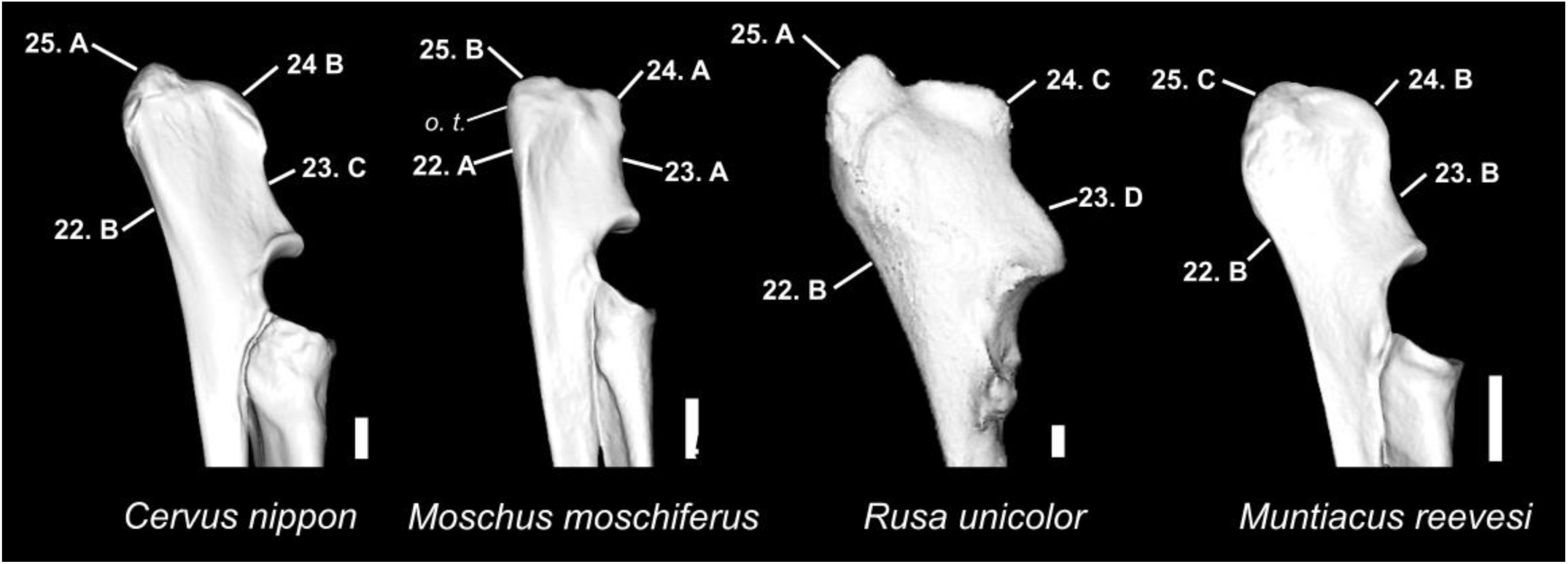
Character states visible on the medial face of the left ulna (characters 22, 23, 24, and 25). Abbreviations: o. t.: olecranon tuberosity. Scale bar = 1 cm.

**Figure 15.**
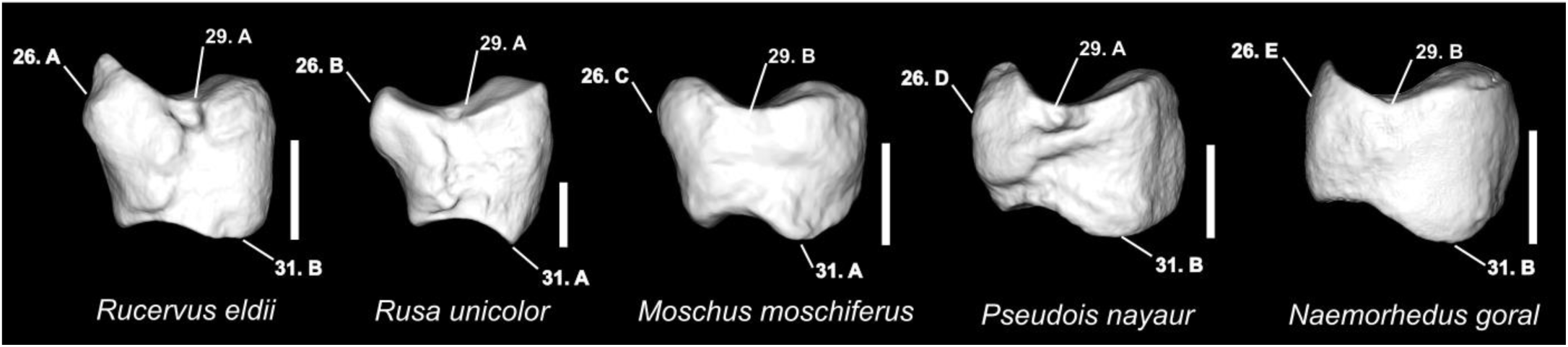
Character states visible on the lateral face of the left scaphoid (characters 26, 29, and 31). Scale bar = 0.5 cm.

**Figure 16.**
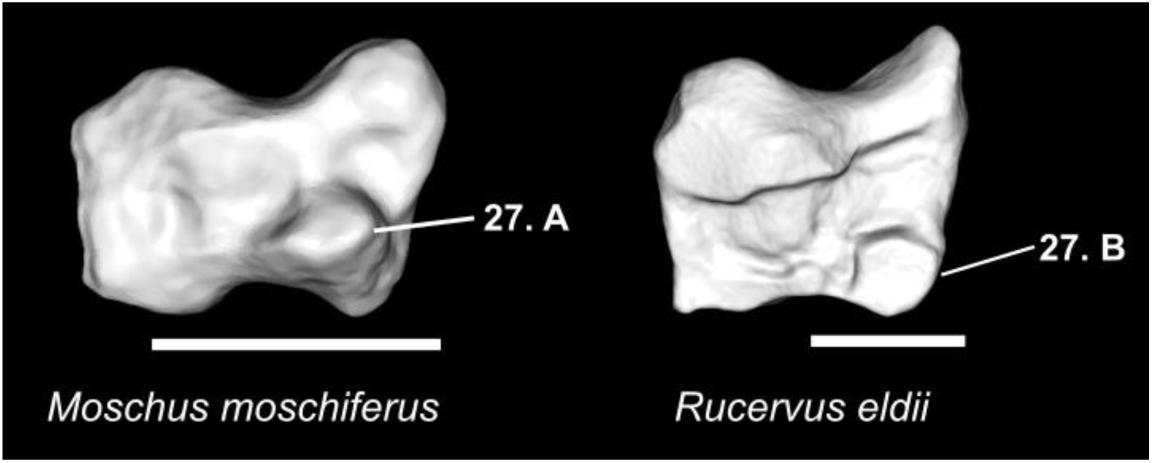
Character states visible on the medial face of the left scaphoid (character 27). Scale bar = 0.5 cm.

**Figure 17.**
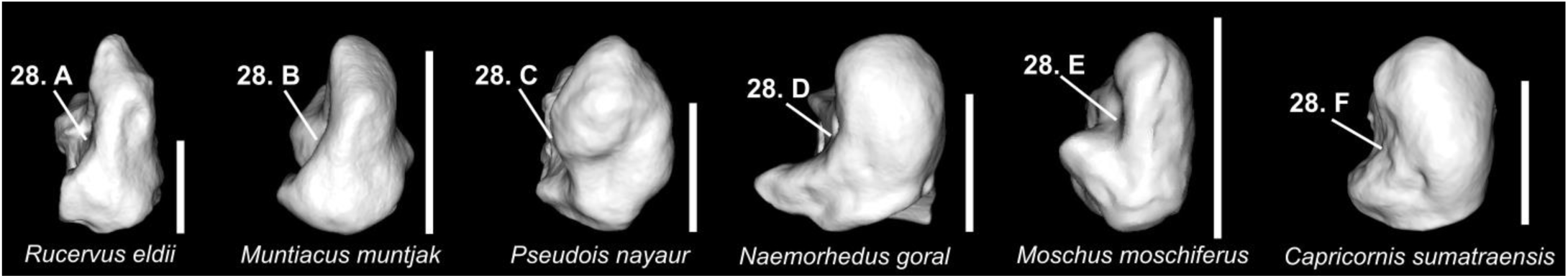
Character states visible on the posterior face of the left scaphoid (character 28). Scale bar = 1 cm.

**Figure 18.**
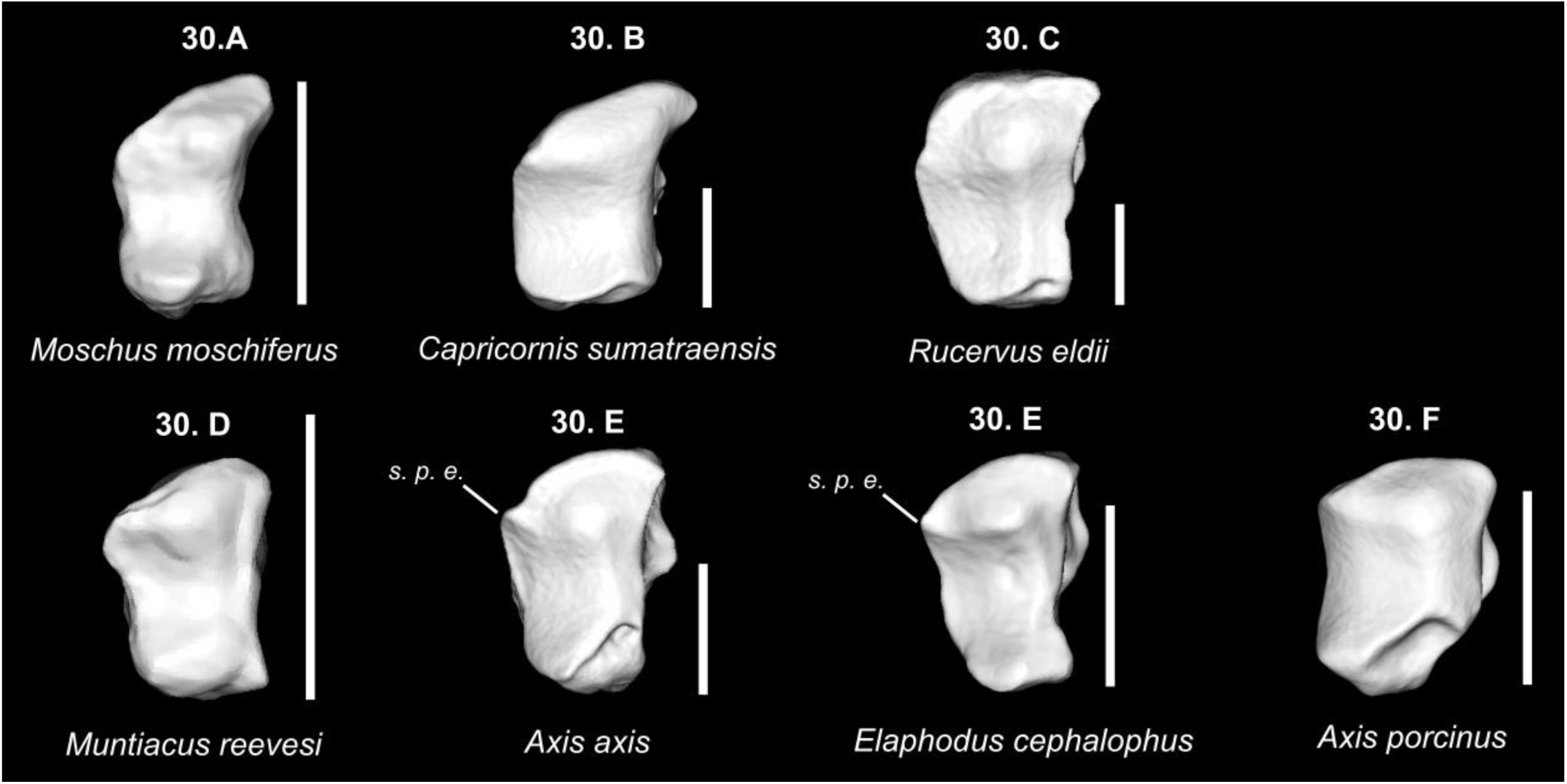
Character states visible on the ventral face of the left scaphoid (character 30). Scale bar = 1 cm.

**Figure 19.**
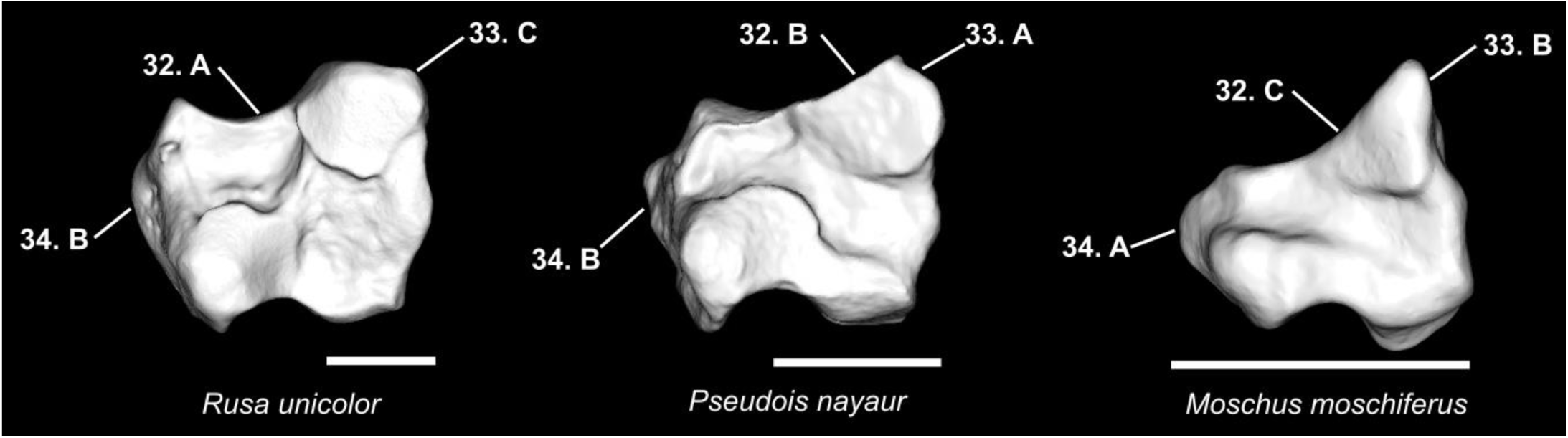
Character states visible on the lateral face of the right lunatum (characters 32, 33, and 34). Scale bar = 1 cm.

**Figure 20.**
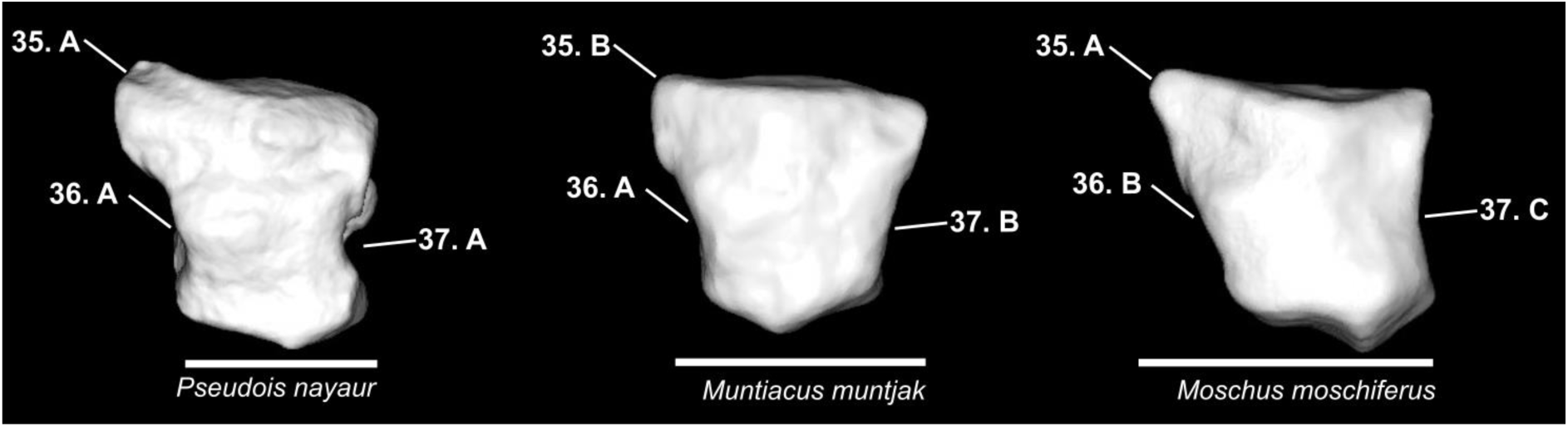
Character states visible on the anterior face of the right lunatum (characters 35, 36, and 37). Scale bar = 1 cm.

**Figure 21.**
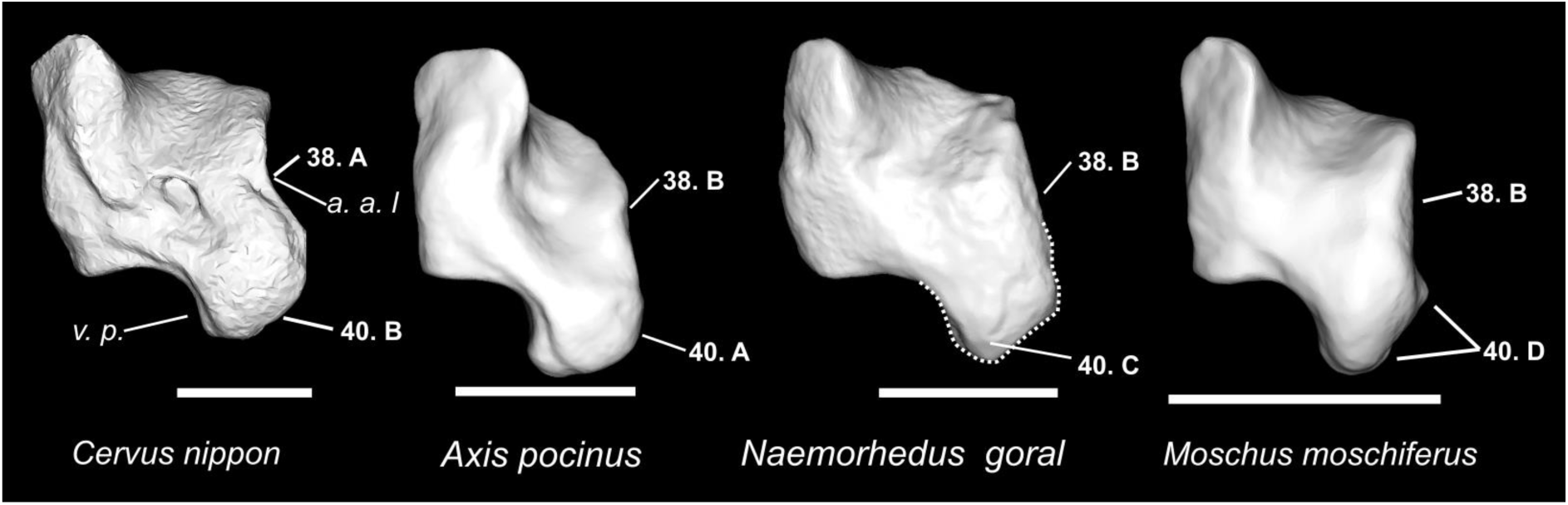
Character states visible on the lateral face of the left pyramidal. (Characters 38 and 40). Abbreviations: a. a. l.: anterior articular surface with the lunatum; v. p.: ventral process. Scale bar = 1 cm.

**Figure 22.**
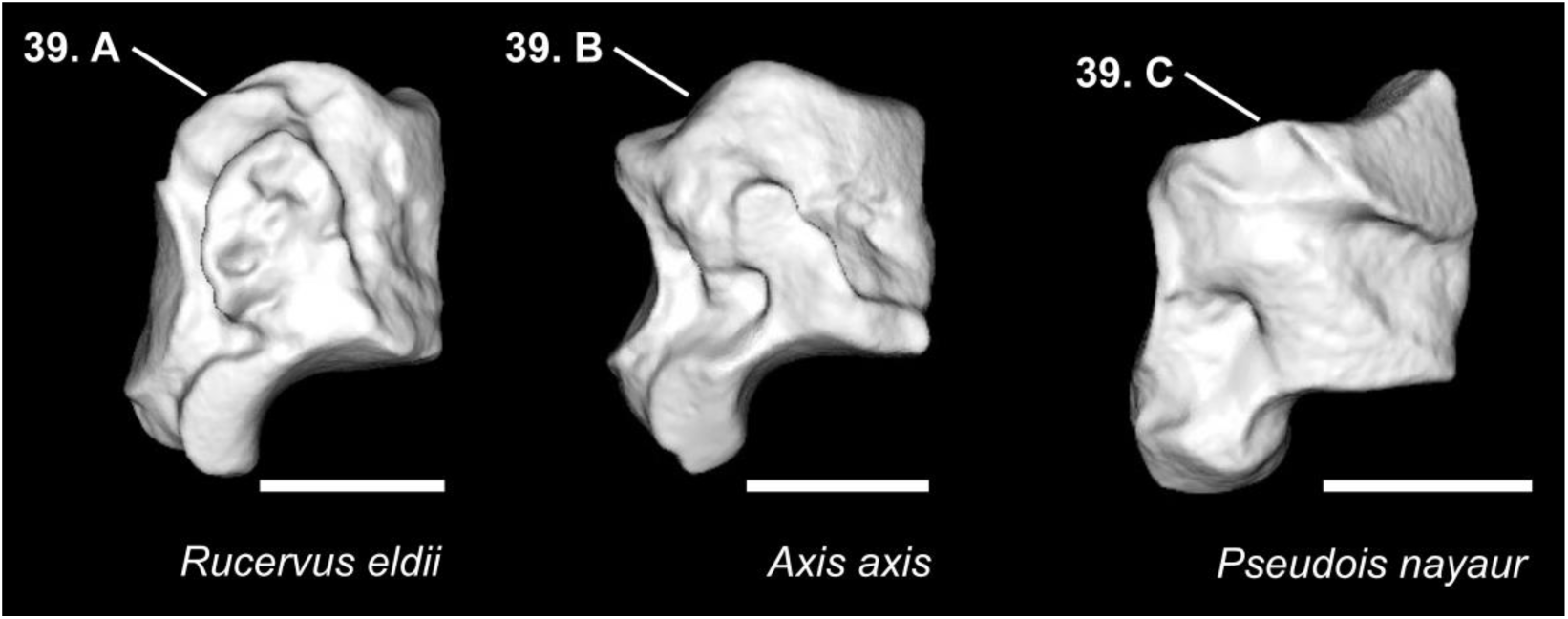
Character states visible on the medial face of the left pyramidal. (Character 39). Abbreviations: m. a. s.: medial articular surface. Scale bar = 1 cm.

**Figure 23.**
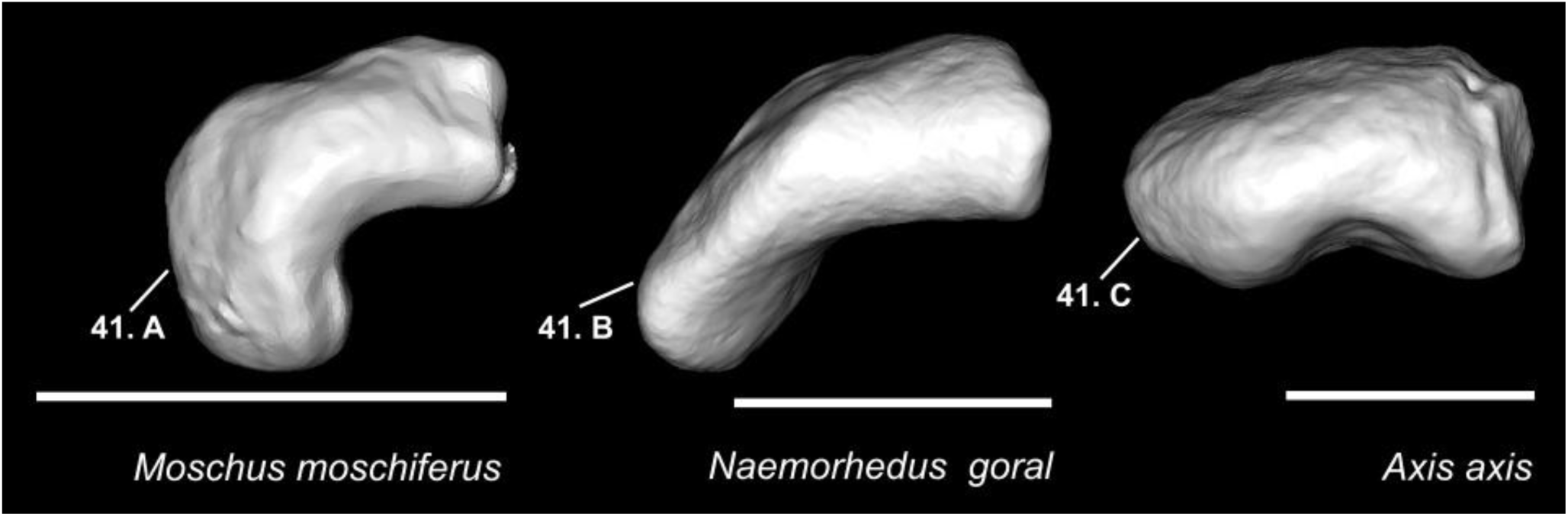
Character states visible on the ventral face of the left pisiform (Character 41). Scale bar = 1 cm.

**Figure 24.**
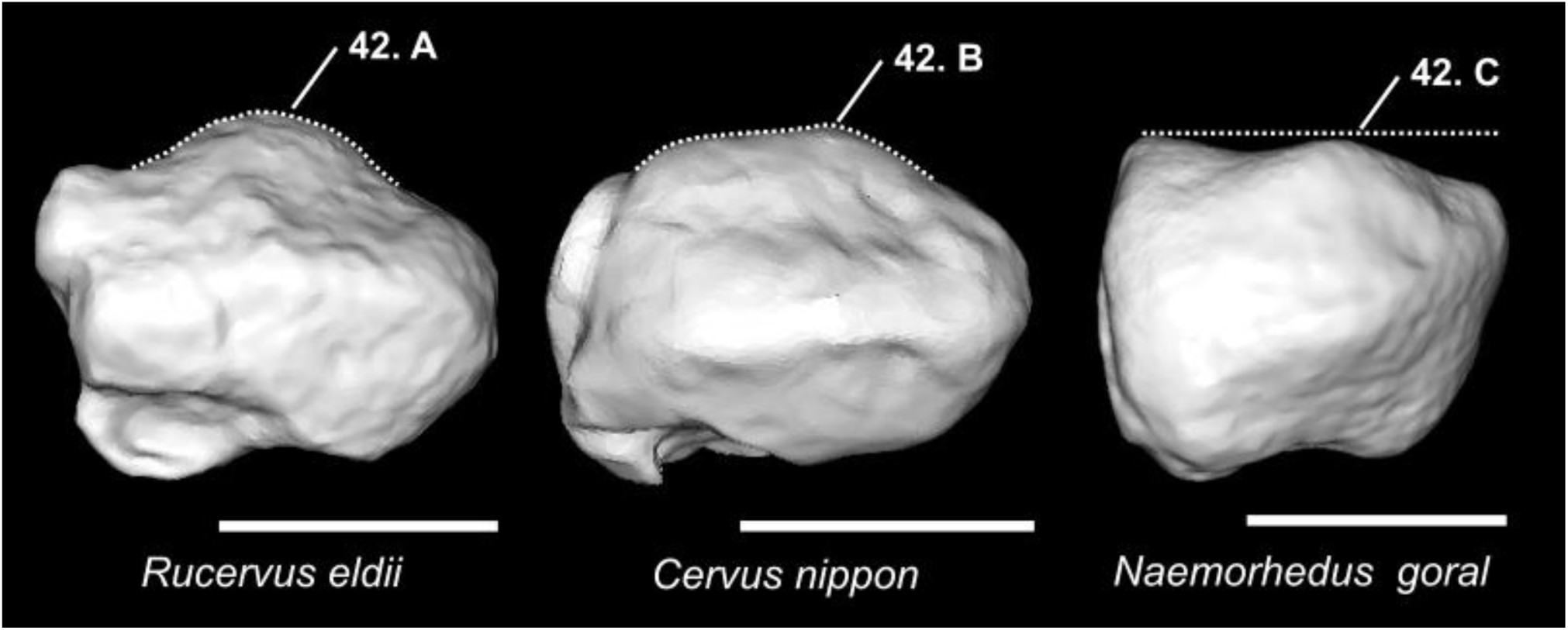
Character states visible on the lateral face of the left pisiform (character 42). Scale bar = 1 cm.

**Figure 25.**
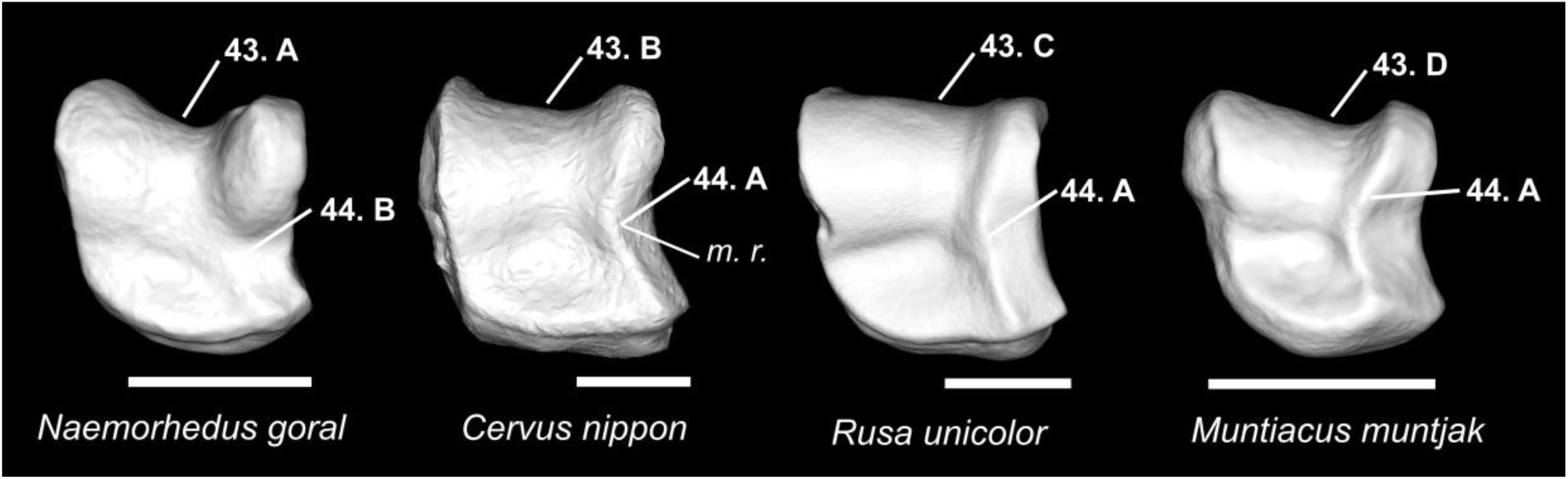
Character states visible on the dorsal face of the left magnum (characters 43 and 44). Abbreviations: m. r.: medial ridge. Scale bar = 1 cm.

**Figure 26.**
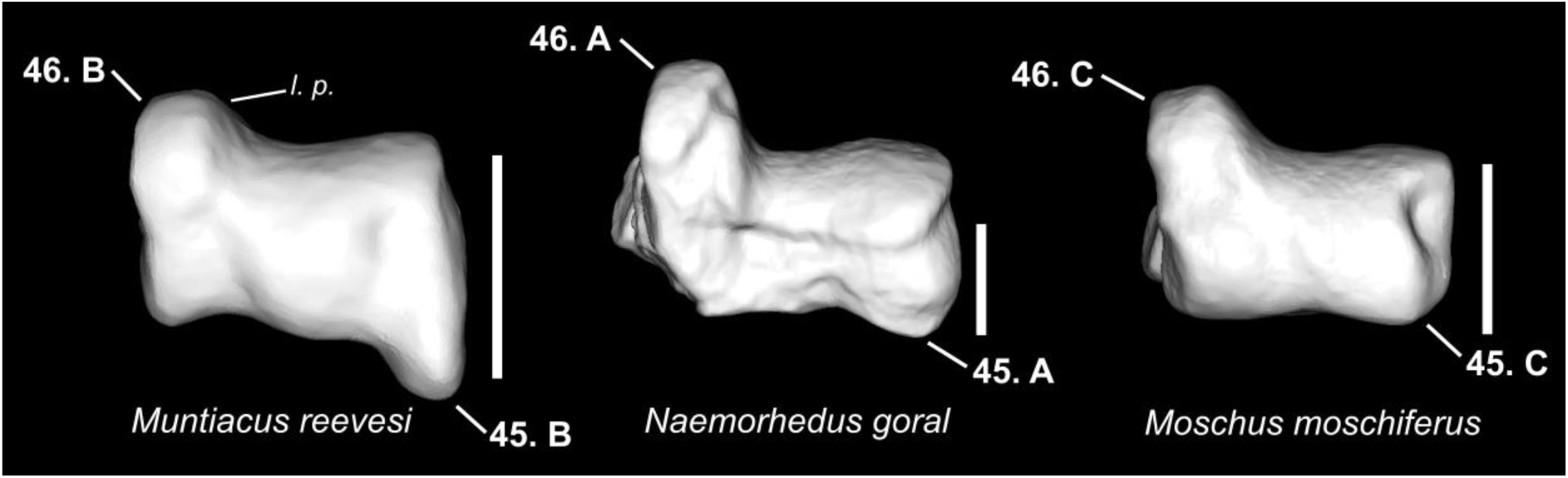
Character states visible on the posterior face of the left magnum (characters 45 and 46). Abbreviations: l. p.: lateroventral projection. Scale bar = 0.5 cm.

**Figure 27.**
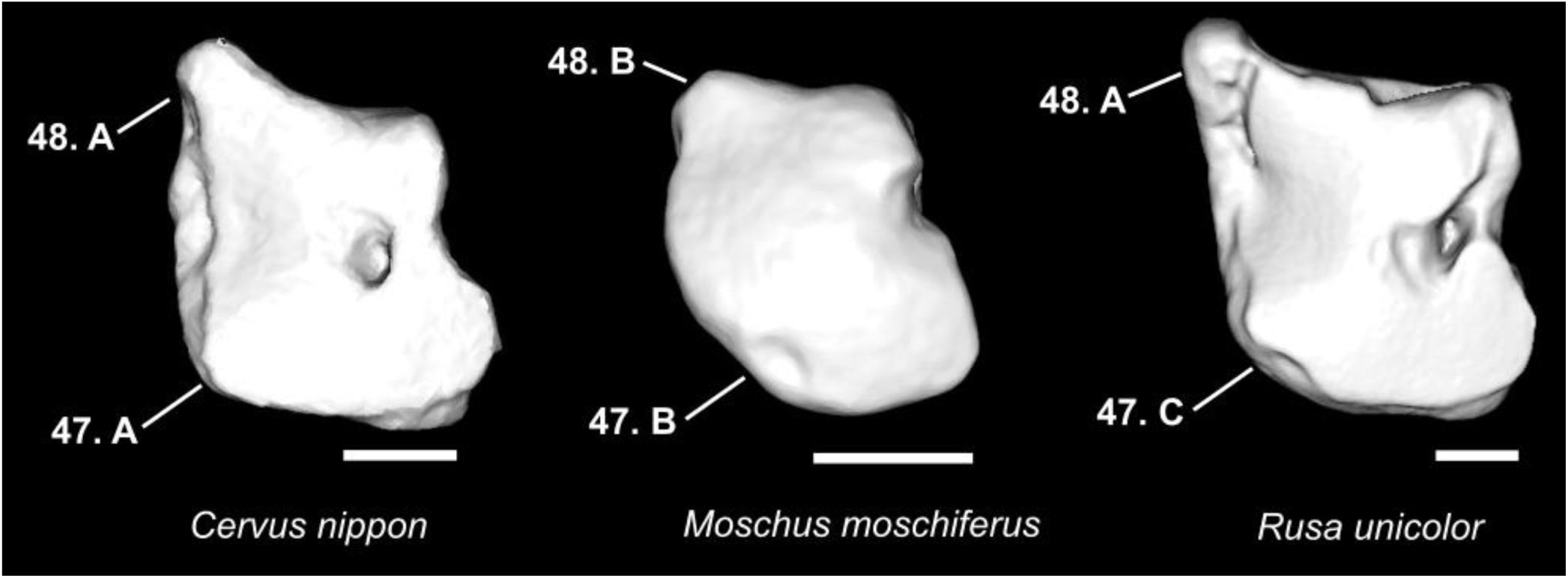
Character states visible on the ventral face of the right magnum (characters 47 and 48). Scale bar = 0.5 cm.

**Figure 28.**
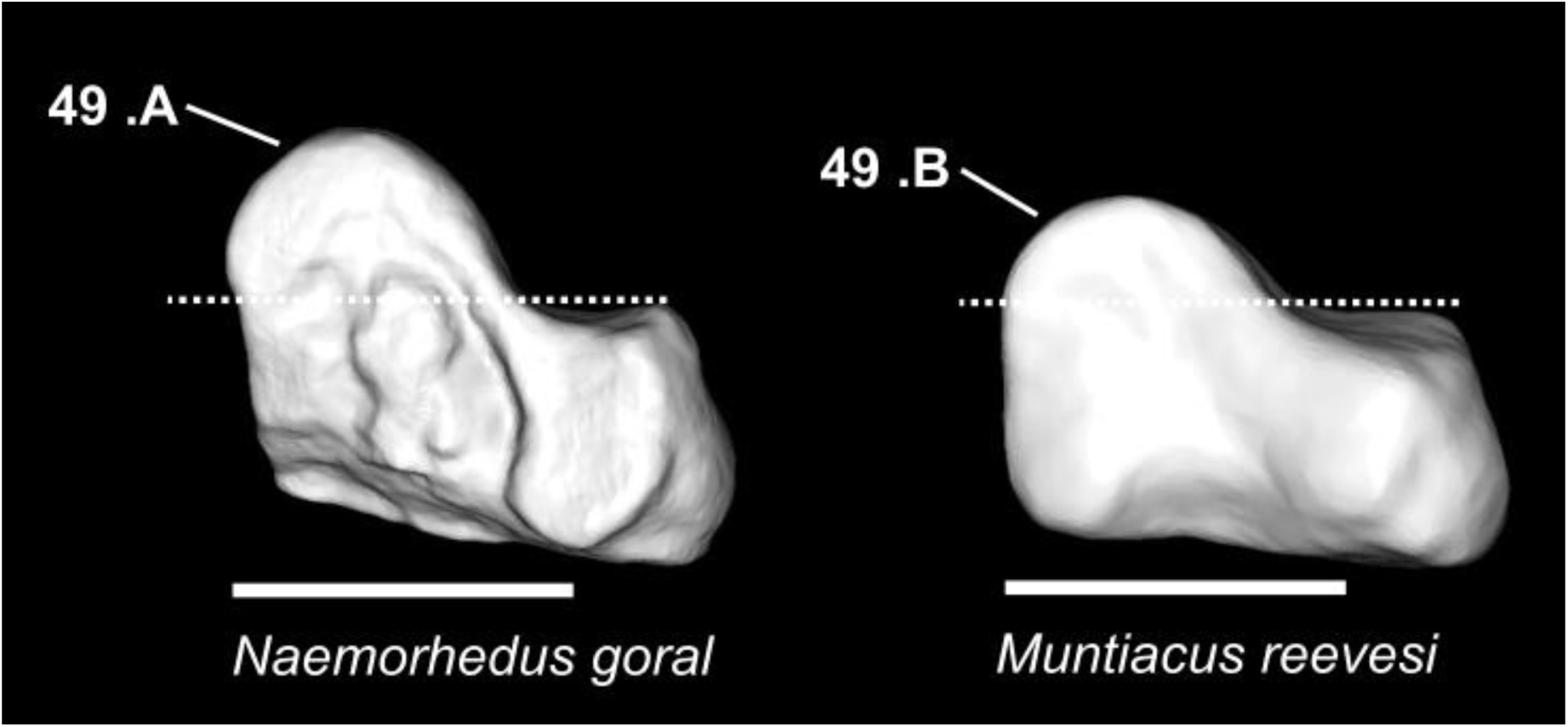
Character states visible on the medial face of the right magnum (character 49). Scale bar = 0.5 cm.

**Figure 29.**
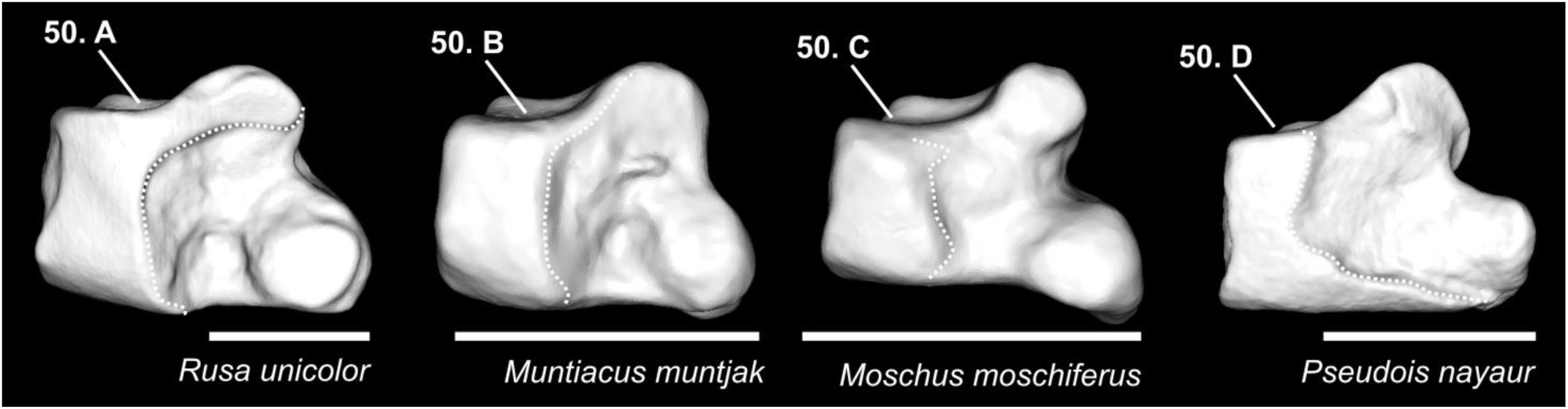
Character states visible on the medial face of the left unciform (character 50). Scale bar = 1 cm.

**Figure 30.**
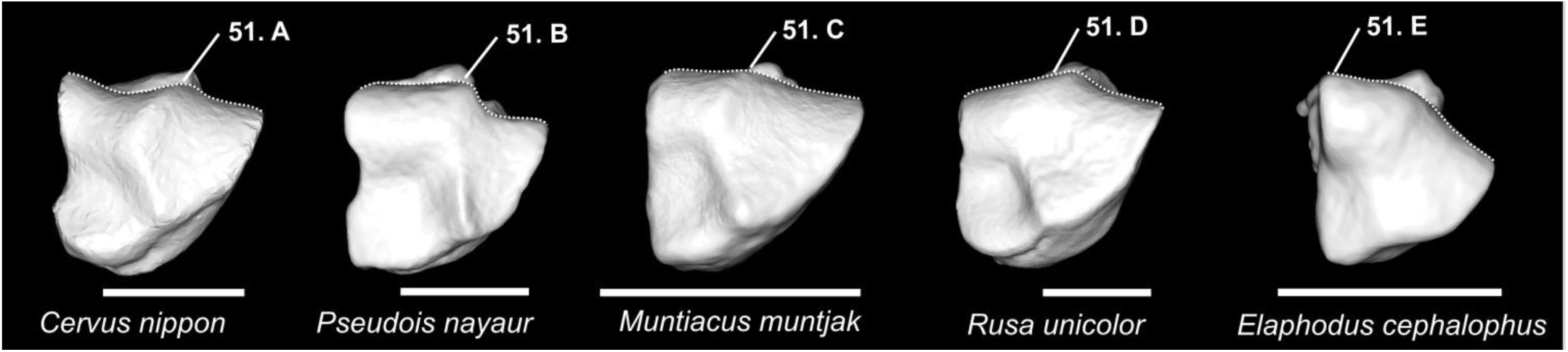
Character states visible on the dorsal face of the left unciform (character 51). Scale bar = 1 cm.

**Figure 31.**
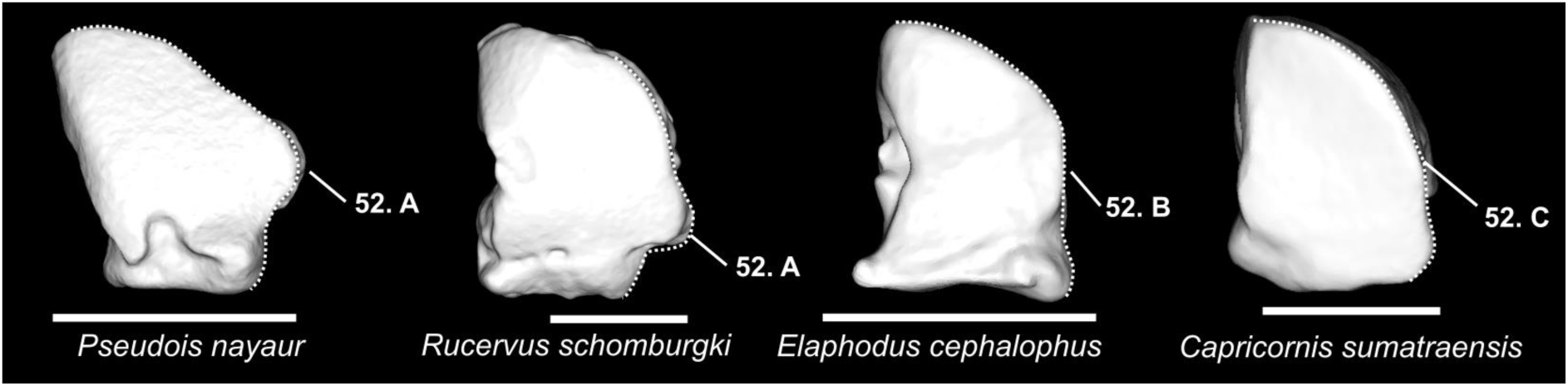
Character states visible on the ventral face of the left unciform (character 52). Scale bar = 1 cm.

**Figure 32.**
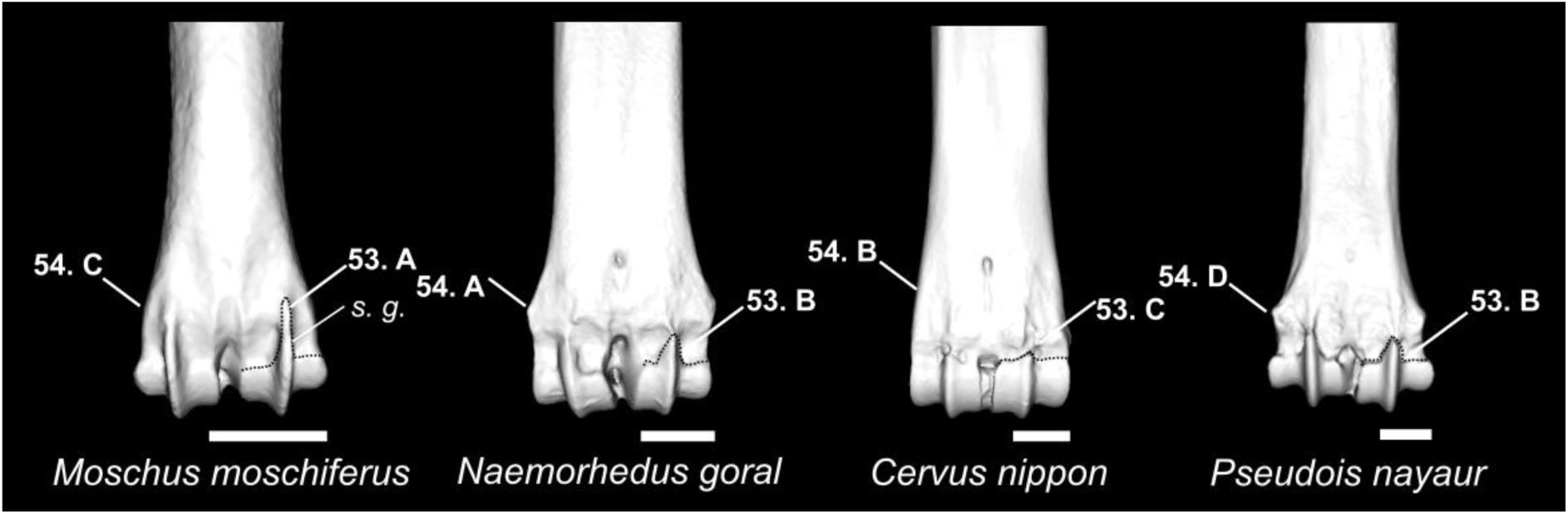
Character states visible on the posterior face of the distal part of the right metacarpus (Characters 53 and 54). Abbreviations: s. r.: sagittal ridge. Scale bar = 1 cm.

**Figure 33.**
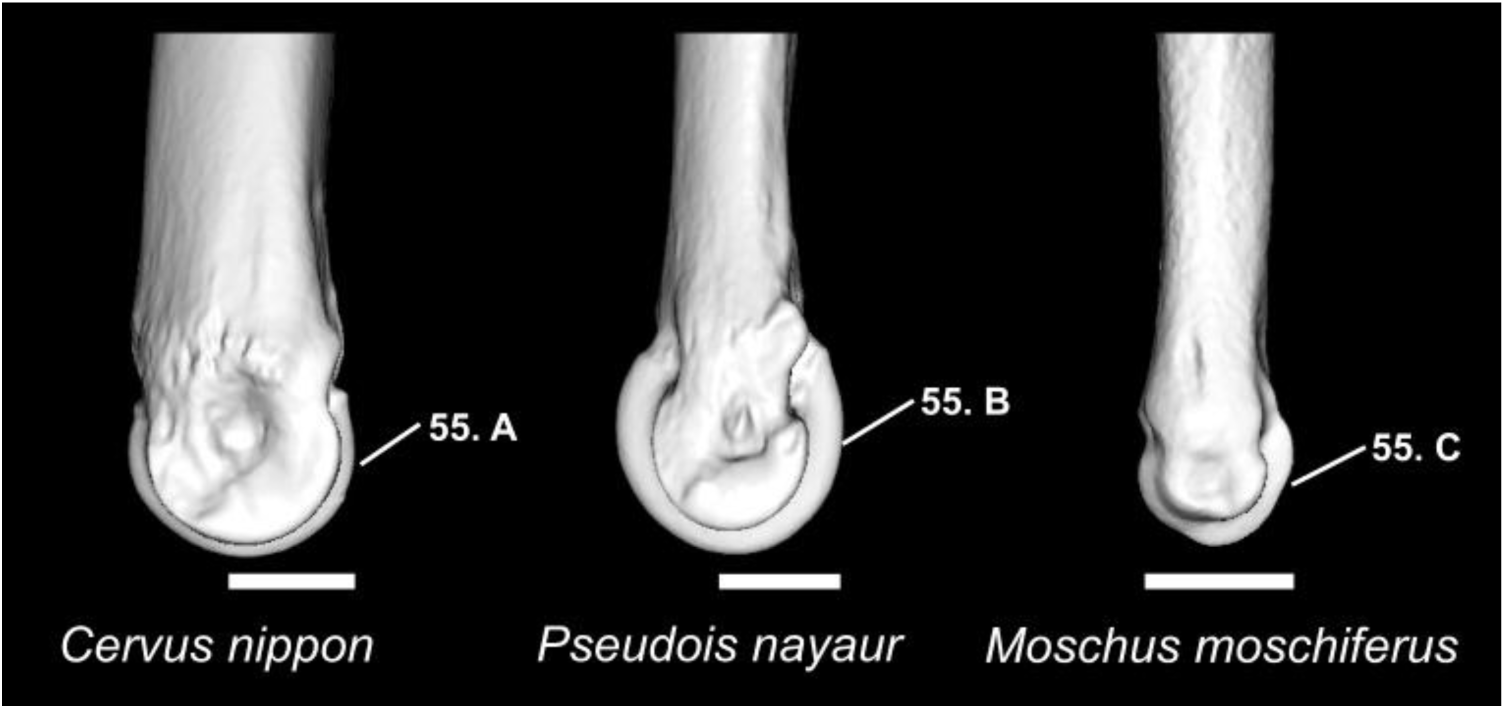
Character states visible on the medial face of the distal extremity of the right metacarpus (character 55). Abbreviations: m. c.: medial condyle. Scale bar = 1 cm.

**Figure 34.**
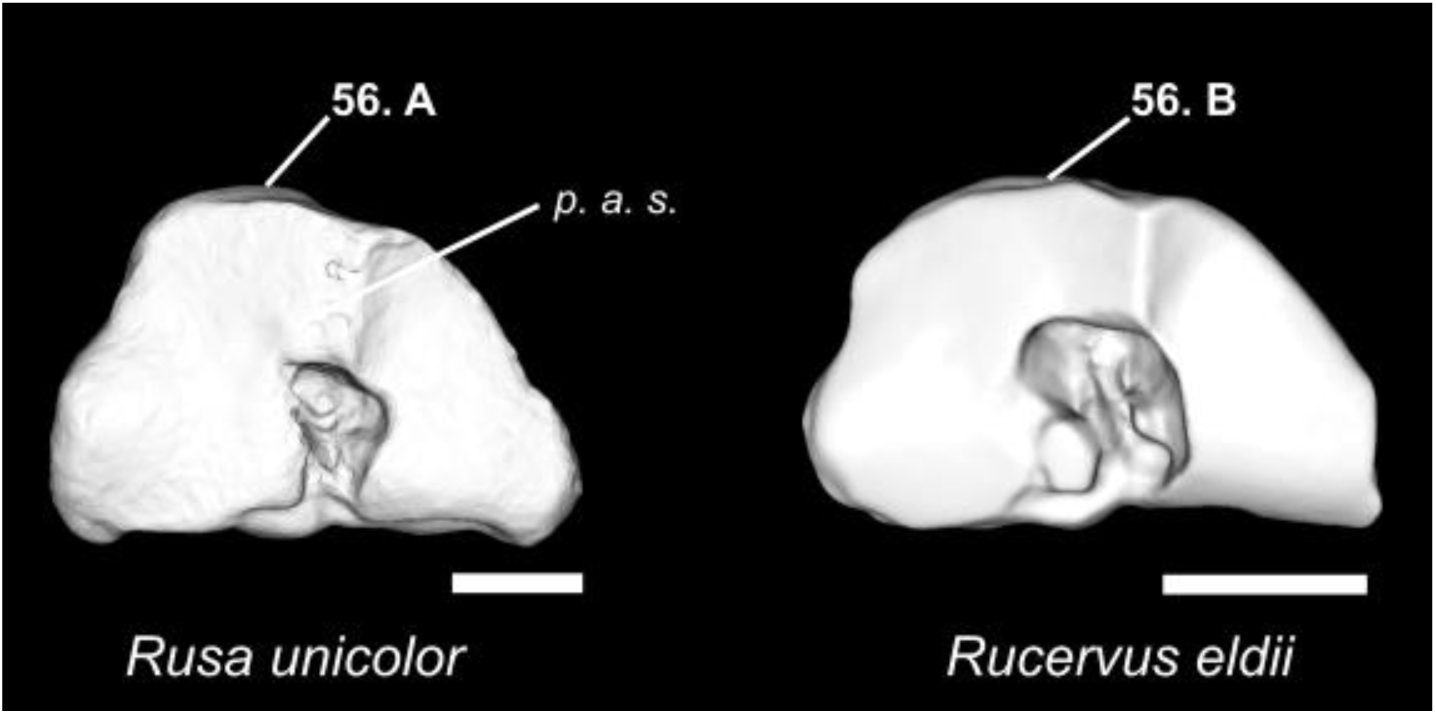
Character states visible on the dorsal face of the right metacarpus (character 56). Abbreviations: p. a. s.: proximal articular surface. Scale bar = 1 cm.

**Figure 35.**
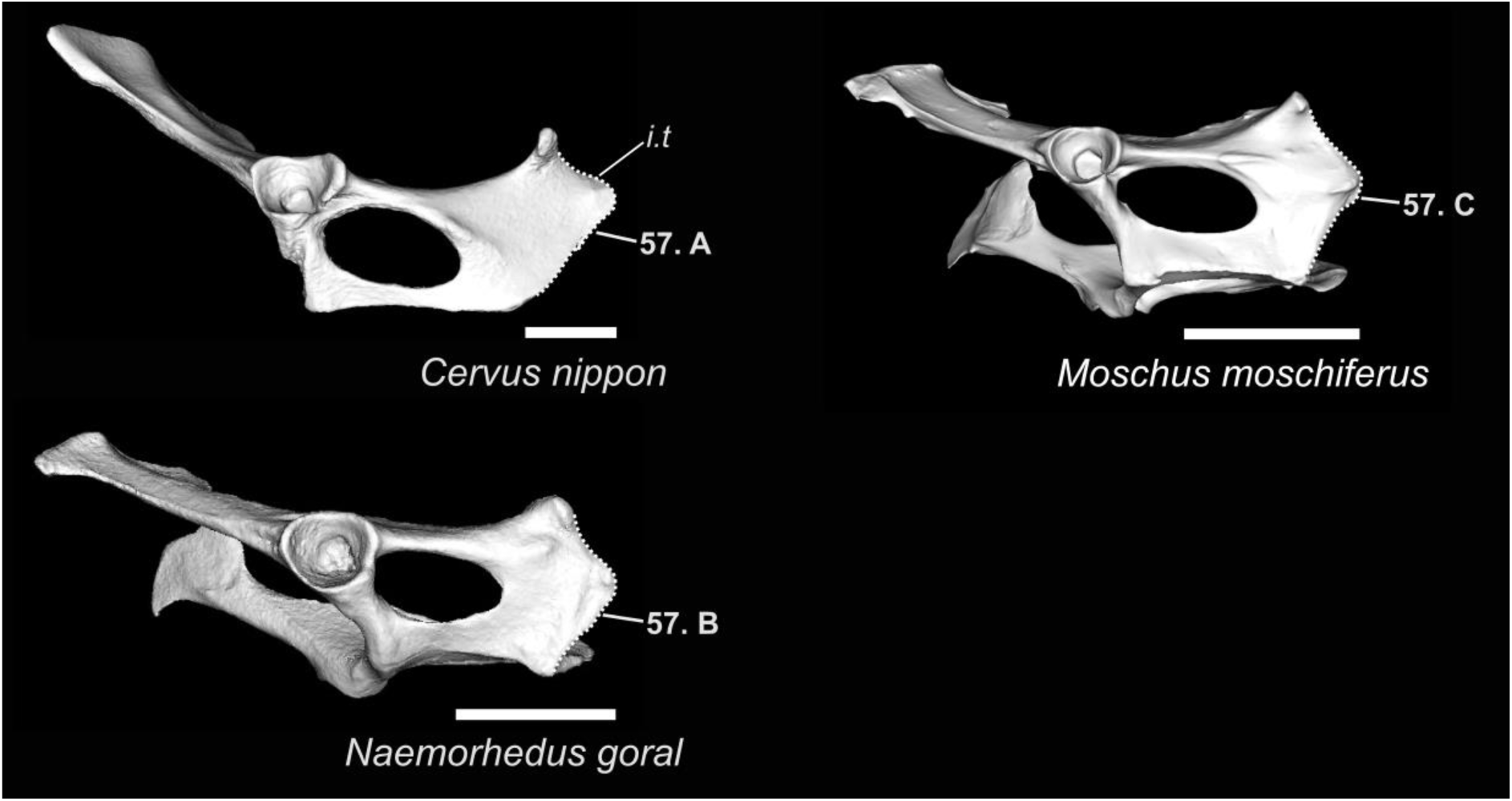
Character states visible on the lateral face of the left coxal (character 57). Abbreviations: i.t.: ischiatic tuberosity. Scale bar = 5 cm.

**Figure 36.**
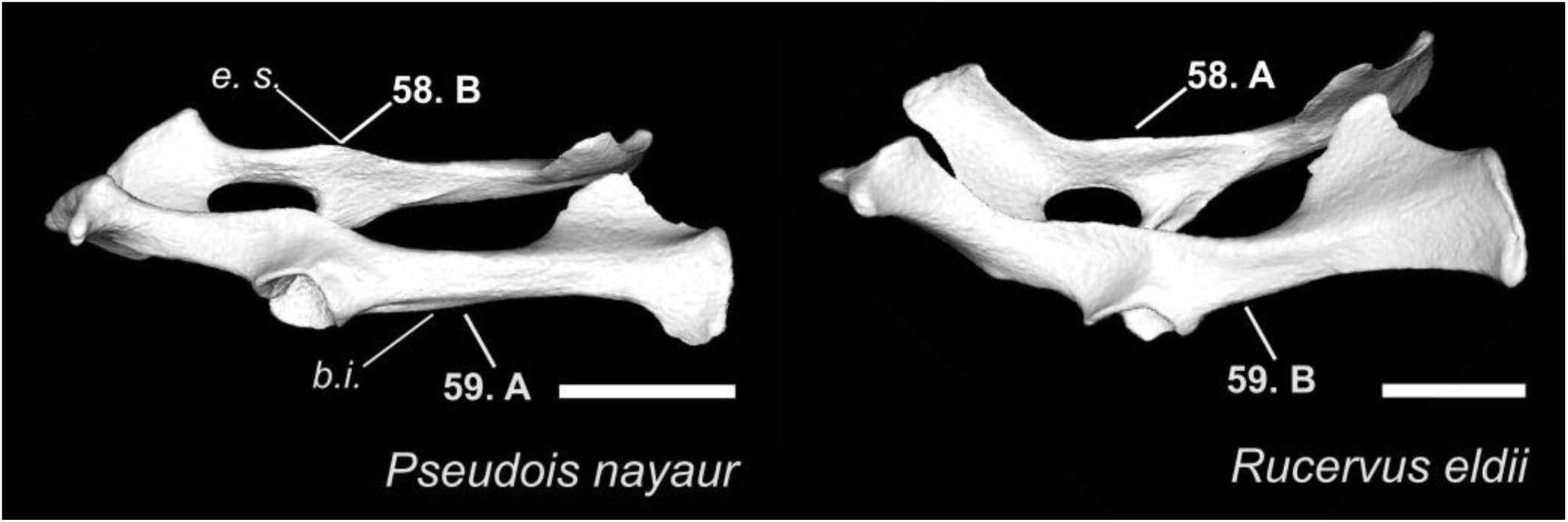
Character states visible on the lateral face of the coxal bones (characters 58 and 59). Abbreviations: s. s.: sciatic spine, b.i.: body of ilium. Scale bar = 5 cm.

**Figure 37.**
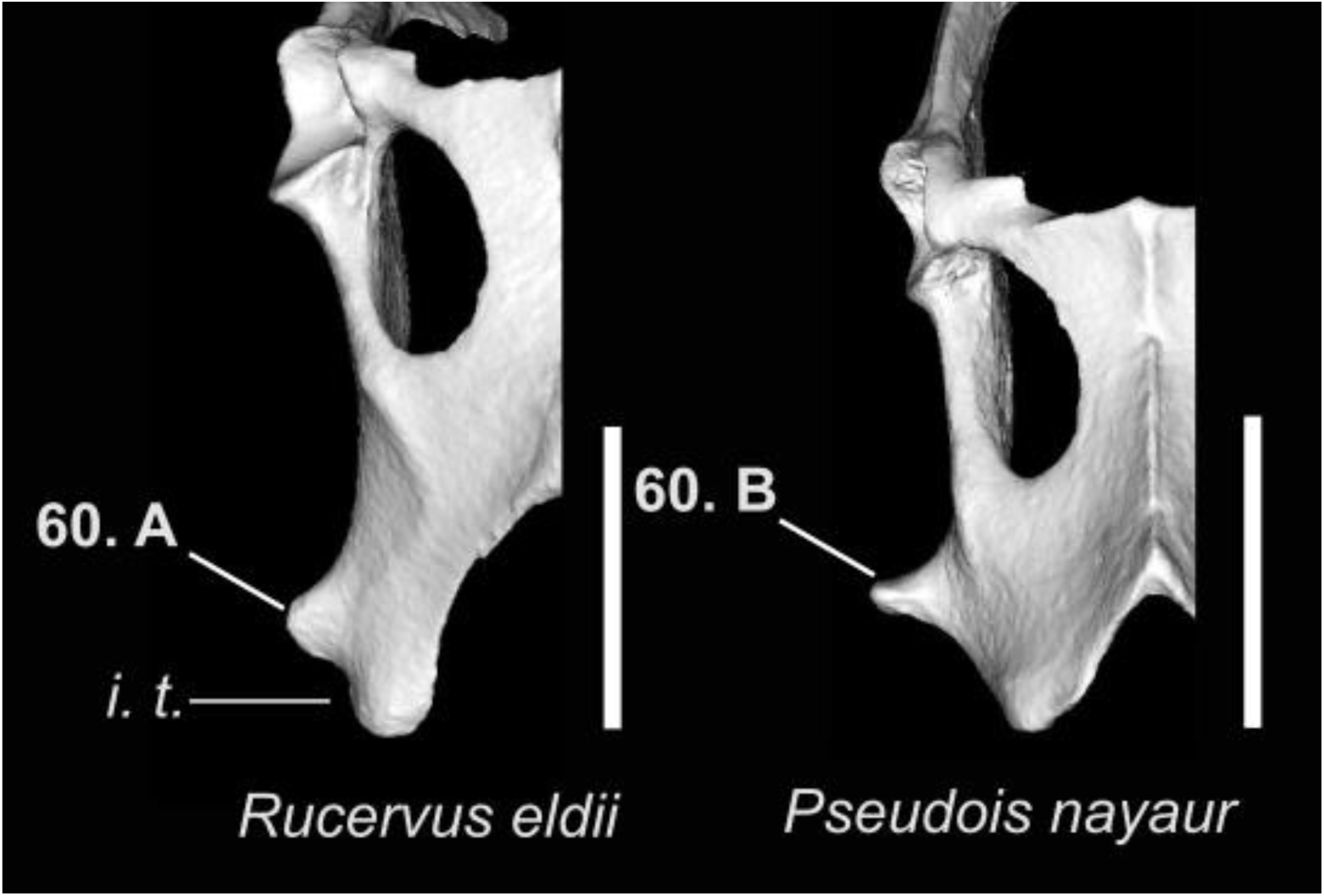
Character states visible on the ventral face of the coxal bones (character 60). Abbreviation: i. t.: ischiatic tuberosity. Scale bar = 5 cm.

**Figure 38.**
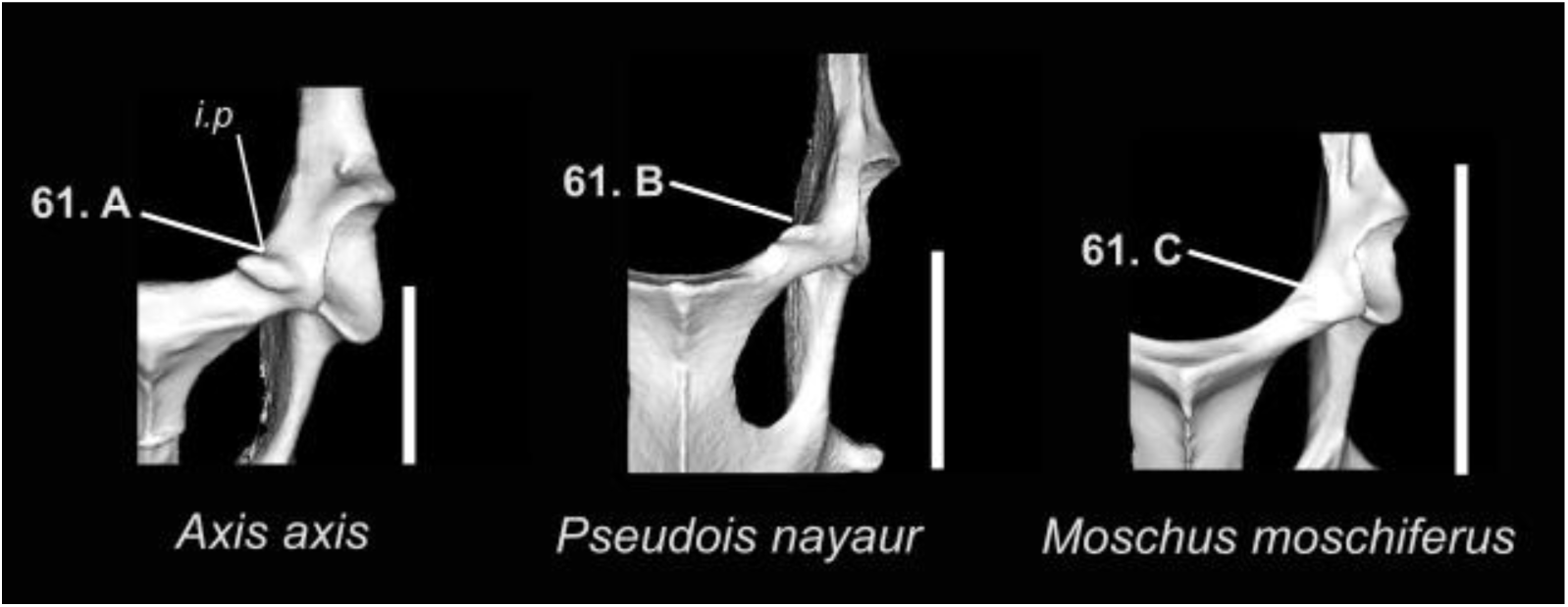
Character states visible on the ventral face of the coxal (character 61). Abbreviation: i. p.: iliopubic eminence. Scale bar = 5 cm.

**Figure 39.**
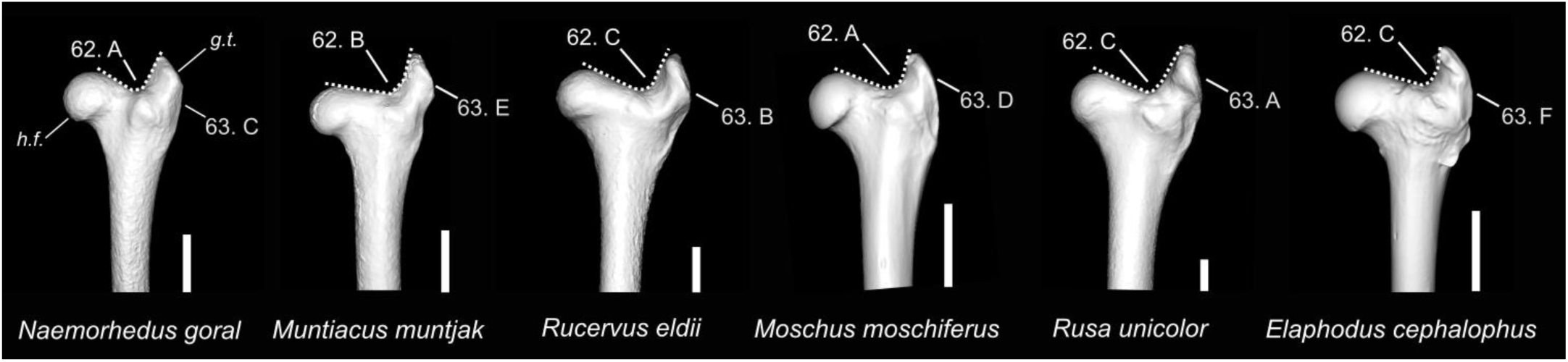
Character states visible on the anterior face of the left femur (characters 62 and 63). Abbreviations: g. t.: greater trochanter; h. f.: head of the femur. Scale bar = 2 cm.

**Figure 40.**
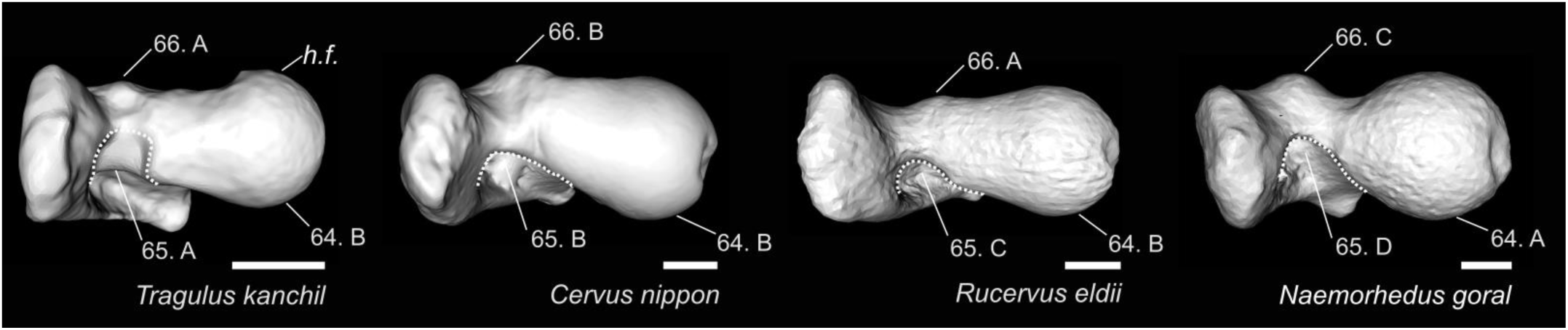
Character states visible on the dorsal face of the left femur (characters 64, 65, and 66). *Tragulus kanchil* is figured here because none of the specimens scanned for the species included in this study displayed these character states. Abbreviation: h. f.: head of the femur. Scale bar = 1 cm.

**Figure 41.**
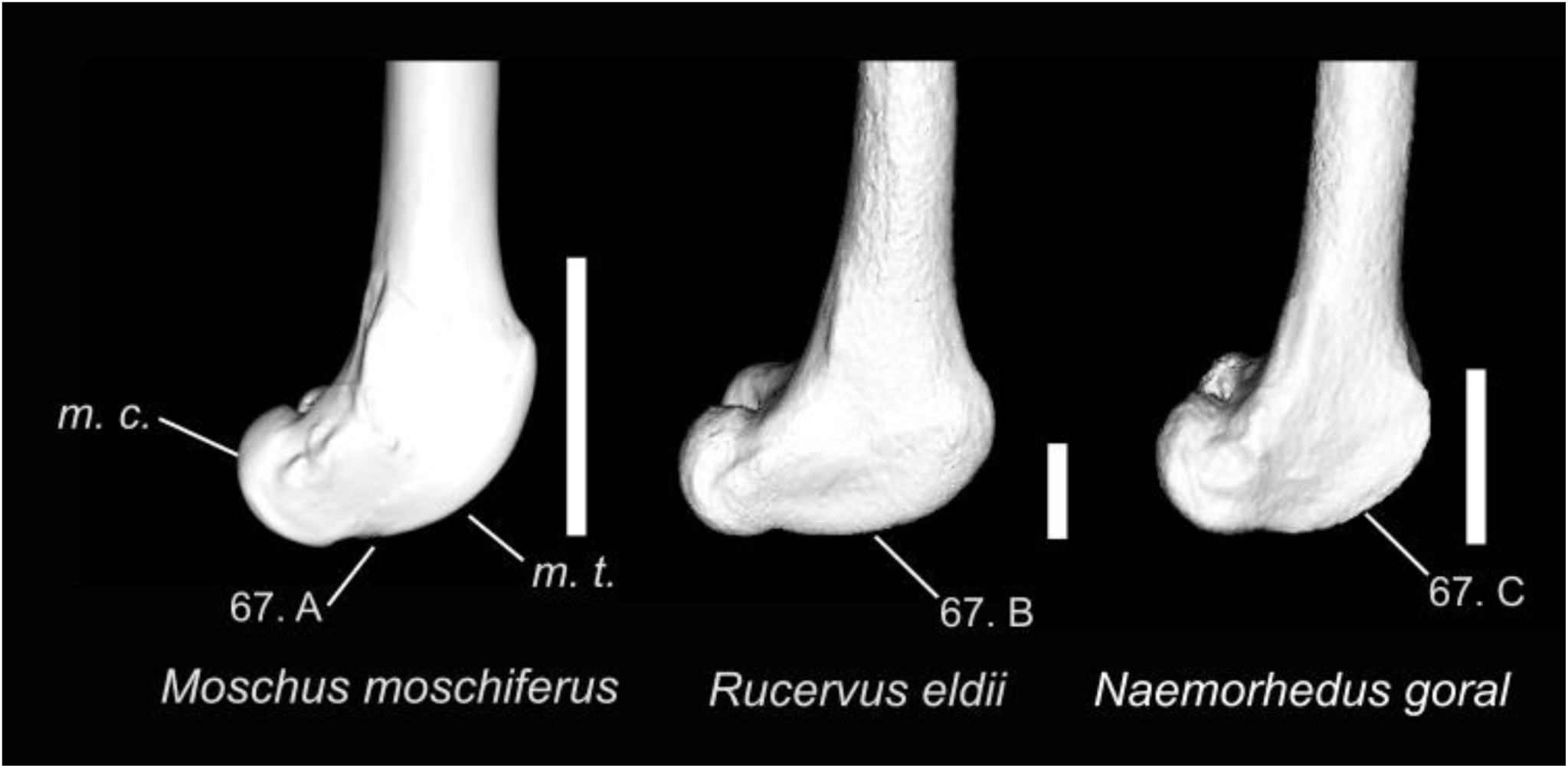
Character states visible on the medial face of the femur (character 67). Abbreviations: m. t.: medial lip of the trochlea; m. c.: medial condyle. Scale bar = 2 cm.

**Figure 42.**
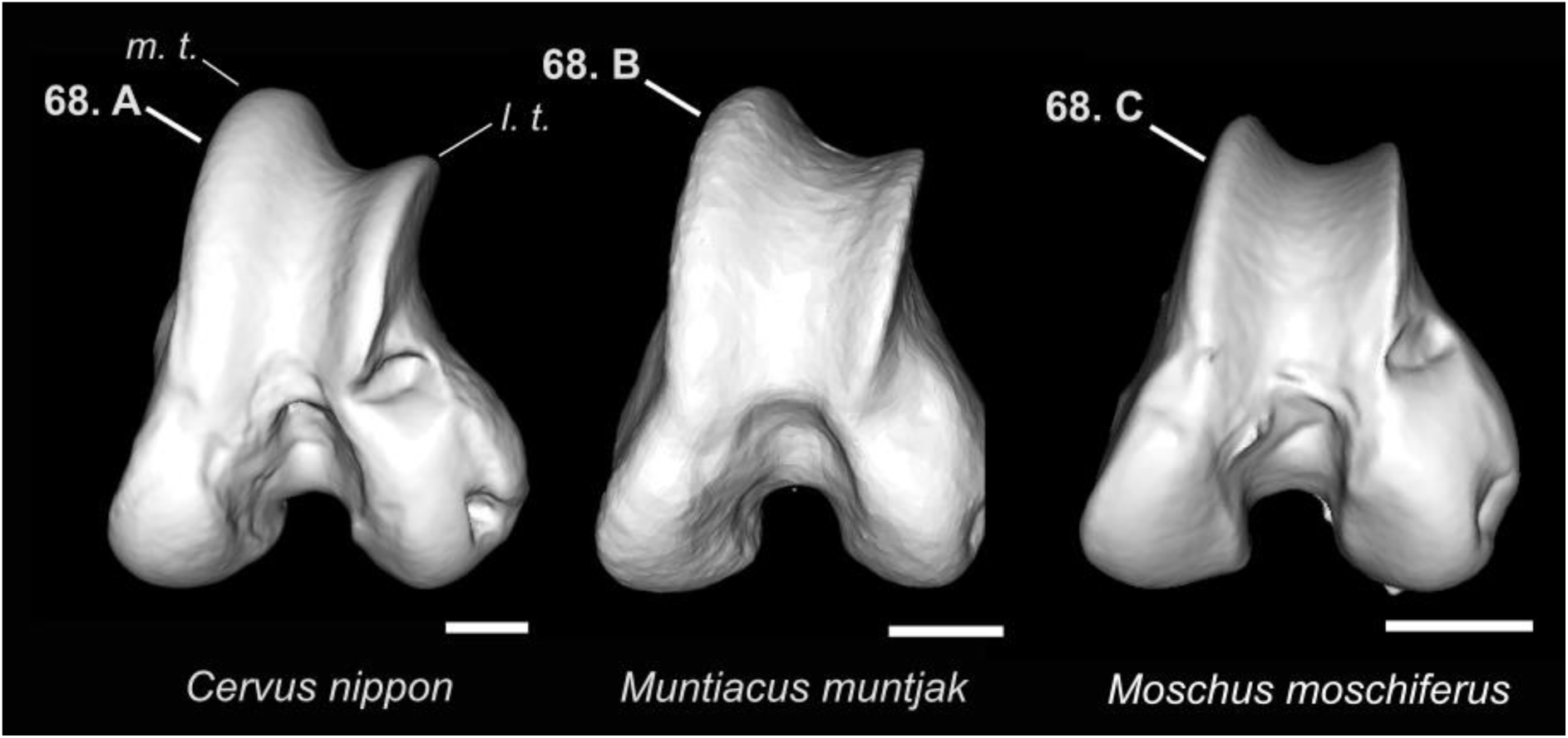
Character states visible on the ventral face of the left femur (character 68). Abbreviations: m. t.: medial lip of the trochlea; l. t.: lateral lip of the trochlea. Scale bar = 1 cm.

**Figure 43.**
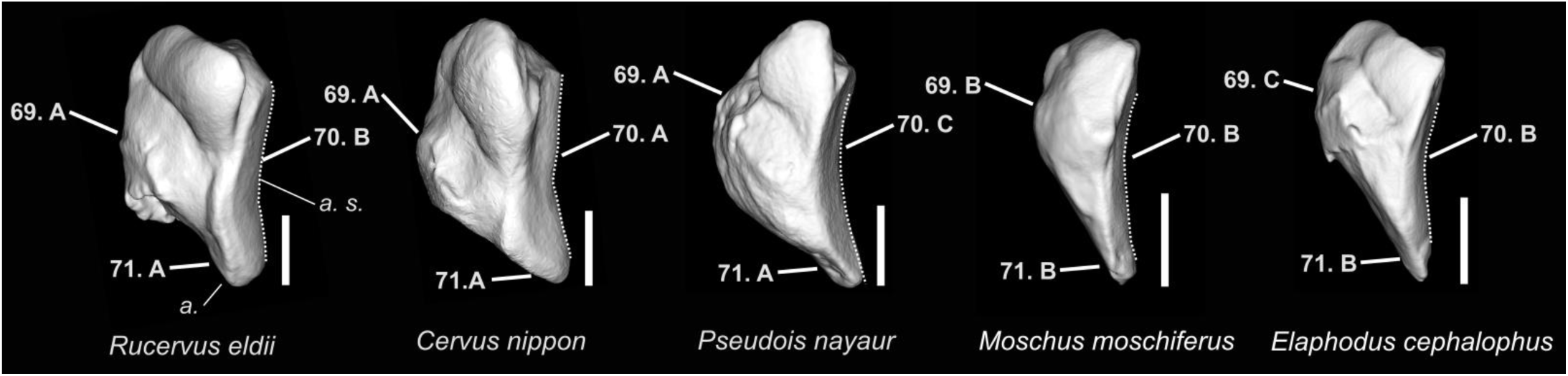
Character states visible on the medial face of the left patella (characters 69, 70, and 71). Abbreviations: a.: apex; a. s.: articular surface. Scale bar = 1 cm.

**Figure 44.**
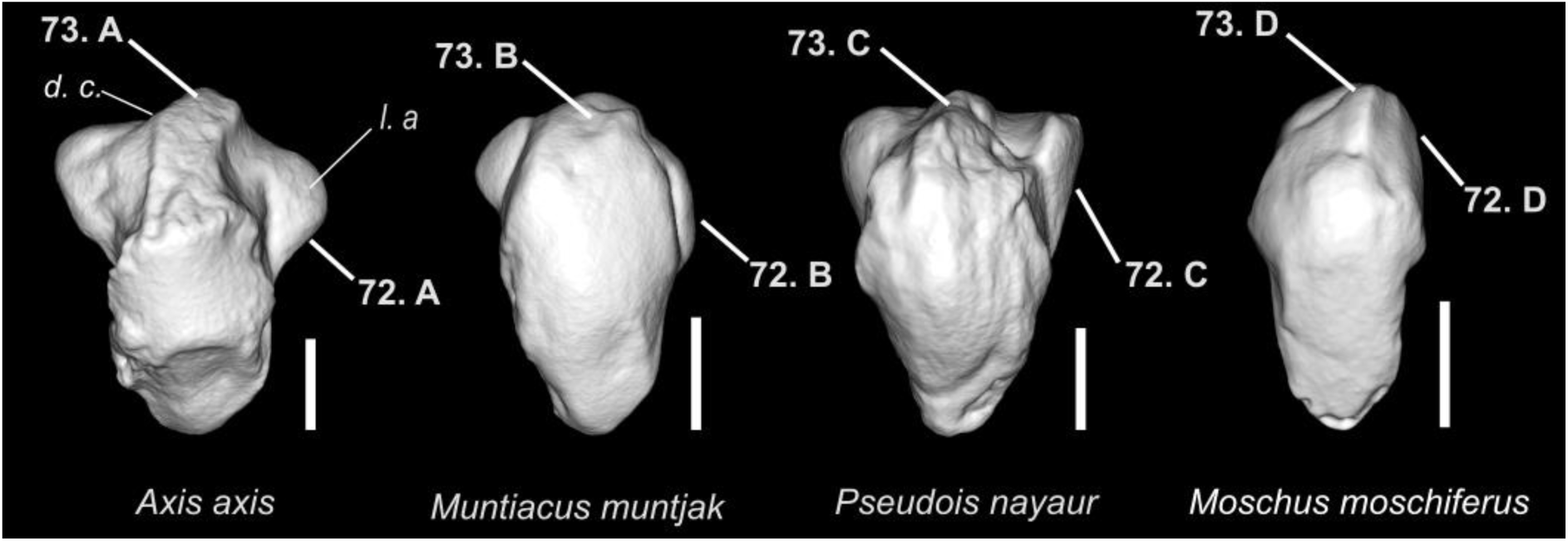
Character states visible on the anterior face of the left patella (characters 72 and 73). Abbreviations: m. a.: medial angle; d. c.: dorsal crest. Scale bar = 1 cm.

**Figure 45.**
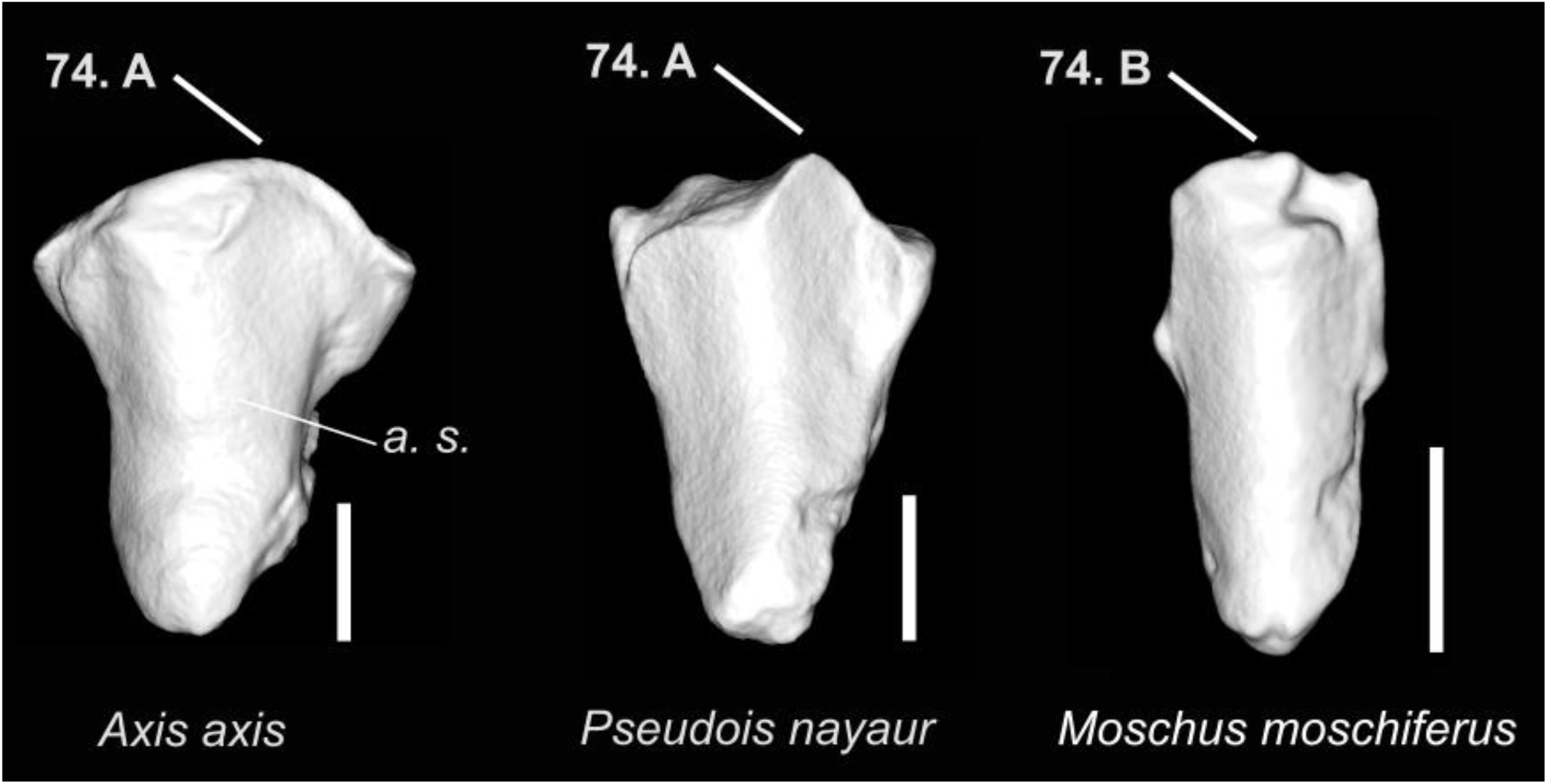
Character states visible on the posterior face of the left patella (character 74). Abbreviations: a. s.: articular surface. Scale bar = 1 cm.

**Figure 46.**
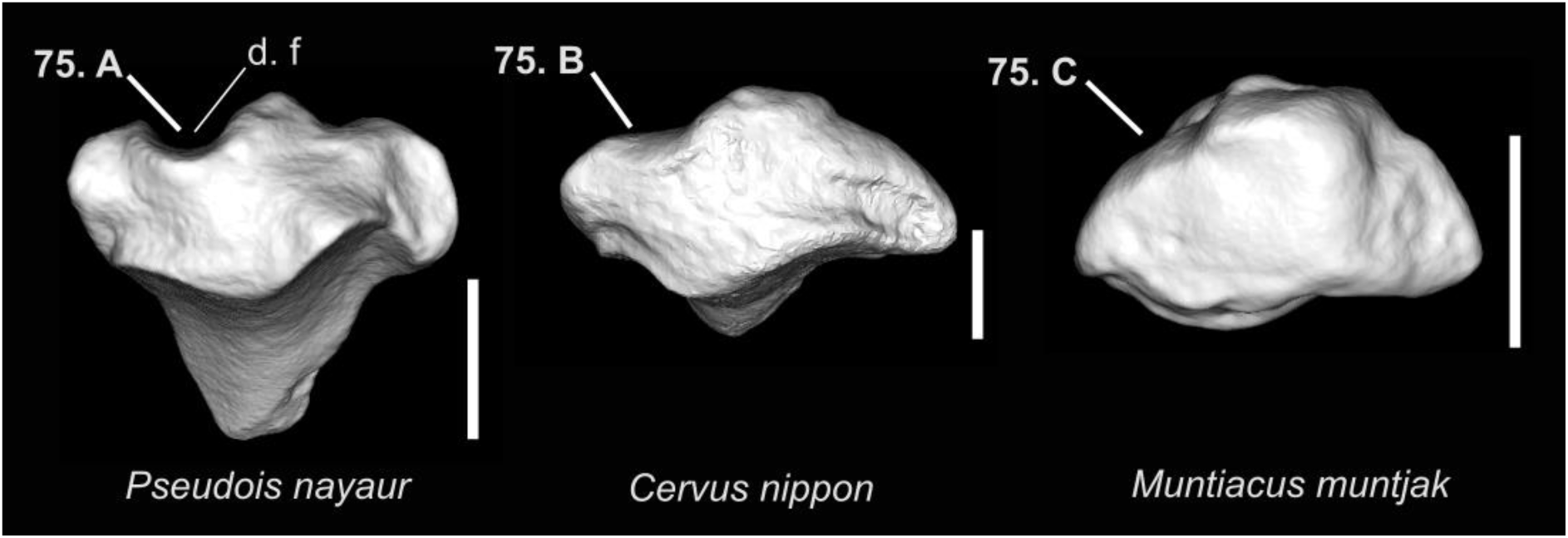
Character states visible on the dorsal face of the left patella (character 75). Abbreviations: d. f.: dorsomedial fossa. Scale bar = 1 cm.

**Figure 47.**
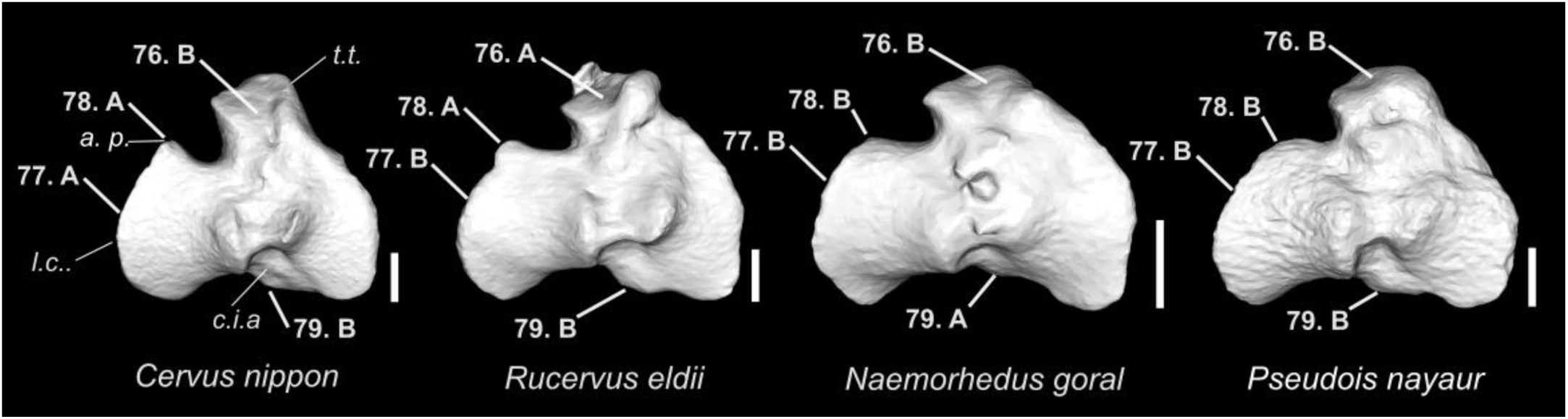
Character states visible on the dorsal face of the left tibia (characters 76, 77, 78, and 79). Abbreviations: t. t.: tibial tuberosity; l. c.: lateral condyle; c. i. a.: caudal intercondylar area; a. p.: anterior projection. Scale bar = 1 cm.

**Figure 48.**
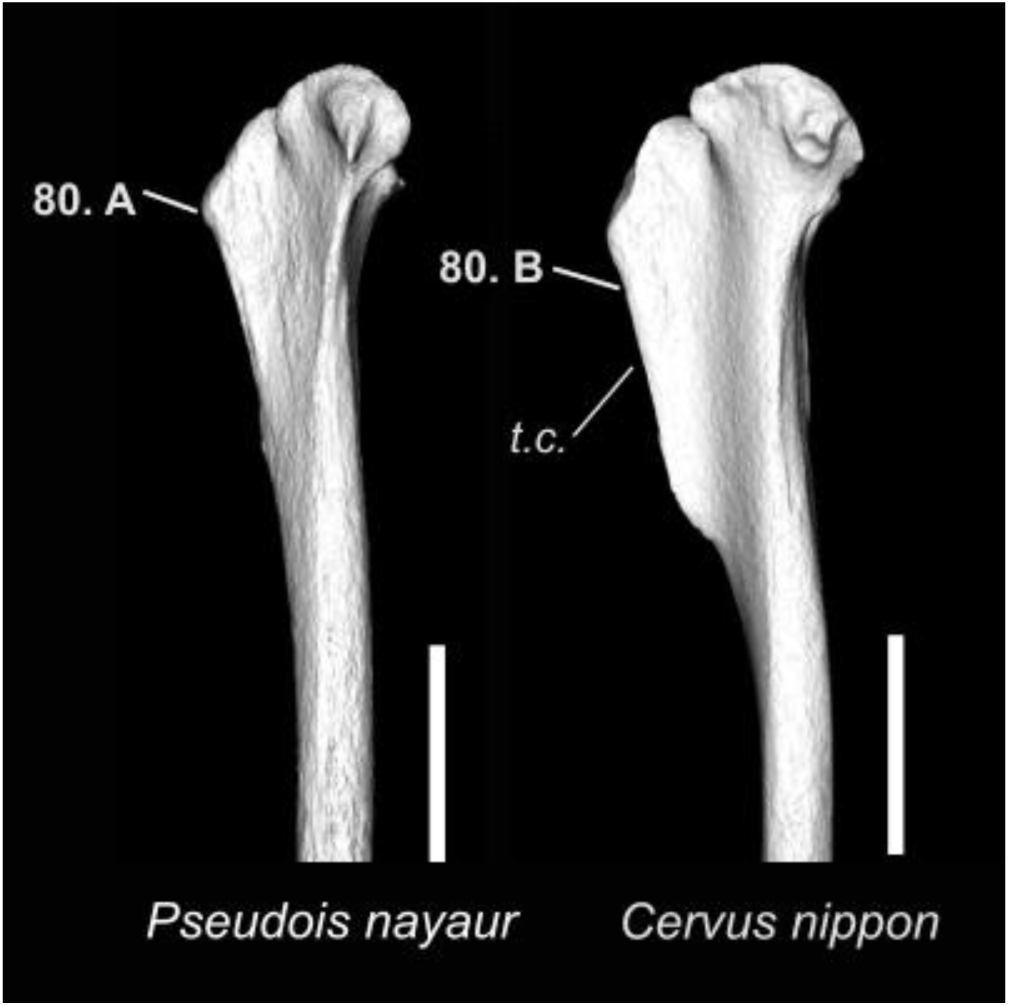
Character states visible on the lateral face of the left tibia (character 80). Abbreviations: t. c.: tibial crest. Scale bar = 5 cm.

**Figure 49.**
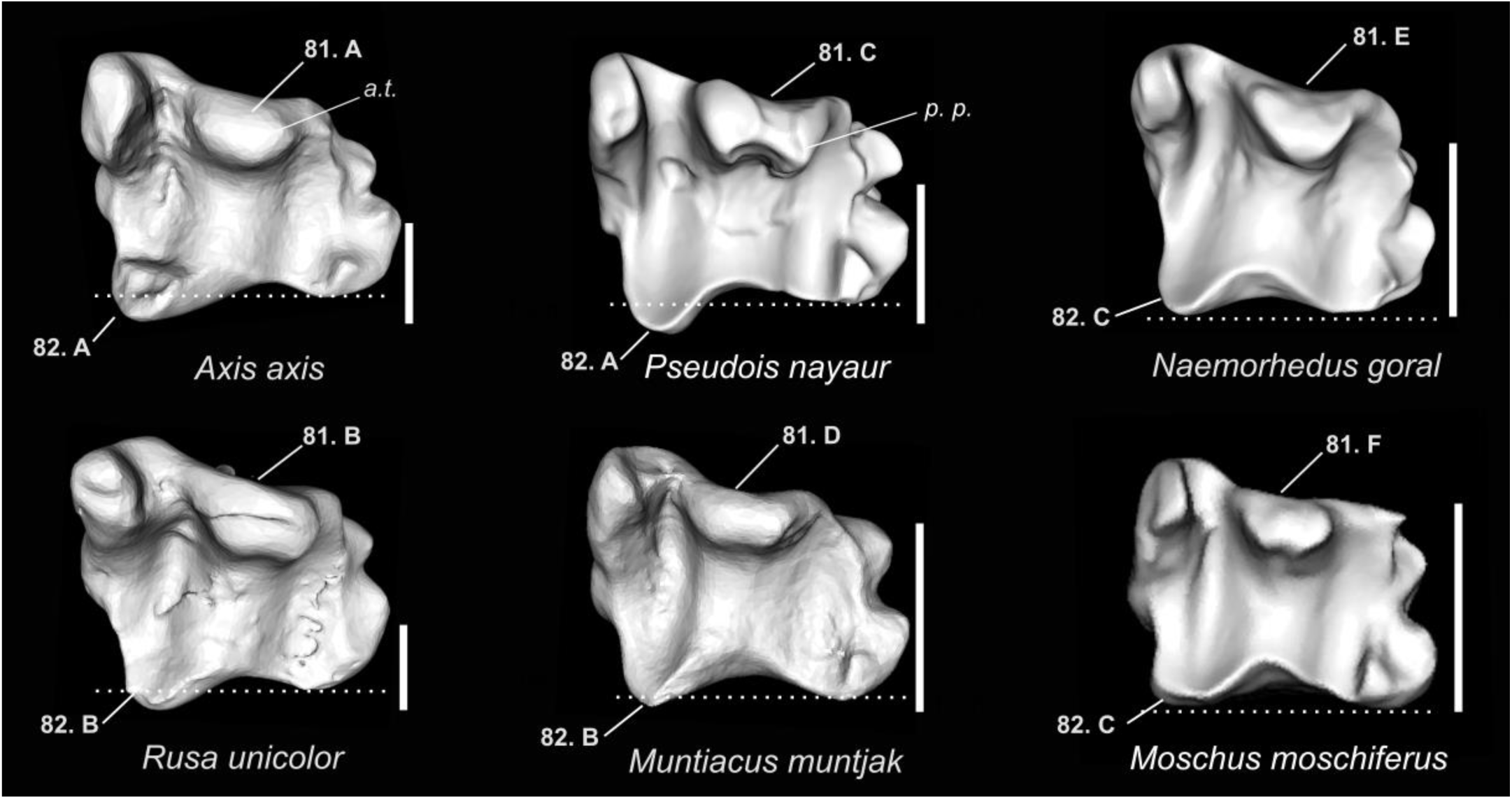
Character states visible on the ventral face of the left tibia (characters 81 and 82). Abbreviations: a. t.: anterior tubercle; p. p.: posterior projection. Scale bar = 1 cm.

**Figure 50.**
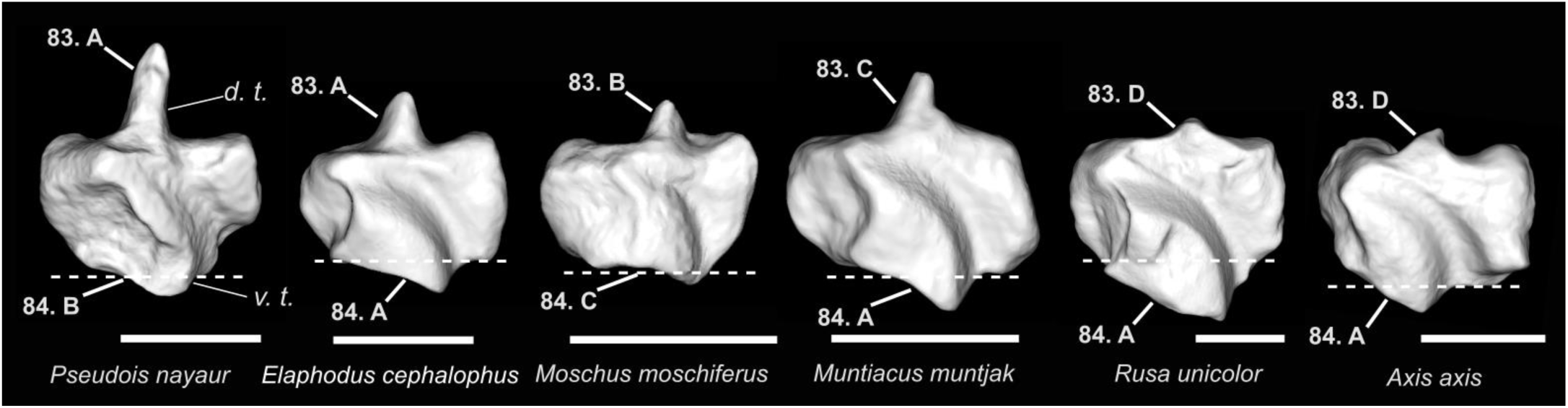
Character states visible on the medial face of the left malleolar bone (characters 83 and 84). Abbreviations: d. t.: dorsal tip; v. t.: ventral tip. Scale bar = 1 cm.

**Figure 51.**
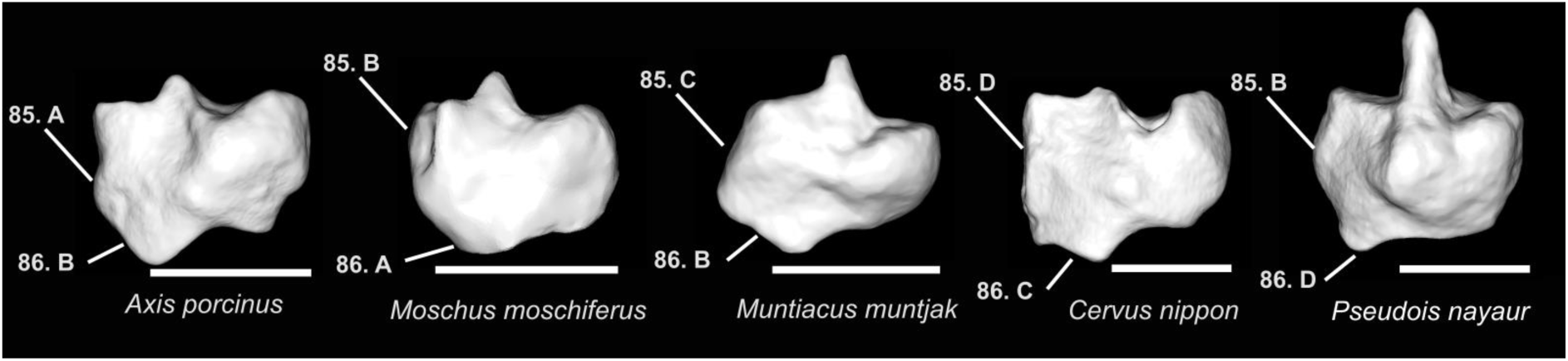
Character states visible on the lateral face of the left malleolar bone (characters 85 and 86). Scale bar = 1 cm.

**Figure 52.**
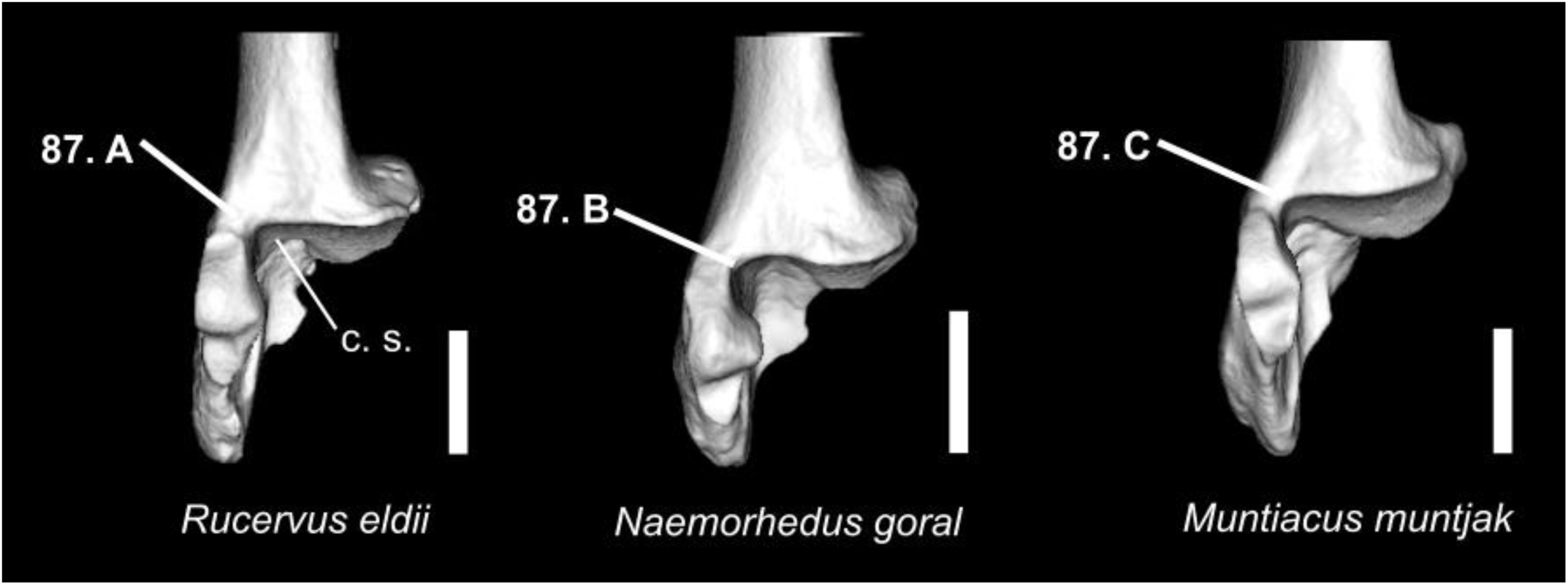
Character states visible on the dorsal face of the right calcaneus (character 87). Abbreviation: c. s.: calcaneus socket. Scale bar = 1 cm.

**Figure 53.**
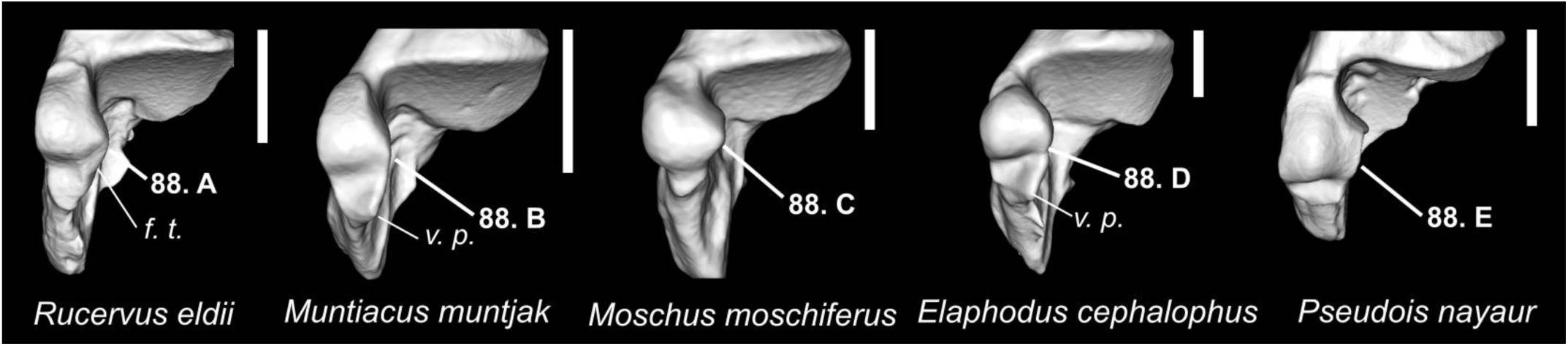
Character states visible on the dorsal face of the right calcaneus (character 88). Abbreviations: f. t.: fibular trochlea; v. p.: ventromedial projection. Scale bar = 1 cm.

**Figure 54.**
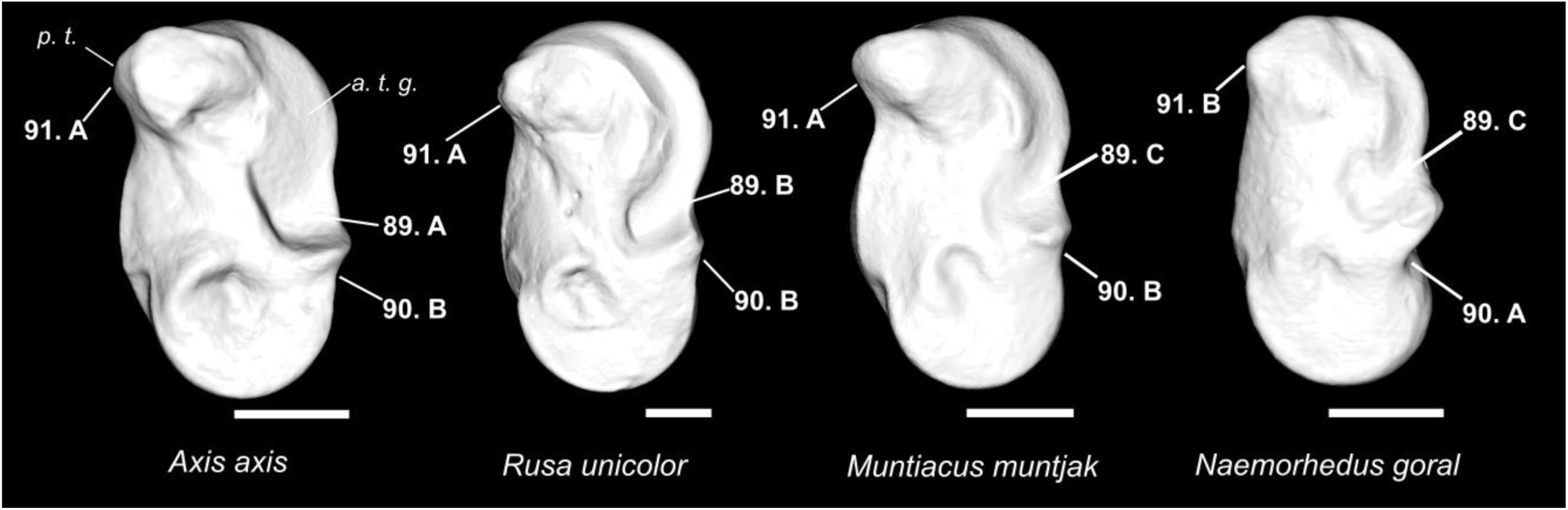
Character states visible on the medial face of the left astragalus (characters 89, 90, and 91). Abbreviations: a. t. g.: anterior trochlea groove; p. t.: proximal trochlea. Scale bar = 1 cm.

**Figure 55.**
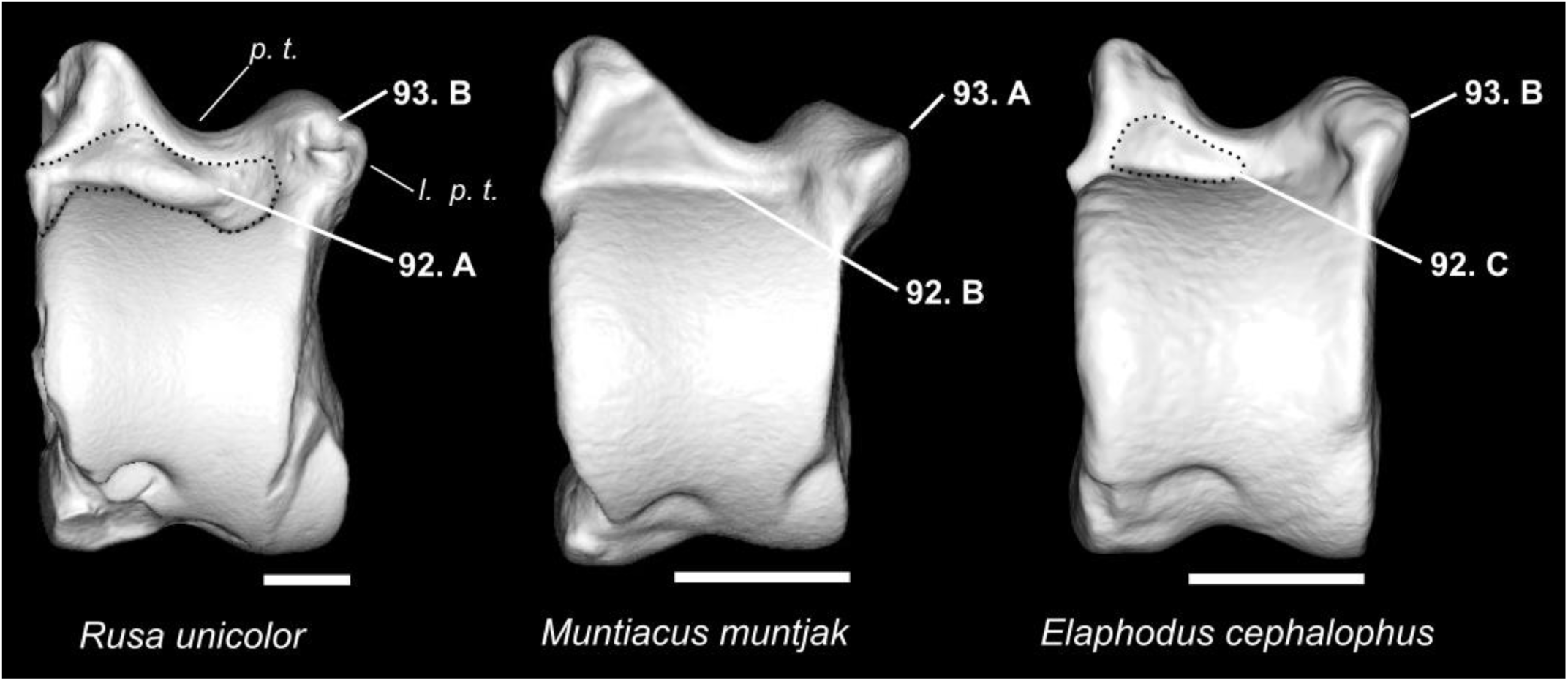
Character states visible on the posterior face of the left astragalus (characters 92 and 93). Abbreviations: p. t.: proximal trochlea; l. p. t.: lateral part of the proximal trochlea. Scale bar =1 cm.

**Figure 56.**
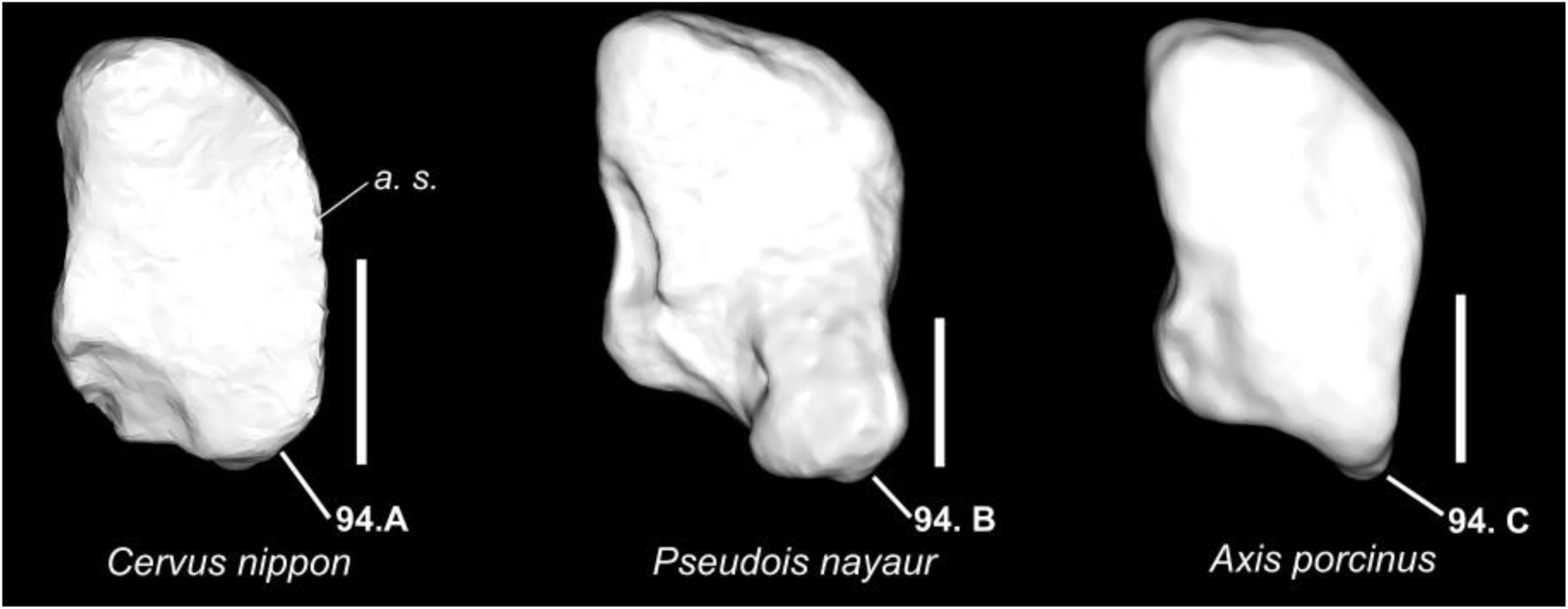
Character states visible on the dorsal face of the left large cuneiform (character 94). Abbreviations: a. s.: articular surface. Scale bar = 0.5 cm.

**Figure 57.**
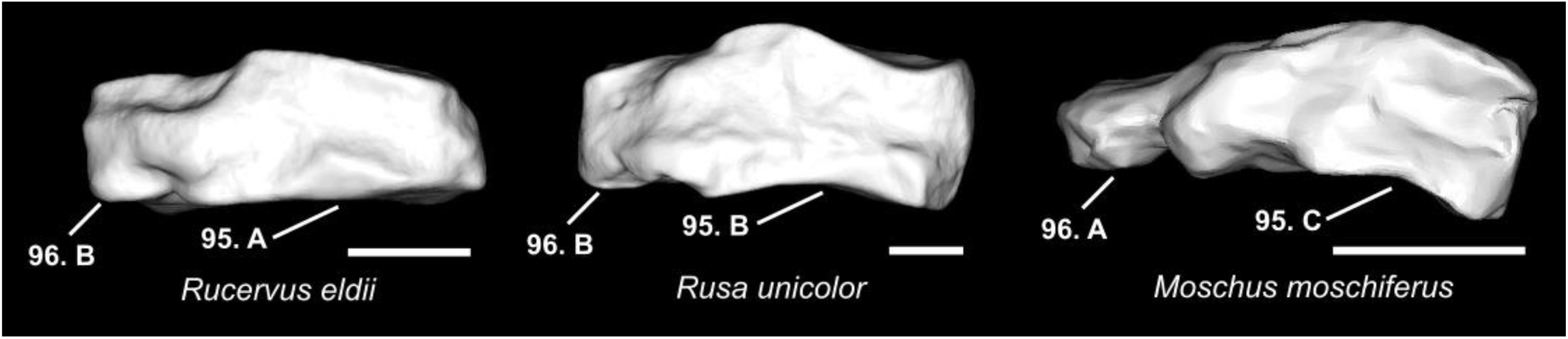
Character states visible on the medial face of the left large cuneiform (characters 95 and 96). Scale bar = 0.5 cm.

**Figure 58.**
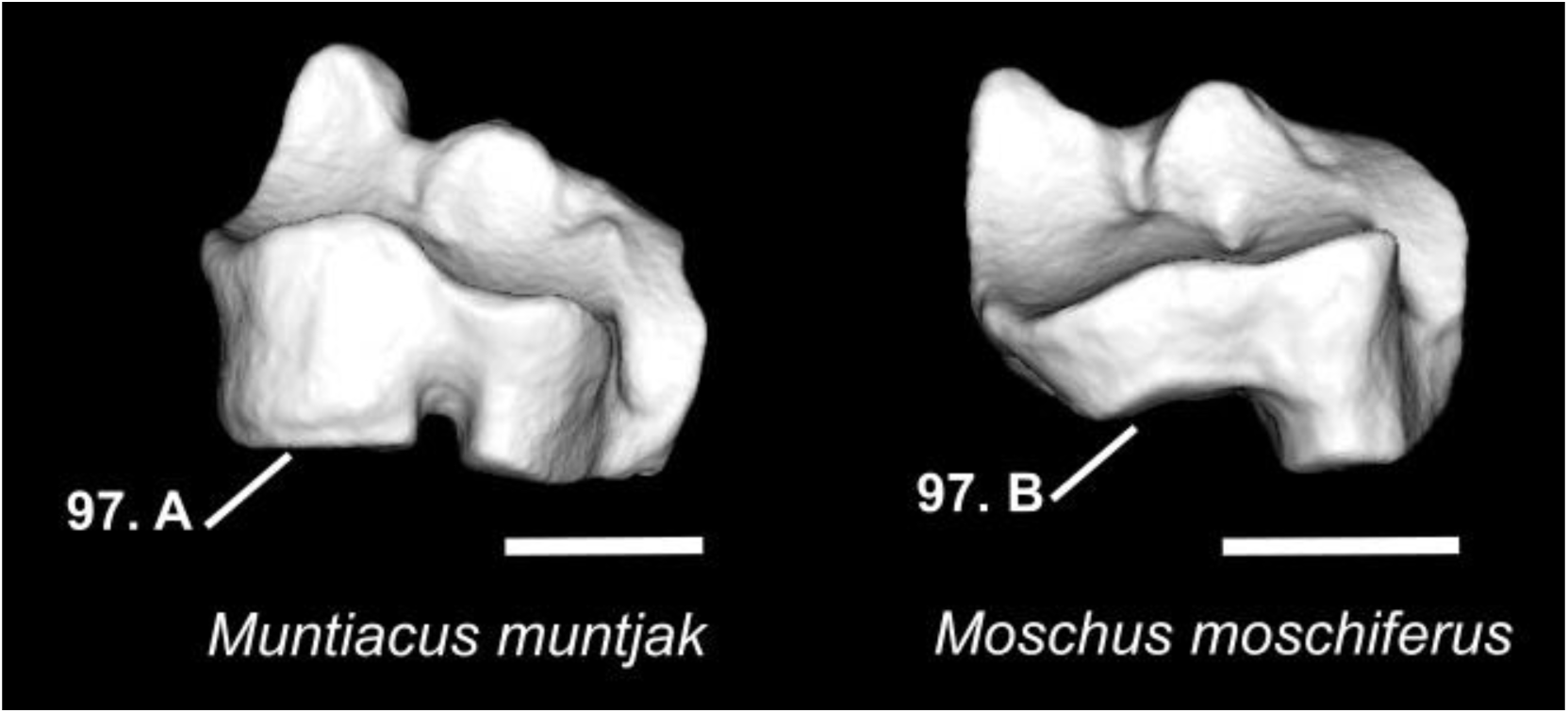
Character states visible on the anterior face of the left navicular-cuboid bone (character 97). Scale bar = 1 cm.

**Figure 59.**
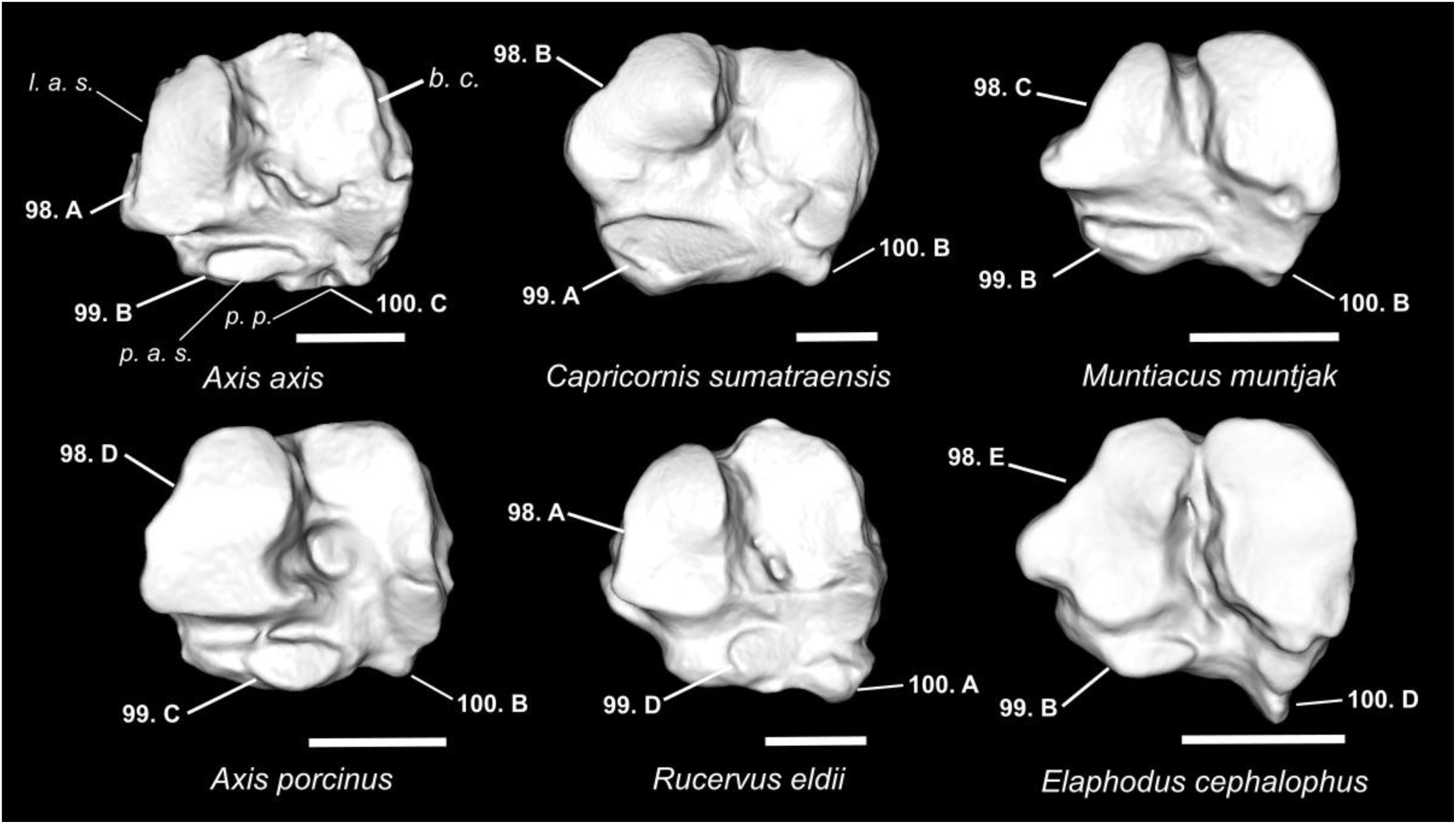
Character states visible on the ventral face of the navicular-cuboid bone (characters 98, 99, and 100). Abbreviations: l. a. s.: lateral articular surface; p. p.: posteromedial projection; p. a. s.: posterior articular surface; l. c.: large cuneiform. Scale bar = 1 cm.

**Figure 60.**
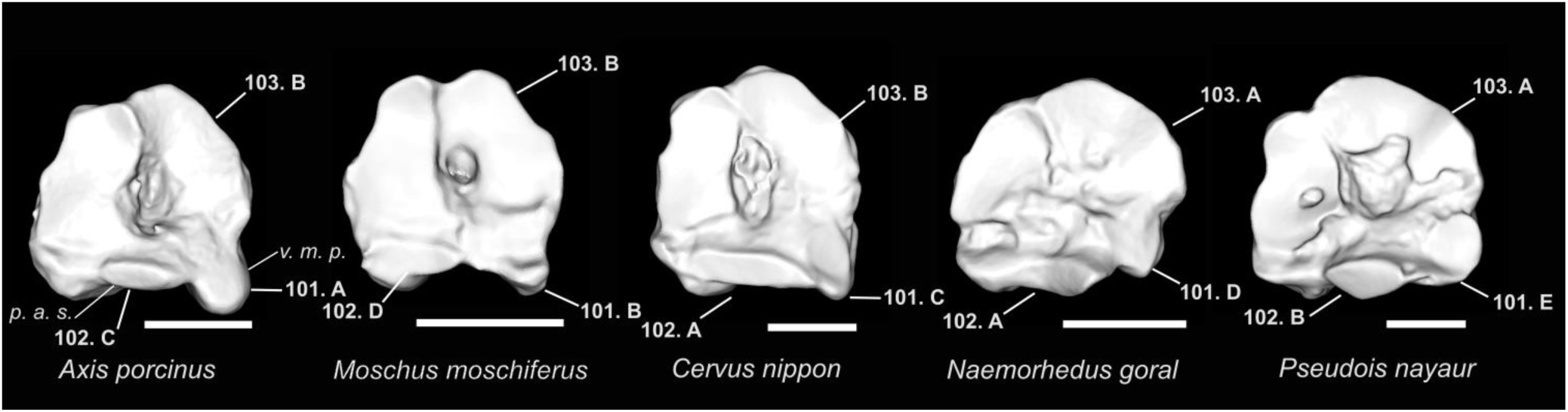
Character states visible on the dorsal face of the left metatarsus (characters 101, 102, and 103). Abbreviations: v. m. p.: ventromedial projection; p. a. s.: posterior articular surface. Scale bar = 1 cm.

**Figure 61.**
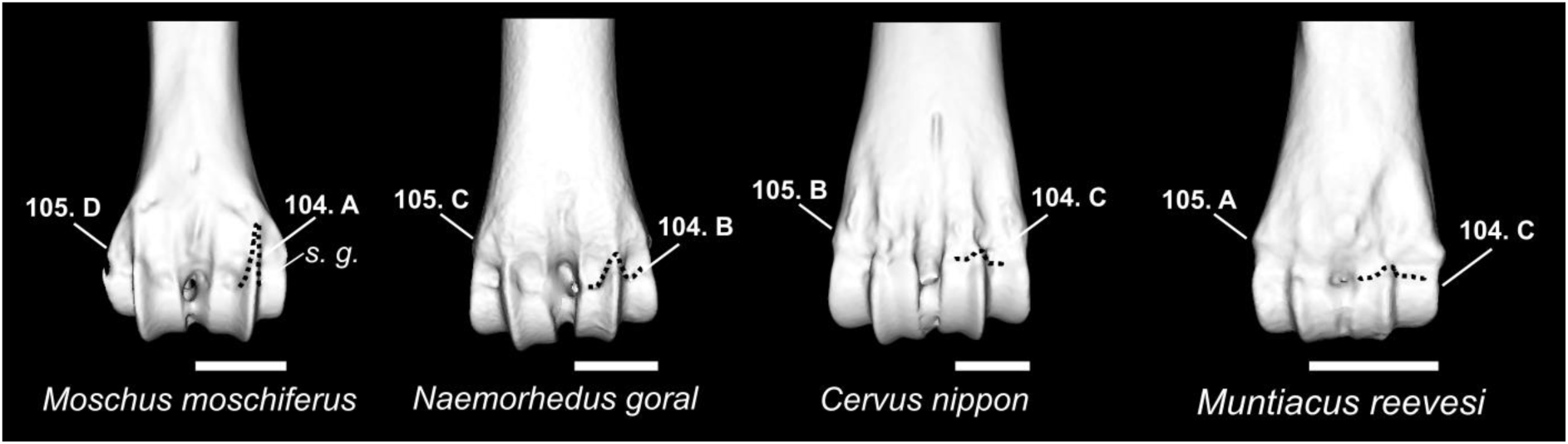
Character states visible on the posterior face of the distal extremity of the left metatarsus (Characters 104 and 105). Abbreviations: s. g.: sagittal ridge. Scale bar = 1 cm.

**Figure 62.**
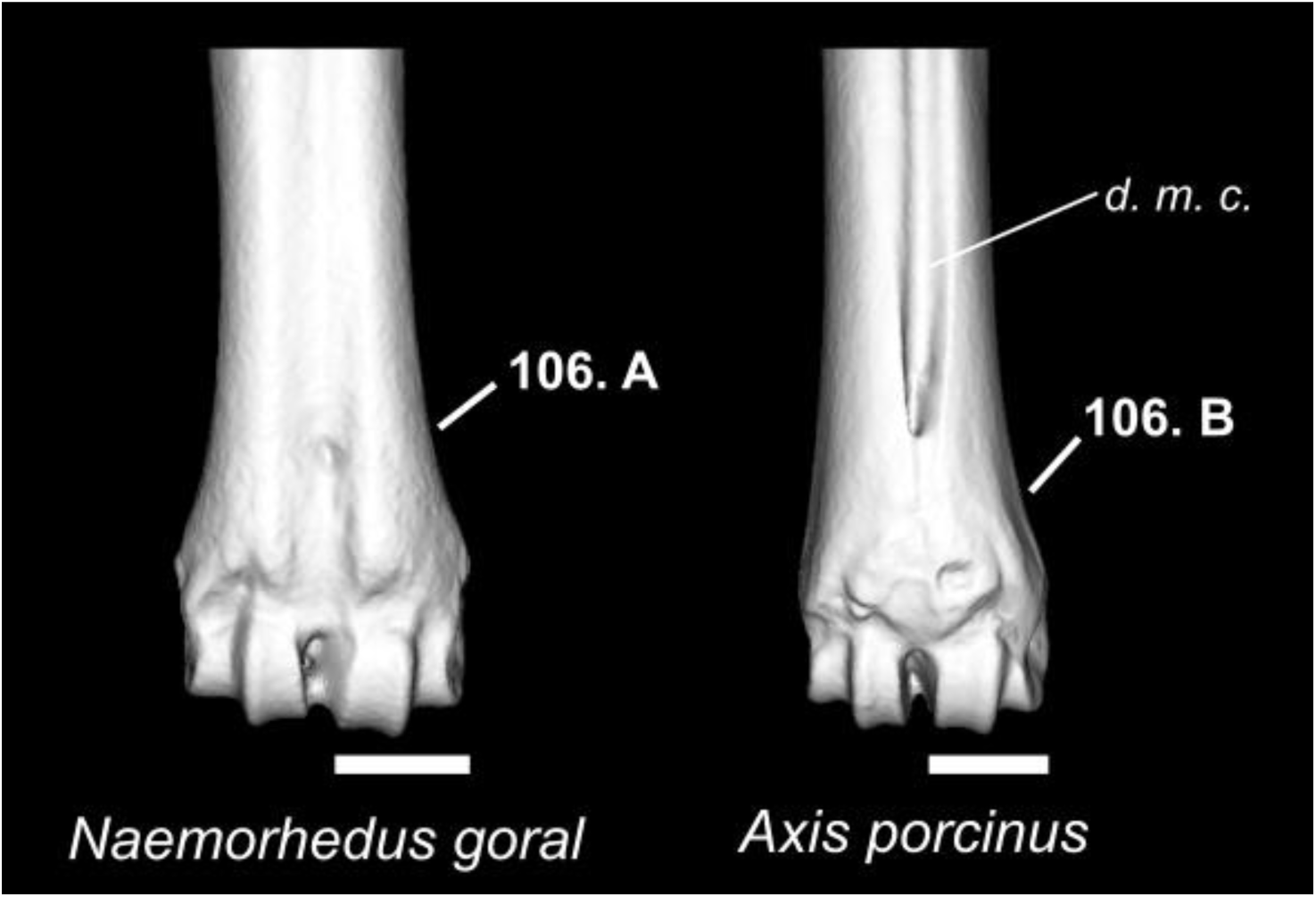
Character states visible on the anterior face of the distal area of the left metatarsus (character 106). Abbreviations: d. m. c.: distal metatarsal canal. Scale bar = 1 cm.

**Figure 63.**
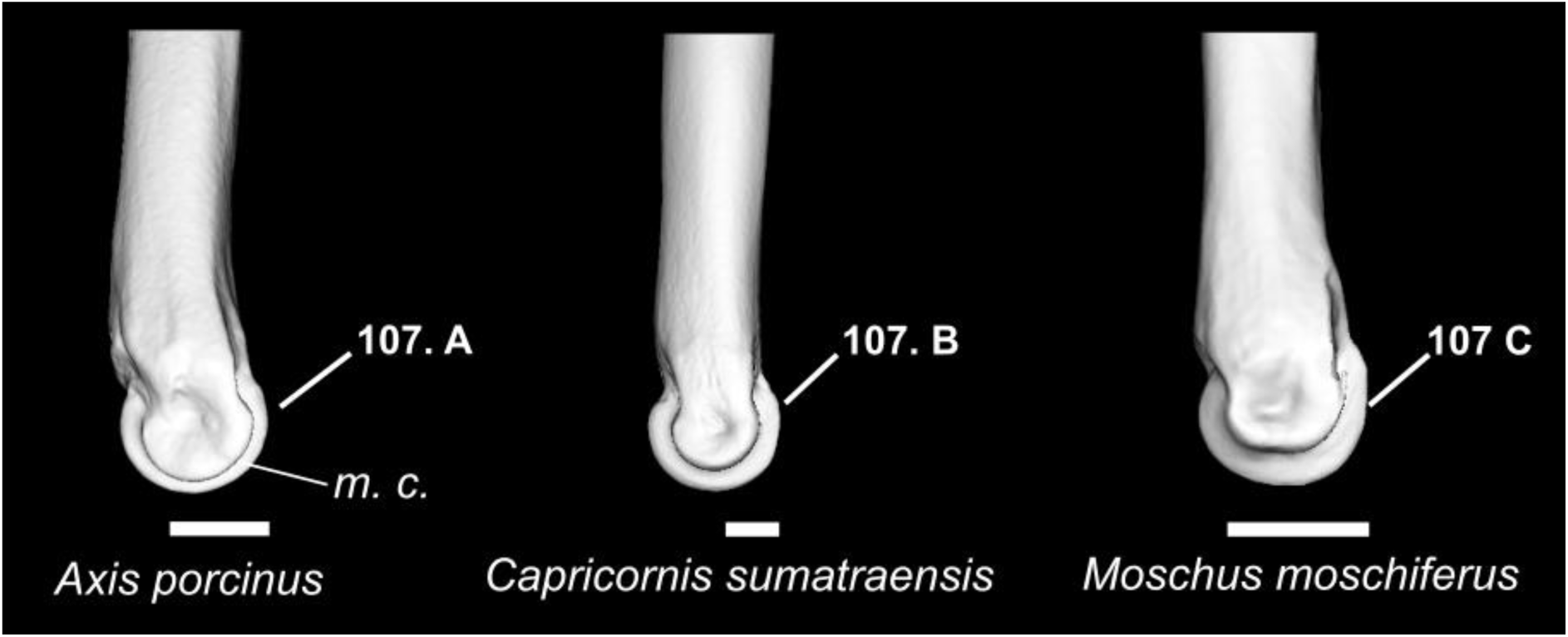
Character states visible on the medial face of the distal area of the left metatarsus (character 107). Abbreviations: m. c.: medial condyle. Scale bar = 1 cm.

**Figure 64.**
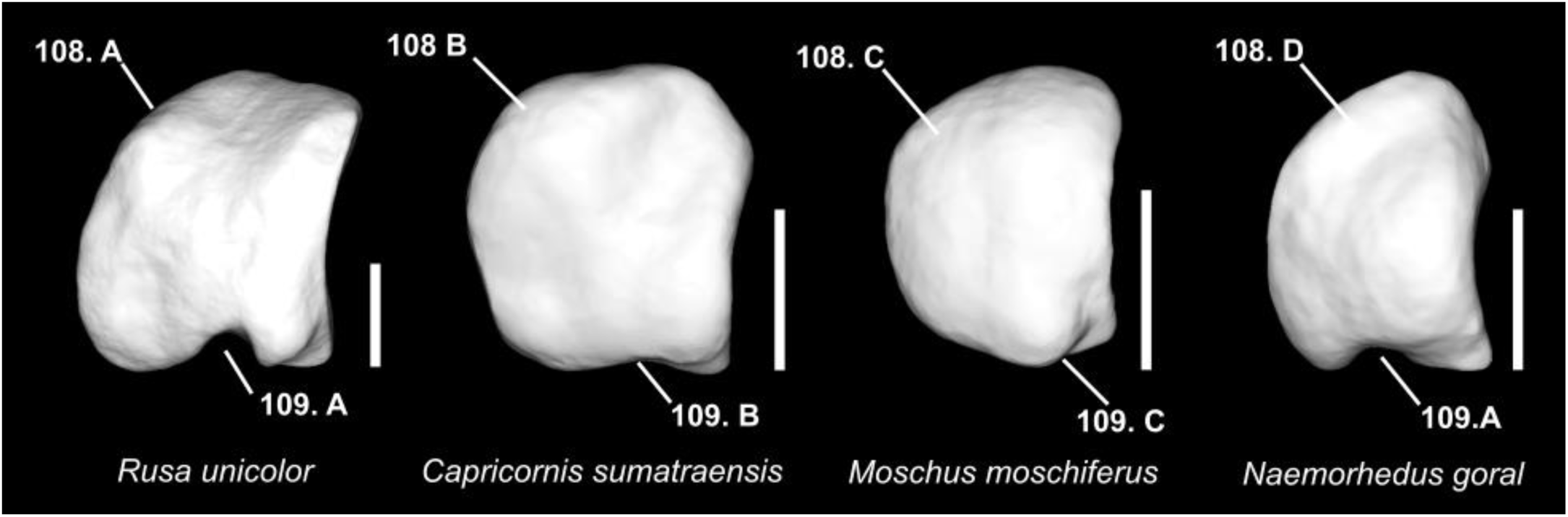
Character states visible on the lateral face of the large sesamoid (characters 108 and 109). Scale bar = 0.5 cm.

**Figure 65.**
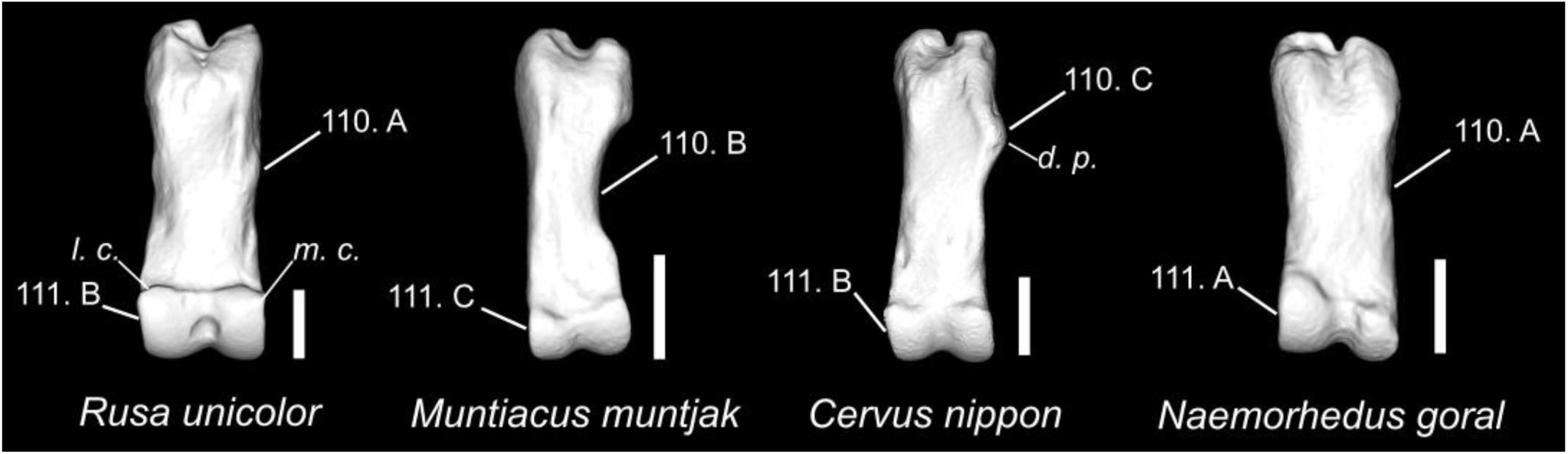
Character states are visible on the posterior face of the first phalanx. (Characters 110 and 111). Abbreviations: l. c.: lateral condyle; m. c.: medial condyle; d. p.: dorsomedial projection. Scale bar = 1 cm.

**Figure 66.**
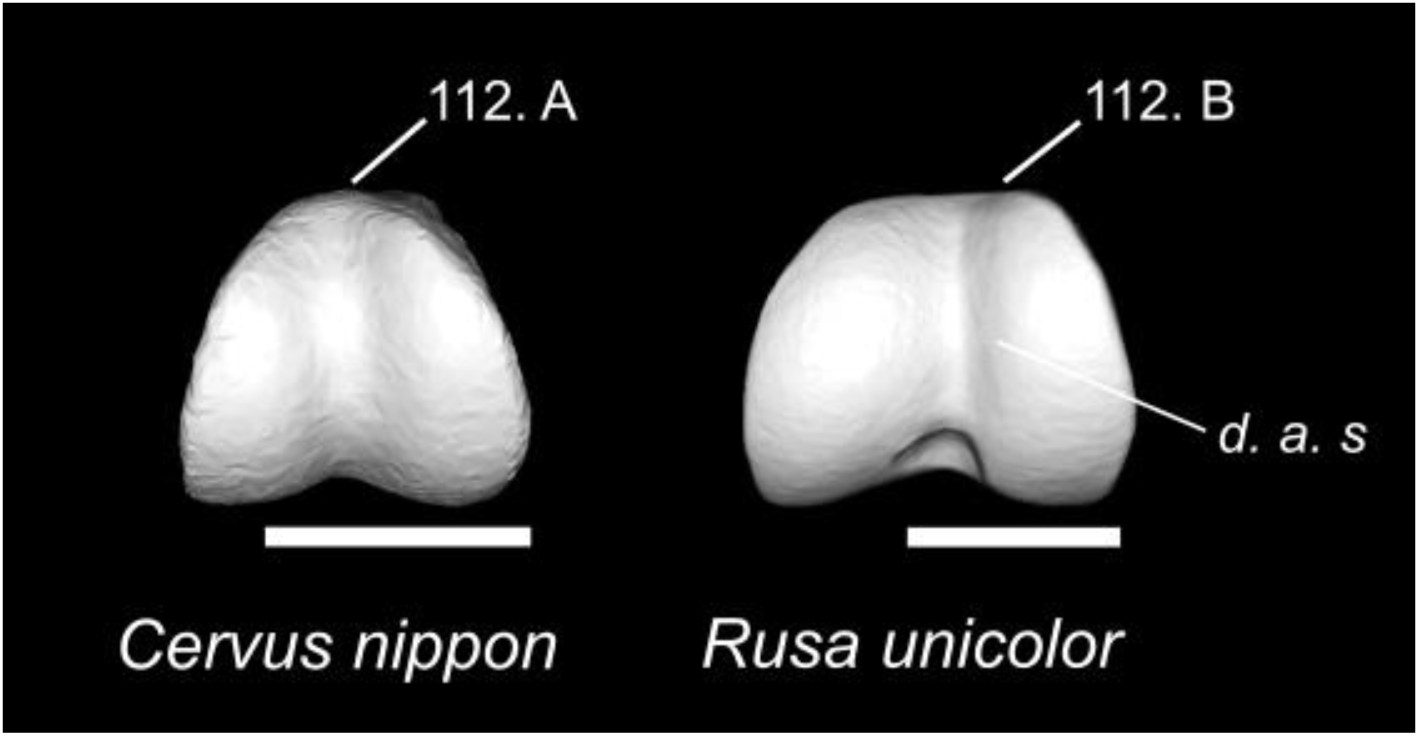
Character states are visible on the ventral face of the first phalanx. (Character 112). Abbreviations: d. a. s.: distal articular surface. Scale bar = 1 cm.

**Figure 67.**
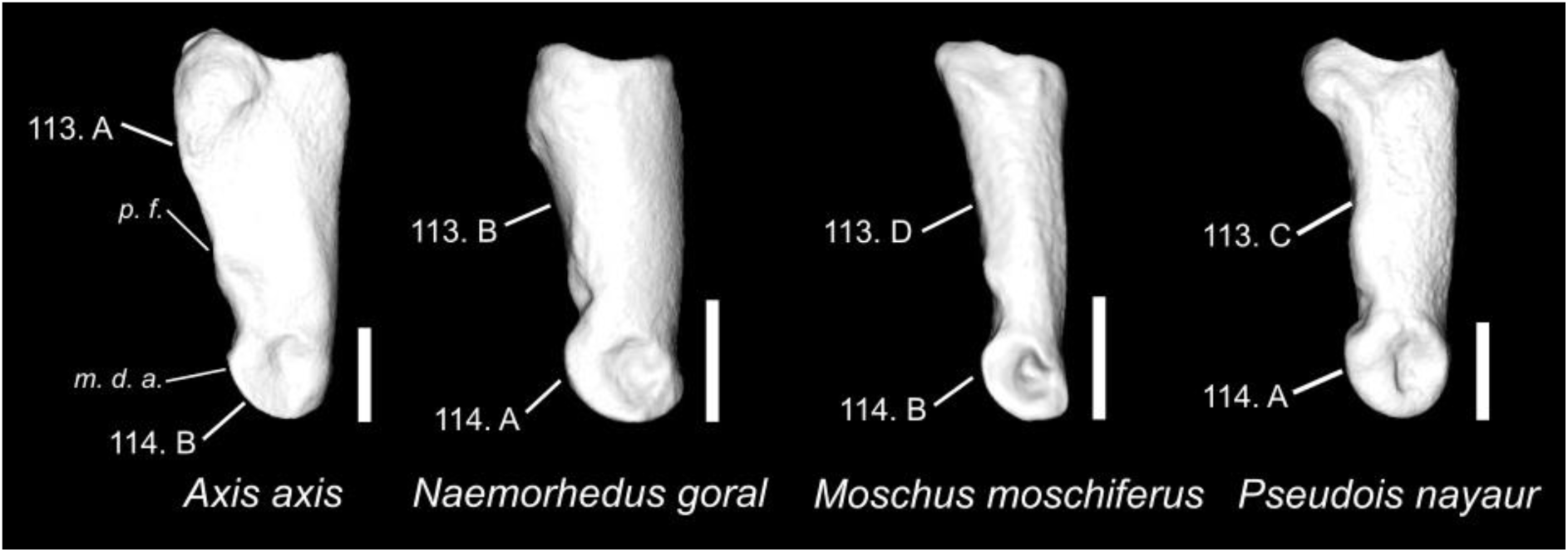
Character states are visible on the axial face of the first phalanx. (Characters 113 and 114). Abbreviations: p. f.: palmar face; m. d. a.: medial distal articular facet. Scale bar = 1 cm.

**Figure 68.**
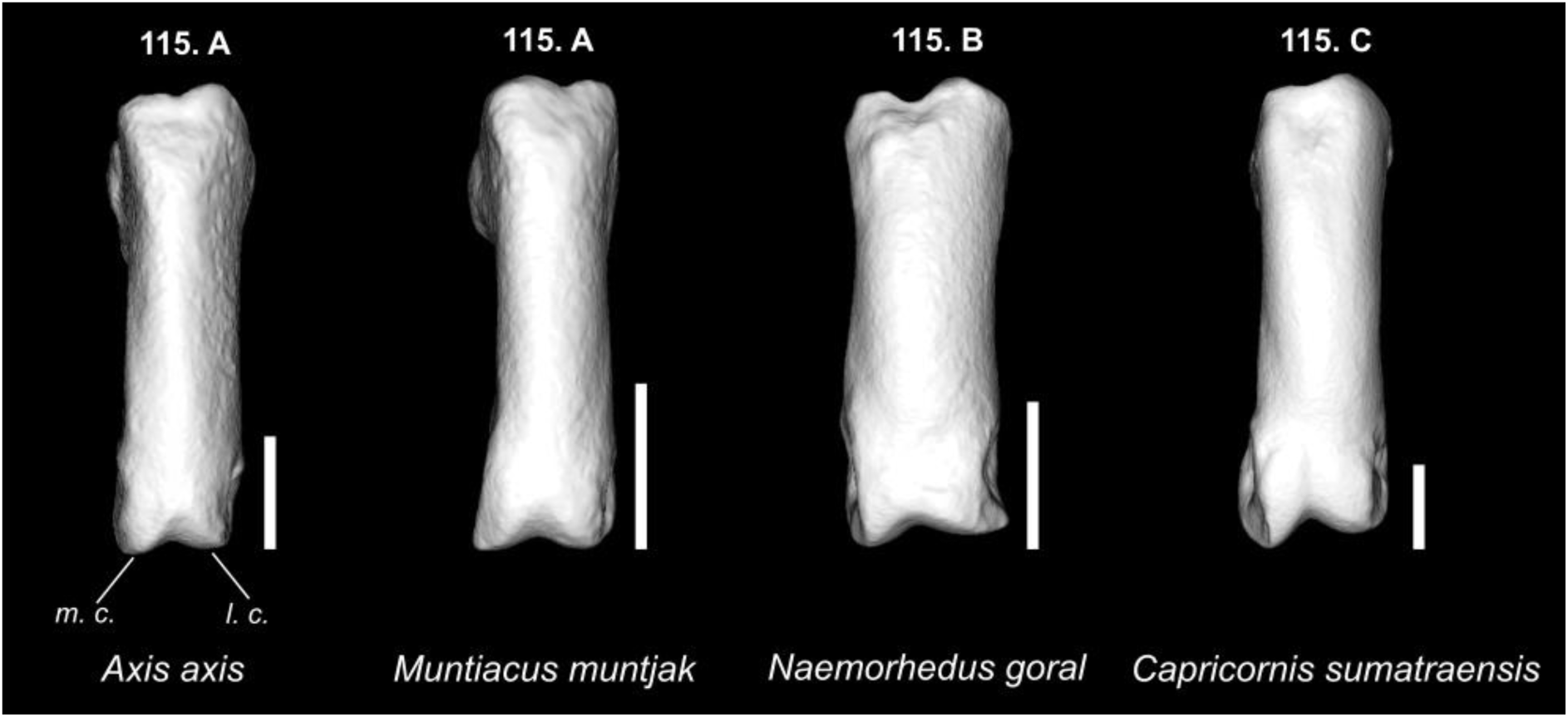
Character states are visible on the anterior face of the first phalanx. (Character 115). Abbreviations: m. c.: medial condyle; l. c.: lateral condyle. Scale bar = 1 cm.

**Figure 69.**
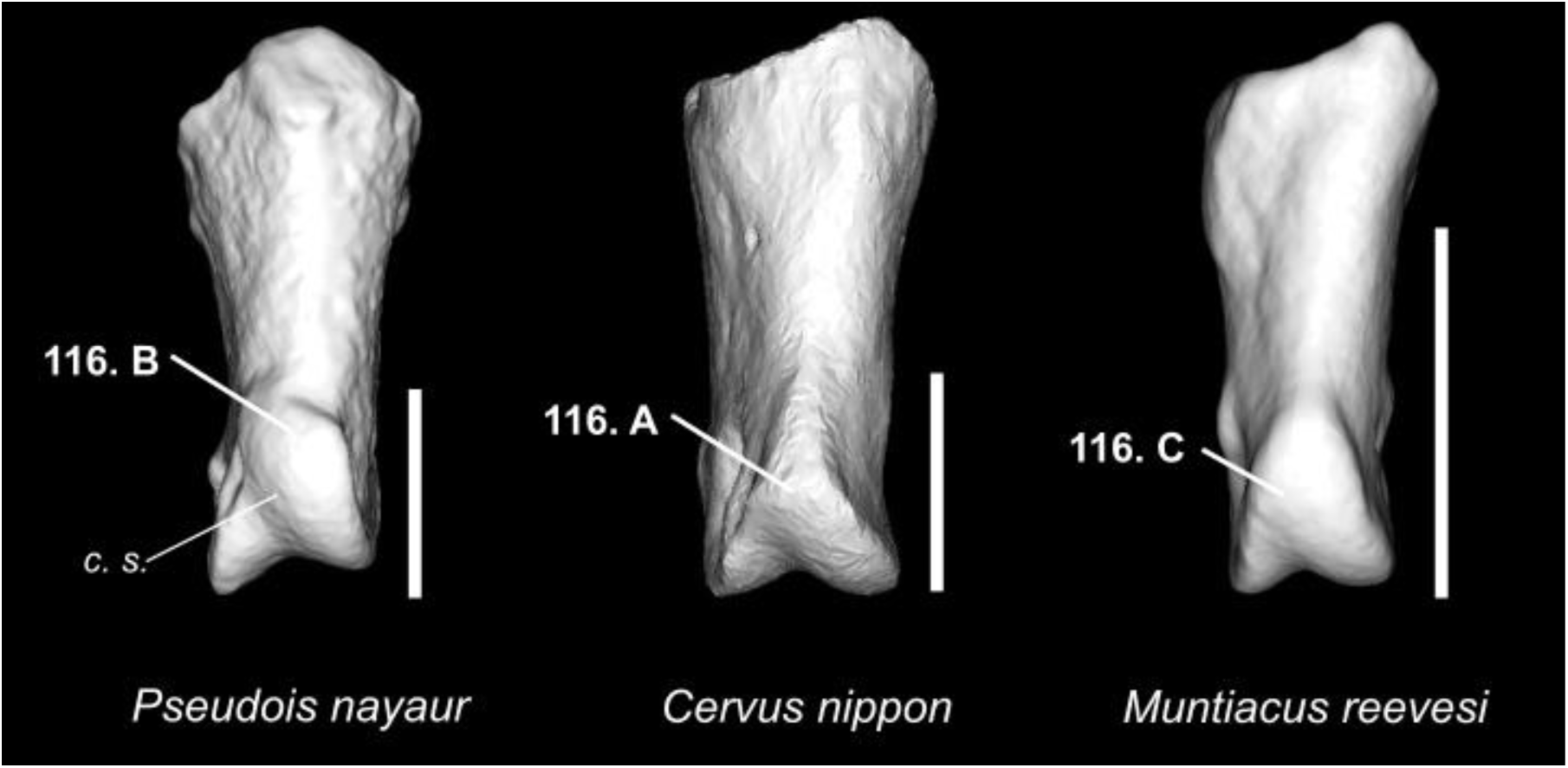
Character states are visible on the anterior face of the second phalanx. (Character 116). Abbreviations: m. c.: medial condyle; l. c.: lateral condyle. Scale bar = 1 cm.

**Figure 70.**
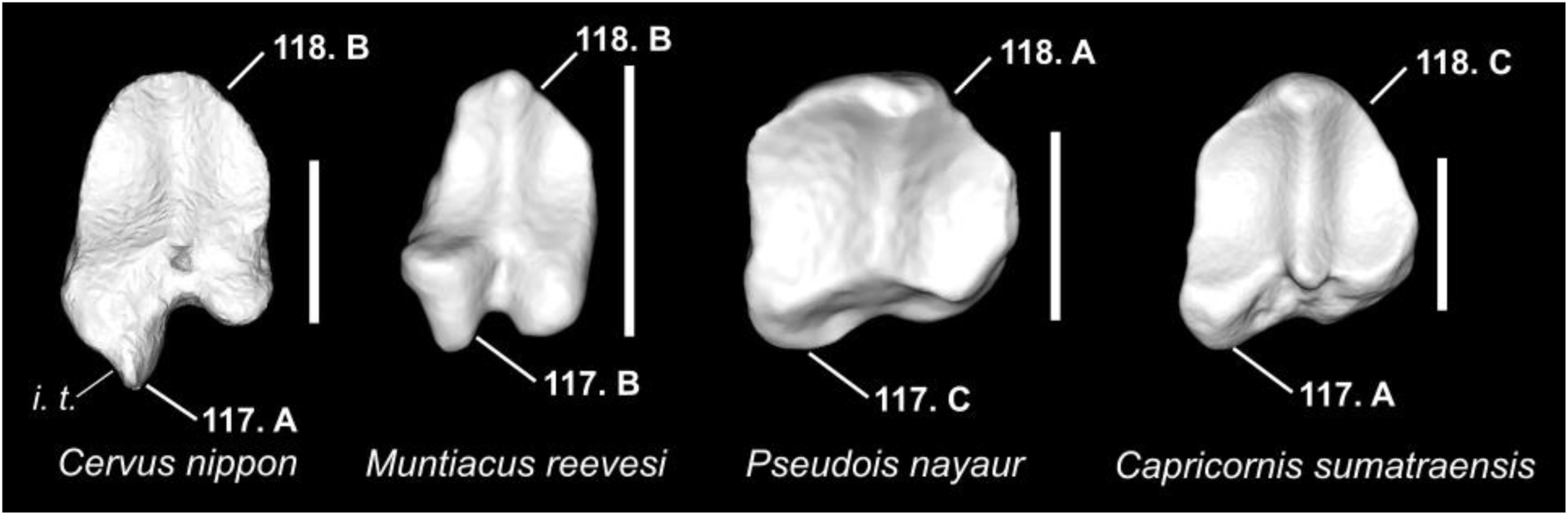
Character states are visible on the dorsal face of the second phalanx. (Characters 117 and 118). Abbreviations: i. t.: insertion tubercle. Scale bar = 1 cm.

**Figure 71.**
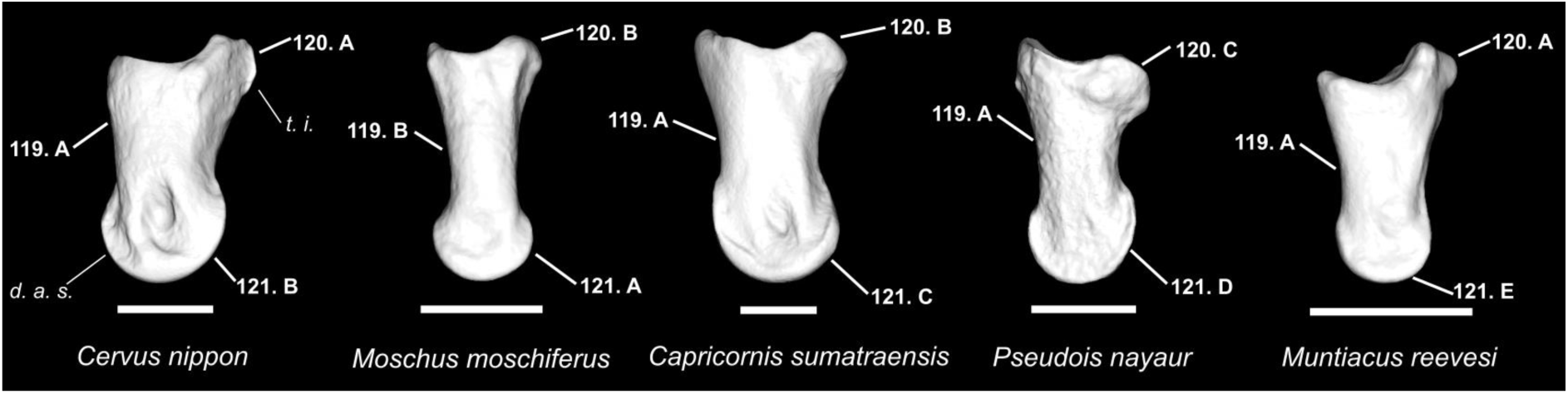
Character states are visible on the axial face of the second phalanx. (Characters 119, 120, and 121). Abbreviations: i. t.: insertion tubercle; d. a. s.: distal articular surface. Scale bar = 1 cm.

**Figure 72.**
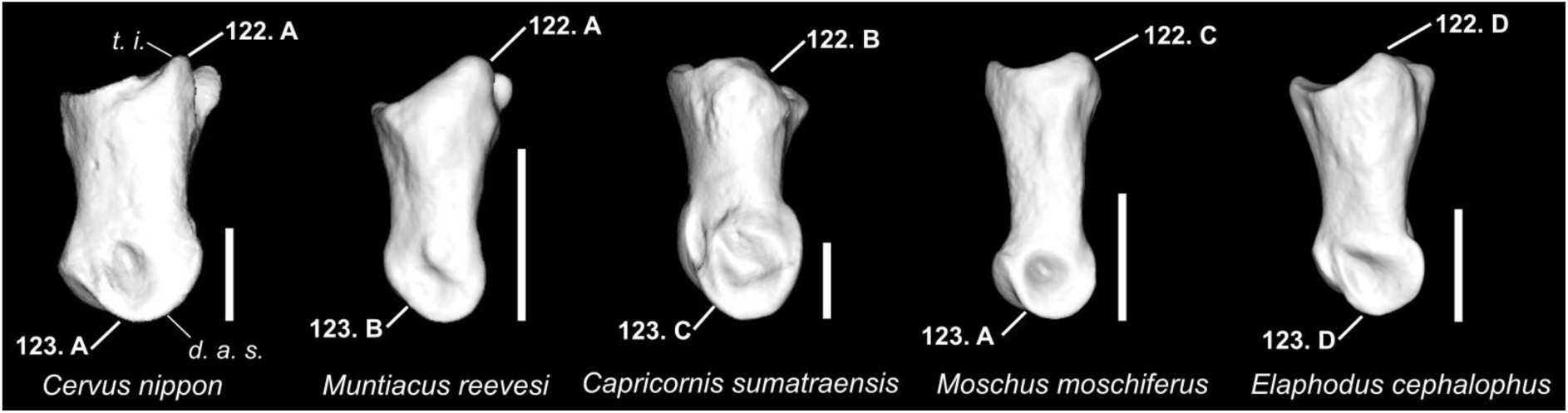
Character states are visible on the abaxial face of the second phalanx. (Characters 122 and 123). Abbreviations: i. t.: insertion tubercle; d. a. s.: distal articular surface. Scale bar = 1 cm.

**Figure 73.**
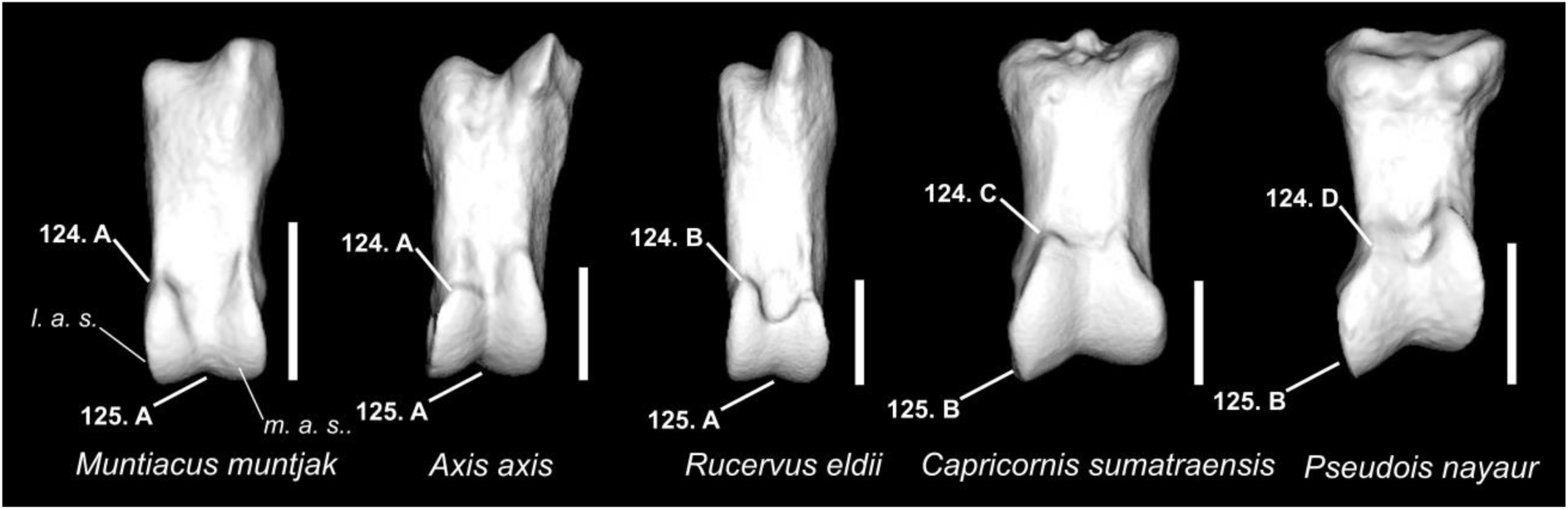
Character states are visible on the posterior face of the second phalanx. (Characters 124 and 125). Abbreviations: l. a. s.: lateral articular surface; m. a. s.: medial articular surface. Scale bar = 1 cm.

**Figure 74.**
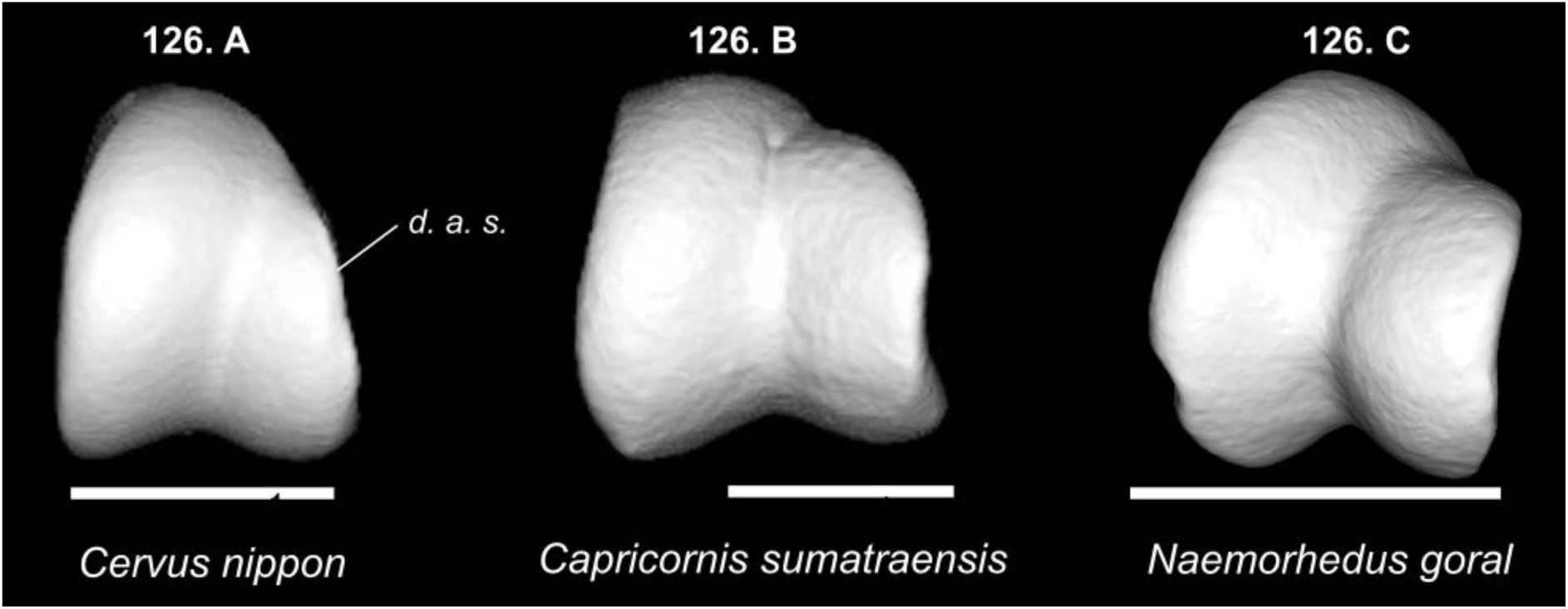
Character states are visible on the ventral face of the second phalanx. (Character 126). Abbreviations: d. a. s.: distal articular surface. Scale bar = 1 cm.

**Figure 75.**
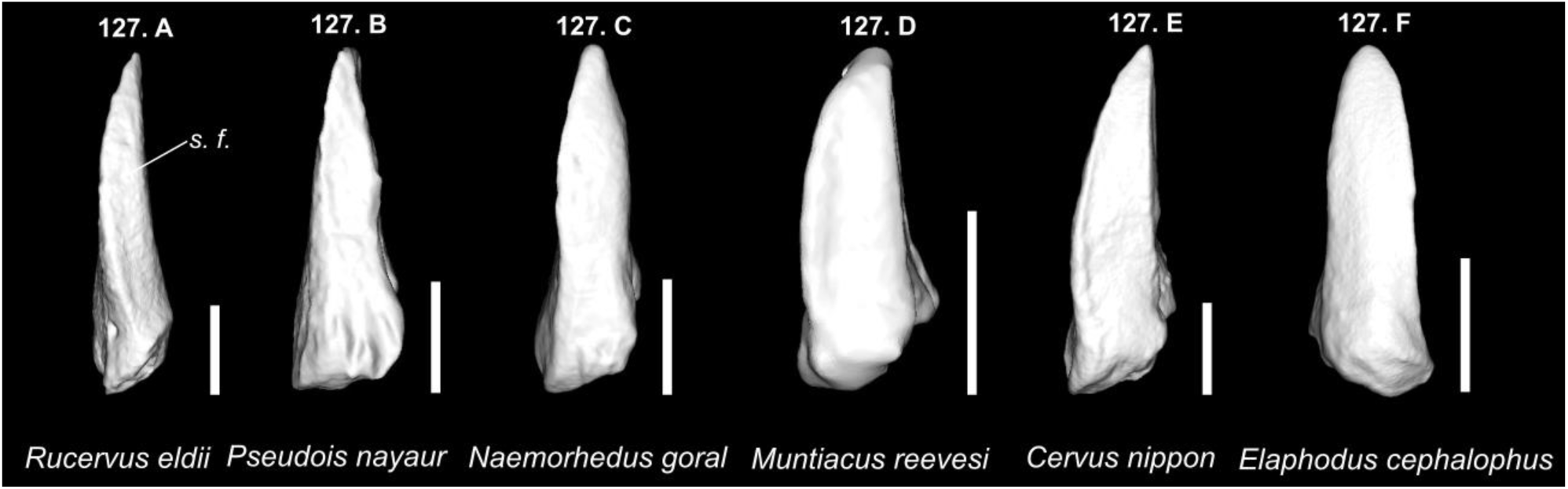
Character states are visible on the ventral face of the third phalanx. (Character 127). Abbreviations: s. f.: solar face. Scale bar = 1 cm.

**Figure 76.**
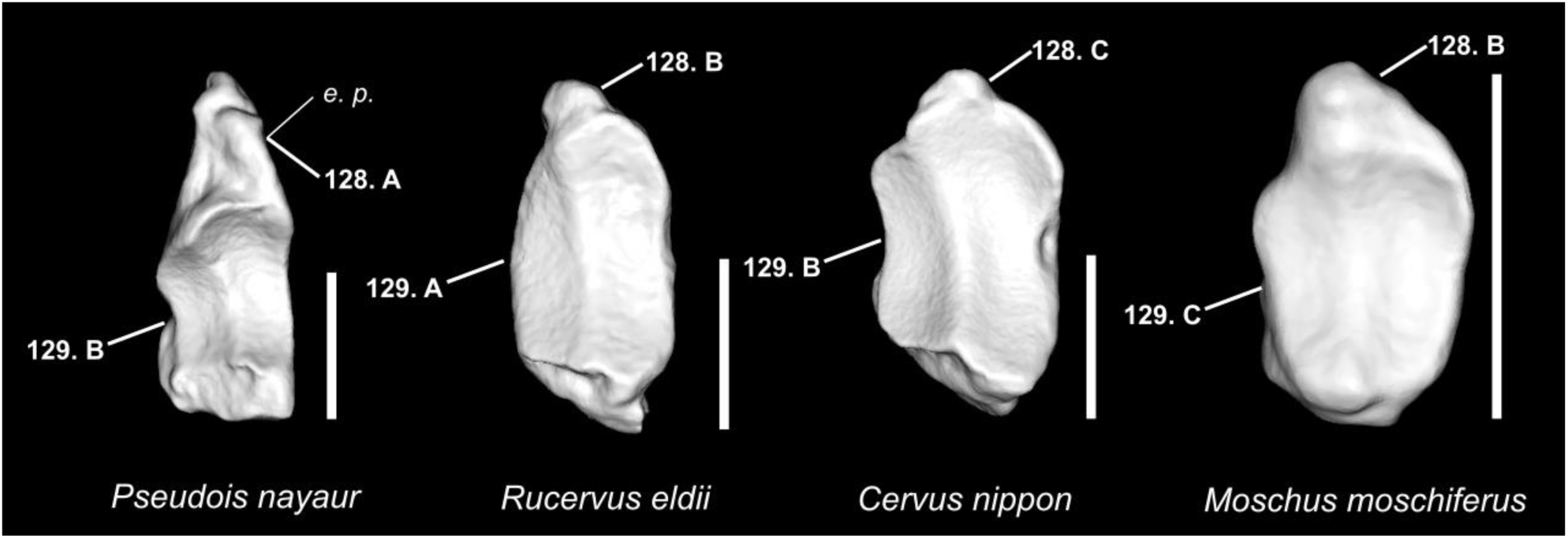
Character states are visible on the posterior face of the third phalanx. (Characters 128 and 129). Abbreviations: e. p.: extensor process. Scale bar = 1 cm.

**Figure 77.**
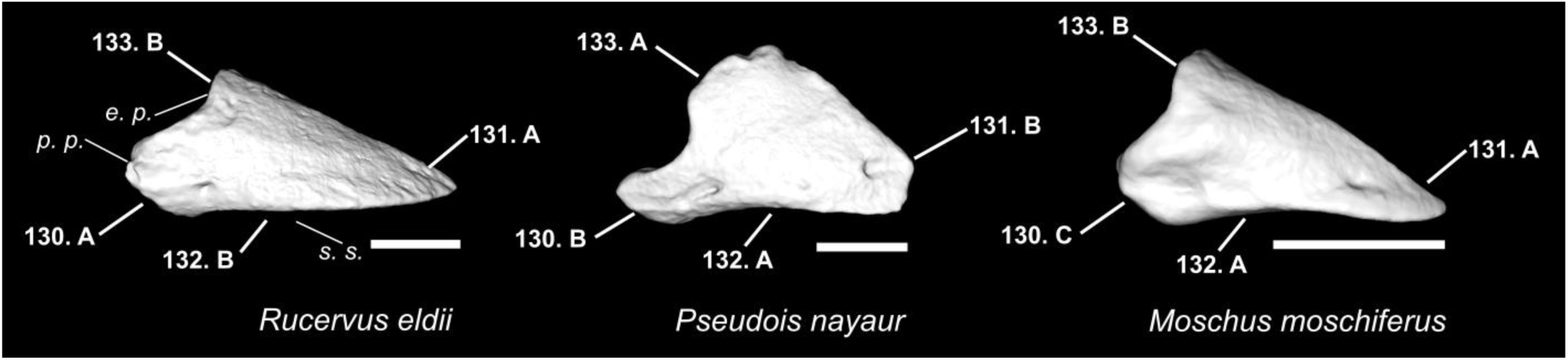
Character states are visible on the lateral face of the third phalanx. (Characters 130, 131, 132, and 133). Abbreviations: p. p.: palmar process; s. s.: solar surface; e. p.: extensor process. Scale bar = 1 cm.

**Figure 78.**
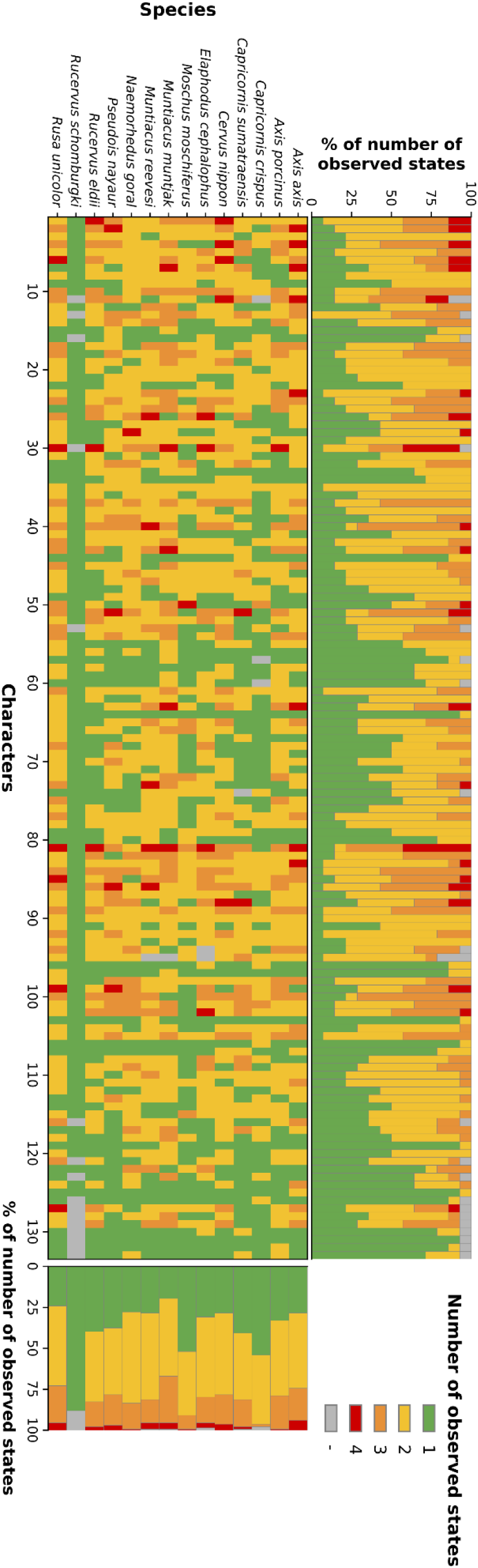
Number of observed states for each character in each taxon.

Nevertheless, 22 character states were still unique to a single taxon and occurred in more than 70 % of its specimens (**Table 01**). Five of these character states appeared useful for identification at the species level. These character states were observed in *Axis porcinus* (17: E), *Pseudois nayaur (28: C and 131: B), and Moschus moschiferus (28: E and 105: D)*. At the genus scale, only a single character state was relevant to identify the genus Muntiacus (121: E). Regarding families, with the exception of Moschidae (identified by the same character states as *Moschus moschiferus*), 16 characters were considered relevant (Table 01) to identify the Bovidae/Caprinae (6 characters) and Cervidae (10 characters) families. If character states occurring in between 25 and 70% of the specimens of each considered taxon or clade are also considered (**Table 01**; **Figure 79**), we accounted for 12 autapomorphic character states corresponding to 11 characters. The remaining specimens show one or more alternative, non-exclusive states of the same character. We also observed 37 synapomorphic characters, presenting 46 synapomorphic character states unique to a supraspecific clade (**Table 01**; **Figure 79**).

**Figure 79.**
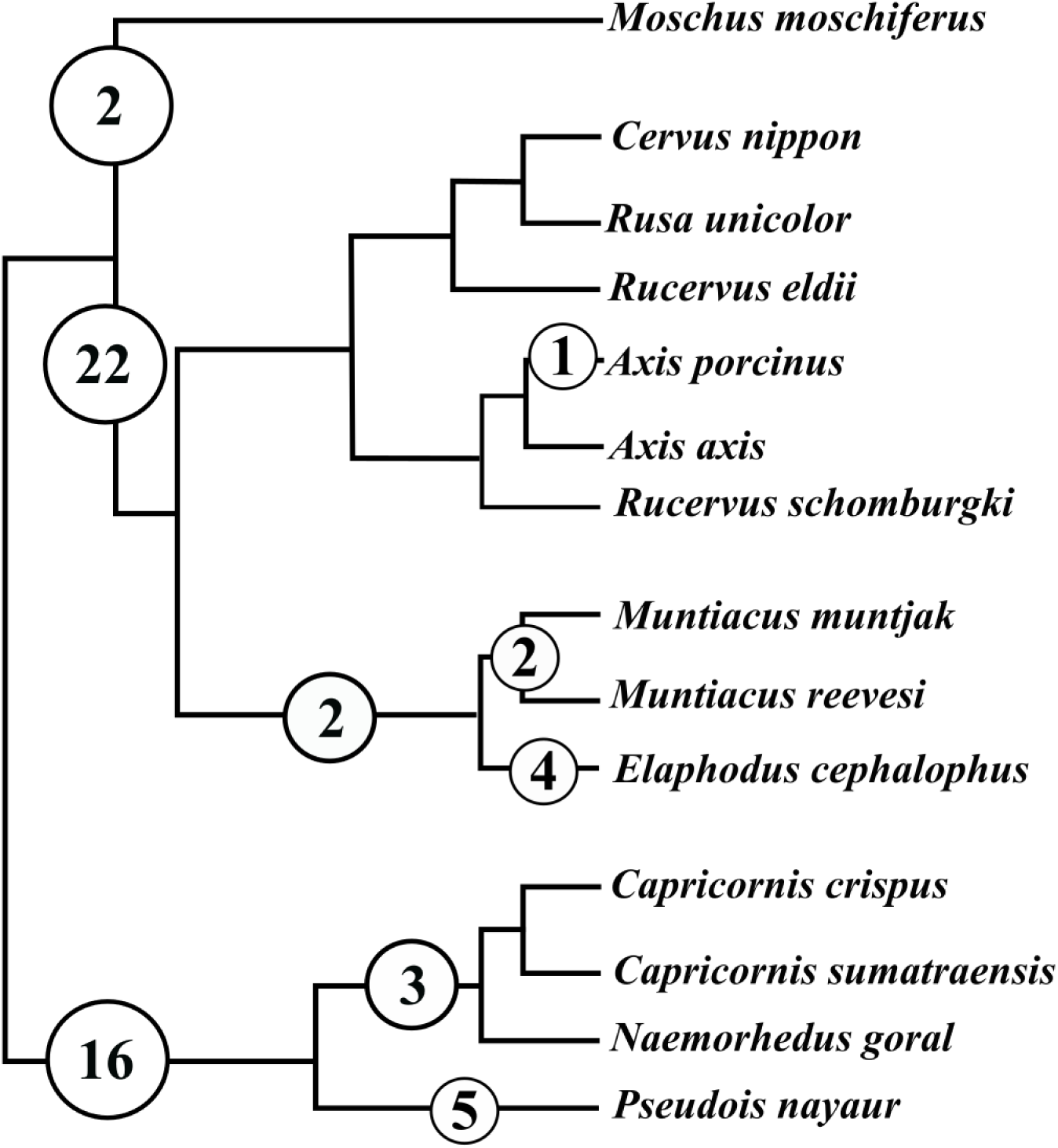
Distribution of the autapomorphic and synapomorphic character states listed in **Table 01** across the phylogeny. The backbone tree is a composite of Fernández & Vrba (2005) for the higher phylogeny, Heckeberg (2020) for the Cervidae, and Calamari (2021) for the Bovidae.

**Table 01.** Character states observed as unique to a given taxon, with the frequency of their occurrence in all observed specimens of that taxon.

| Taxon | Freq. 100% | Freq. 99-90% | Freq. 89-80% | Freq. 79-70% | Freq. 69-60% | Freq. 59-50% | Freq. 49-25% |
| --- | --- | --- | --- | --- | --- | --- | --- |
| **Bovidae/ Caprinae** | 57:B | 130:B;<br>106:A |  | 126:B;<br>87:B; 11:E |  | 131:B;<br>124:D | 120:C; 101:D; 101:E;<br>28:C; 14:C; 13:E;<br>6:D; 2:D |
| **Cervidae** |  | 125:A;<br>124:A;<br>107:A;<br>64:B; 57:A;<br>55:A | 120:A;<br>73:A; 62:B | 60:A | 72:A | 121:B;<br>65:B;<br>12:A | 127:D; 127:E; 116:A;<br>112:A; 97:A; 81:A;<br>45:B; 30:C; 6:C |
| **Moschidae/ *Moschus moschiferus*** | 28:E | 105:D |  |  |  |  |  |
| **Muntiacinae** |  |  |  |  |  |  | 100:D; 63:F |
| **Ovibovini** |  |  |  |  |  |  | 126:C; 44:B; 28:F |
| ***Muntiacus*** |  |  | 121:E |  | 123:B |  |  |
| ***Pseudois nayaur*** | 131:B | 28:C |  |  | 54:D;<br>14:C | 6:D |  |
| ***Elaphodus cephalophus*** |  |  |  |  | 127:F | 122:D | 98:E; 51:E |
| ***Axis porcinus*** |  |  |  | 17:E |  |  |  |

The maximum frequencies of the character states show strong disparities between characters. When all taxa are considered together, the mean maximum frequency is 82% with a standard deviation of 19% and an interquartile range of 67–100%. This spread thus reflects genuine, widespread heterogeneity across characters rather than a few outliers. In the dataset, 21 characters exhibit frequencies above 95%, indicating high stability, while 17 characters fall below 70%. This shows that it is difficult to reduce the morphology of the anatomical structures of a given taxon to a single character state and highlights the general morphological variability impacting all of the bones investigated in our dataset.

These values exhibit significant variation between bones: some (such as the 2nd and 3rd phalanges, coxal, and metacarpus) have mean values above or very close to 85% across all taxa, whereas the scapula and malleolar bone reach only a maximum of 74% (**Table 02**).

**Table 02.** Mean across bones of the highest character-state frequency observed for each character. Values are reported for the families and subfamilies included in the study (Caprinae and Moschinae are treated here as equivalents of Bovidae and Moschidae within this specimen sample, respectively).

| Bone | All taxa | Bovidae | Cervidae | Moschidae | Cervinae | Muntiacinae |
| --- | --- | --- | --- | --- | --- | --- |
| 1st Phalanx | 0.86 | 0.83 | 0.86 | 0.91 | 0.91 | 0.77 |
| 2nd Phalanx | 0.91 | 0.92 | 0.91 | 0.96 | 0.92 | 0.88 |
| 3rd Phalanx | 0.88 | 0.93 | 0.85 | 0.96 | 0.84 | 0.86 |
| Large cuneiform | 0.83 | 0.79 | 0.87 | 0.72 | 0.87 | 0.90 |
| Calcaneus | 0.77 | 0.79 | 0.76 | 0.78 | 0.74 | 0.79 |
| Magnum | 0.84 | 0.79 | 0.86 | 0.86 | 0.87 | 0.84 |
| Coxal | 0.90 | 0.90 | 0.88 | 0.99 | 0.90 | 0.86 |
| Navicular-cuboid | 0.82 | 0.75 | 0.84 | 0.98 | 0.85 | 0.81 |
| **Femur** | 0.85 | 0.90 | 0.82 | 0.93 | 0.86 | 0.76 |
| **Unciform** | 0.76 | 0.78 | 0.77 | 0.66 | 0.80 | 0.71 |
| **Humerus** | 0.79 | 0.75 | 0.81 | 0.80 | 0.81 | 0.81 |
| **Large sesamoid** | 0.80 | 0.66 | 0.84 | 1.00 | 0.90 | 0.72 |
| **Lunatum** | 0.82 | 0.78 | 0.83 | 0.89 | 0.84 | 0.82 |
| **Malleolar** | 0.63 | 0.59 | 0.63 | 0.78 | 0.66 | 0.57 |
| **Metacarpus** | 0.88 | 0.91 | 0.85 | 1.00 | 0.86 | 0.84 |
| **Metatarsus** | 0.84 | 0.80 | 0.85 | 0.99 | 0.86 | 0.82 |
| **Patella** | 0.86 | 0.90 | 0.84 | 0.86 | 0.89 | 0.75 |
| **Pisiform** | 0.86 | 0.90 | 0.84 | 0.93 | 0.89 | 0.73 |
| **Pyramidal** | 0.75 | 0.76 | 0.74 | 0.83 | 0.79 | 0.64 |
| **Radius** | 0.81 | 0.80 | 0.82 | 0.82 | 0.83 | 0.79 |
| **Scaphoid** | 0.81 | 0.78 | 0.81 | 0.91 | 0.85 | 0.73 |
| **Scapula** | 0.74 | 0.70 | 0.75 | 0.82 | 0.77 | 0.72 |
| **Astragalus** | 0.76 | 0.70 | 0.77 | 0.89 | 0.77 | 0.77 |
| **Tibia** | 0.79 | 0.79 | 0.77 | 0.93 | 0.78 | 0.75 |
| **Ulna** | 0.80 | 0.71 | 0.82 | 0.95 | 0.85 | 0.76 |
| **Mean** | **0.82** | **0.81** | **0.82** | **0.90** | **0.84** | **0.78** |

When considering families separately, the mean maximum frequency is higher for Moschidae (89.6%) than for Cervidae (82%) and Bovidae (81%). Among Cervidae, Muntiacinae (78%) present stronger character state variations than Cervinae (84%). Overall, these results suggest that, when used in isolation, the recorded characters have limited identification potential. Therefore, a more advanced approach considering multiple characters from a single bone needed to be pursued.

### III) Results of the Random Forest and Naïve Bayes approaches

The analyses we performed combined several characters from single bones but did not combine different bones to simulate an application to the archaeological record, where the bones are most often found isolated.

In general, both the Random Forest (RF) and Naïve Bayes (NB) approaches performed very similarly in our tests. The ability of the analyses to correctly identify a bone is strongly influenced by the taxonomic rank considered. Indeed, correct family-level identification can be achieved on average in 85.6% (NB) or 87.2% (RF) of cases. For subfamily-level identification, this percentage falls to 68.6% (NB) or 69.4%(RF), then to 50.1% (NB) or 50.0% (RF) for genus-level identification, and to 34.9% (NB) or 33.2% (RF) for species-level identification. The number of characters considered (**Figure 80-A**) also has a strong impact on the reliability of the results. Indeed, shifting from one to six input characters improves identification reliability for every taxonomic rank (+13-15% for family level, and +20-22% for the other ranks). There are also differences between bones, as not all anatomical parts have the same discriminatory power. The mean identification accuracy of the different bones (**Figure 80-B**) follows an approximately normal distribution centered on 60–65% accuracy, with some bones presenting mean accuracies below or around 50% (Pyramidal, Unciform, Lunatum, Malleolar bone, Large cuneiform, and Astragalus) and some above or around 65% for both methods (Femur, Humerus, Patella, and 2nd Phalanx). The complete list of mean accuracies for the different ranks, bones, and numbers of characters k is reproduced here to provide a reliability estimate for each possible case (Supplementary Material 06; Mithong et al., 2026).

**Figure 80.**
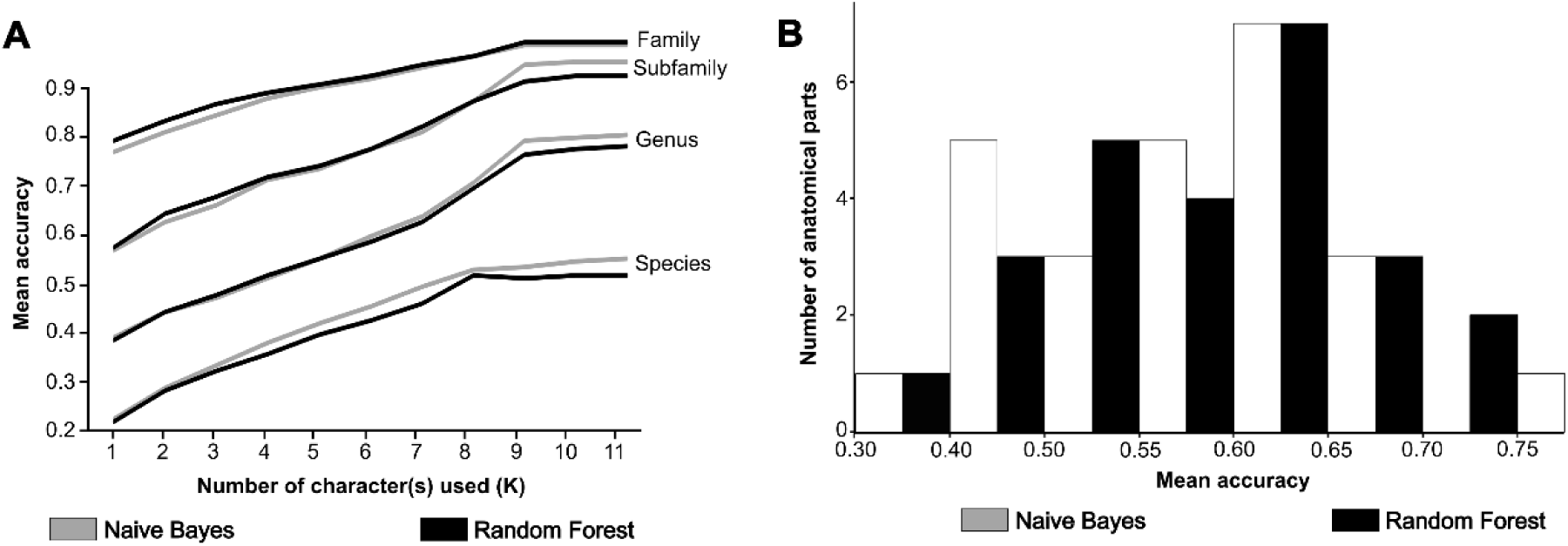
A) Mean accuracy of the identifications for a given number of characters (K) and a given taxonomic rank for both methods. Characters are not mixed between several bones. B) Mean accuracy of the considered anatomical parts for both methods.

There are strong disparities between the identification accuracies of the different taxa, mainly because our dataset was not strictly balanced between the groups considered at the different ranks. At the family rank, Cervidae are identified with 97% (RF) or 98% (NB) accuracy, but the identification of Moschidae (79% RF, 70% NB) and Bovidae (74% RF, 63% NB) is less reliable. The same disparity persists at the subfamily rank, where Cervinae (89% RF, 95% NB) are better identified than Caprinae (77% RF, 69% NB), Moschinae (81% RF, 74% NB), and Muntiacinae (56% RF, 49% NB). Regarding the genera, some can be identified with reasonable confidence, such as *Cervus* (*sensu lato*) (78% RF, 87% NB), *Muntiacus* (62% RF, 59% NB), *Pseudois (73% RF, 67% NB)*, and *Moschus* (84% RF, 80% NB), but identification of the others is more challenging (*Capricornis*: 45–43%, *Naemorhedus: 42–35%, and Axis: 30–23%) or very difficult (Elaphodus: 24–18%)*. Regarding species, with the exception of those from monospecific families/subfamilies (*moschiferus*: 88–85%, and *nayaur*: 76–73%), the others are either difficult (reevesi, axis, and goral: 57–41%), very difficult (*sumatraensis*, *unicolor*, and *muntjak*: 46–25%), or nearly impossible to characterize properly (*nippon*, *cephalophus*, and *eldii*: 29–14%, and porcinus and crispus at 12% or below). The random thresholds being 33% for families, 25% for subfamilies, 12.5% for genera, and 7.1% for species, only in a few cases do the accuracies of the analyses approach these values, and the accuracies for higher taxonomic ranks are especially high. The observed variations in accuracy are mostly explained by imbalances in the number of specimens across the different taxa, but not exclusively. They also reveal the strong morphological similarity between *Capricornis* and *Naemorhedus*, and among the members of the genus *Cervus* (*sensu lato*).

## Discussion

### General remarks regarding the polymorphism of the described characters and their use for taxonomic identification

The multiple analyses carried out on the characters first described on a small set of specimens and tested on a larger comparative sample provide several interesting insights. First, they indicate that although the initially described set of characters allowed the distinction of the specimens used in the initial description phase, the final picture is far more ambiguous. Indeed, the postcranial characters we investigated present very strong degrees of polymorphism, with only one character (131) not being polymorphic at all. This seems to indicate that postcranial elements are far more subject to morphological variations due to sex, ontogeny or other sources of intraspecific morphological variability, than the teeth, antlers, and other skull elements most often investigated for Ruminantia. As a result, the testing phase indicates that the individual identification power of our characters was weak, and that reaching even a decent identification accuracy was only possible through the combination of several characters and the investigation of higher taxonomic ranks. These results differ strongly from those of previous studies that used osteological characters providing, in most cases, very good discriminatory power (Bochaton et al., 2016; Hanot & Bochaton, 2018; Zhang et al., 2022). We therefore had to push the analytical protocol one step further with the Random Forest and Naïve Bayes approaches in order to extract meaningful information from a set of highly variable character states. This methodological approach, not developed in previous similar studies designed for the identification of archaeological remains, could be reproduced to allow the consideration of more variable characters and the comparison of closely related species whose morphologies may strongly overlap. This, however, requires accepting uncertainty in taxonomic identification, something that exists in many bone identification attempts but is most of the time not addressed in the zooarchaeological literature.

Nevertheless, we were still able to detect some rare, and most often variable, synapomorphies (**Table 01**; **Figure 79**), with 11 autapomorphic characters presenting a total of 12 autapomorphic character states, and 37 synapomorphic characters presenting 46 synapomorphic character states. This shows that, in spite of the strong variation we observed, postcranial elements can still offer valuable phylogenetic information and may become useful in systematic studies incorporating fossils, as we observed characters allowing the identification of Bovidae, Cervidae, Moschidae, Muntiacinae, Ovibovina (*sensu* Hassanin et al., 2012), *Muntiacus*, *Pseudois nayaur*, *Elaphodus cephalophus* and *Axis porcinus* within our species sample. There is, of course, a possibility that our results could be improved if new observations and documentation of additional characters were conducted. The initial set of characters tested in this study was developed under strong time constraints, as the authors were not working in an institution hosting specimens, and detailed observations were made on 3D models rather than on the specimens themselves. Yet, as no other identification criteria appeared obvious to us during the testing phase of this study, we do not believe this initial limitation represents a major flaw of the present study. We rather believe that our results show how variable the postcranial anatomy of small and medium-sized Southeast Asian ruminants is, and how difficult it is to distinguish their bones in the case of isolated elements. We believe it is very unlikely that this observation is the result of a methodological flaw.

From a more positive perspective, obtaining such results with a clear and detailed approach represents a significant advance. It confirms that discriminating between similar taxa on the basis of their postcranial elements is, in many instances, impossible using conventional approaches, while providing the tools needed to reach the most detailed identification possible in a clear and reproducible way. Our study argues that there is no point in trying to achieve an artificial and likely erroneous precision in the taxonomic attribution of postcranial elements based on apparent morphological similarities established from a limited set of comparative specimens.

The results presented here will greatly facilitate many zooarchaeological and Quaternary paleontological studies of these bones, as it will no longer be required to conduct subjective observations or to reach a decision based on potentially biased arguments. Most of our characters are easy to observe, and consequently, the identification of the studied taxa no longer requires any particular expertise. This is a strong advance in terms of reproducibility, as the quality and relevance of results can be directly assessed and reproduced; accessibility, as anyone with basic anatomical knowledge and no access to modern bone collections can now attempt to identify the investigated elements; and also a significant time saving, as time-consuming comparisons with modern bones are no longer required.

Hereafter, we provide guidelines for the practical use of our study’s results. To ensure true reproducibility, a broadly similar approach must be followed by all users of these characters, and any deviations from this protocol should be clearly stated at the beginning of the study.

### How to use our characters to study zooarchaeological and paleontological material

The characters described and tested in this study have been defined on a limited number of taxa and thus could not be used to identify bone pieces not belonging to these taxa. The user should thus make sure the bone they are trying to identify belongs to a member of Ruminantia, a basic step that can be done with a straightforward comparison with the pictures included in this paper, 3D models of any taxa available online (e.g., on Morphosource.org), or a comparative skeleton if available. As indicated in the Material and Methods section of this paper, the large Southeast Asian Ruminantia species have been excluded from the present sample and analyzed separately in the next paper of this series (Chantasri et al., 2026). The user should first make certain that they are not trying to identify a member of the genus *Bos*, *Bubalus,* or *Budorcas*, which are taxa larger than the largest taxon included in the present study (*Rusa unicolor*). In case of doubt, this could be assessed using the morphometric data from the first paper of this series (Seesod et al., 2026).

Outside of a few rare exceptions (see Table 01), the characters presented in our study cannot be used directly for taxonomic identification, due to their largely polymorphic distribution among species. To be useful, the character states observed on a subfossil or fossil element must be combined in a statistical framework (in our case, a Naïve Bayes model) to obtain an attribution probability for a given taxon.

This can be performed automatically using the application we developed (Bochaton et al., 2026). The application is a single, self-contained HTML file that requires no installation, and can be opened directly in any standard web browser. The Naïve Bayes models, trained in R as described above, are embedded directly within the application (see Material and Methods section).

Once inside the application, the user selects the bone and inputs any number of observable character states from a single anatomical element. Combinations of characters from different anatomical elements are not allowed, as the models were generated and validated separately for each element. After input, the application provides attribution probabilities at the family, subfamily, genus, and species levels. Each taxonomic rank is calculated independently and involves a different number of possible taxa. The final models deployed in the application are trained on the complete reference dataset, whereas the accuracy reported in the present paper is obtained from models trained on only 80% of the specimens across 1000 independent replicates. The accuracies reported here should therefore be considered a conservative, lower-bound estimate of the identification performance actually achieved by the deployed models, which benefit from a larger reference sample. To further inform users of the reliability of a given identification, the application also flags any taxon represented by fewer than five reference specimens, for which probability estimates should be interpreted with additional caution. It also provides several pieces of information to help the reader assess the quality and reliability of the obtained attribution.

Because character states are rarely unique to a single species, results will often be ambiguous, reflecting both morphological proximity among taxa and the number of reference specimens available for each taxon (i.e., for a given frequency of character states, a taxon that is more represented in our sample will be indicated as more likely than a rarely represented one). Probabilities should be read against the values expected under random attribution, which differ widely between ranks in this dataset: 33.3% for the three families, 25% for the four subfamilies, 12.5% for the eight genera, and 7.1% for the fourteen species. A probability close to its baseline therefore carries no information at all. How much a probability above that baseline is worth is addressed below.

The tests reported in the Material and Methods section show that an attribution is correct in more than 90% of cases from 80% upwards at the family, subfamily and genus ranks, and from 95% upwards at the species rank (Supplementary Material 04, Mithong et al., 2026). In order to retain the most precise attribution that can be relied upon, the following rules should be applied: 1) retain the lowest rank whose attribution reaches the threshold set for that rank, and 2) record the element as Ruminantia indet. where none does. In our tests, the species rank is reached for 10.2% of the character combinations tested, the genus rank for 19.7%, the subfamily rank for 37.2%, the family rank for 75.4%, and an attribution retained under these rules proved correct in at least 97% of cases at every rank. An isolated element of a small or medium-sized ruminant will therefore usually be assigned no lower than the family or the subfamily, but the attribution retained can be considered reliable. As indicated by our tests, neither the margin separating the two most likely taxa at a given rank nor the mutual consistency of the attributions returned at the four ranks adds any reliability once these thresholds are applied. These rules are applied automatically by the application, which highlights the lowest rank that can be retained, or highlights none where no attribution meets its threshold, and which displays beside every rank the reliability observed for the probability returned. In specific cases where cranial and/or dental remains attest to the occurrence of a given taxon in the assemblage, less likely generic and specific attributions may be considered, but these are not strong enough to prove the occurrence of that taxon by themselves.

If necessary, the character-based results can be complemented with the metrical criteria developed in the first paper of this series (Seesod et al., 2026) to reduce ambiguity using size data. This may be particularly useful for morphologically similar taxa that differ in size (e.g., Cervini). In such cases, it should be clearly stated that the specific identification is based on size criteria and is therefore subject to the potential biases previously discussed (Seesod et al., 2026). Whatever approach is adopted, it should be clearly described in the Material and Methods section of the study.

Ultimately, these tools can be used to help interpret bone assemblages in which several elements can be considered together to better assess the reliability of the occurrence of a given taxon. For instance, in the case of an assemblage including cranial elements or very distinctive post-cranial pieces, the probabilistic results of our approach could be interpreted in the light of the results obtained on the other elements composing the assemblage. In such cases, attributing additional pieces to a taxon whose occurrence in the assemblage is considered reliable might become the working hypothesis for the more ambiguous pieces. As such, the tools presented in this series of preprints help assess the reliability of taxonomic attributions and provide a means to make the study more reproducible in all cases, regardless of the presumed experience or expertise of the user.

## Conclusion

Although the results presented here highlight the difficulties of reaching a precise taxonomic identification of small and medium-sized Southeast Asian ruminants based on their postcranial elements, our approach still represents a significant step forward for the quality of Late Quaternary studies, particularly regarding zooarchaeological studies. Indeed, the results of these works must, in many instances, rely on the difficult-to-identify bones described in the present study, for which no solid framework was previously available. Our work highlights the limits of what can reasonably be achieved in terms of morphological identification and provides a complete toolkit to perform taxonomic identification in a repeatable and refutable way. These results can also be combined with the metrical approach presented in the first paper of this series to further refine identification, or to discuss size in relation to morphology when the characters provide a confident identification. This approach could also help identify bones relevant for further analyses, particularly molecular approaches, either to gather additional information (e.g., isotopic analyses) or to obtain greater precision regarding taxonomic identification (e.g., ancient DNA). Beyond these technical applications, we hope that such an approach will be reproduced in other regions and for other taxa, as this is an essential step towards reproducibility and transparency in taxonomic identification in zooarchaeological studies. This will also make such data more relevant for advanced studies of past environments and human-environment interactions, which require reliable baseline taxonomic data, especially in biodiversity-rich areas that remain poorly documented from these perspectives.

## Acknowledgements

The authors are grateful to the museums, universities, and staff members who supported this study. We especially thank the Muséum national d’Histoire naturelle (Paris, France) and Joséphine Lesur; the National Museum of Natural History (Washington, D.C., USA) and Darrin Lunde; the Zoologische Staatssammlung München (Munich, Germany) and Anneke H. van Heteren; the Museum für Naturkunde (Berlin, Germany) and Christiane Funk; and the Royal Belgian Institute of Natural Sciences (Brussels, Belgium) and Olivier Pauwels.

## Data, scripts, code, and supplementary information availability

Supplementary information is available online: https://doi.org/10.23708/VVFRZX; Mithong et al., 2026.

## Author contributions

**Conceptualization**: Supalak Mithong and Corentin Bochaton; **Data curation**: Supalak Mithong and Corentin Bochaton; **Formal analysis**: Supalak Mithong, Corentin Bochaton, and Julien Claude; **Funding acquisition**: Corentin Bochaton; **Data acquisition**: Supalak Mithong; **Supervision**: Corentin Bochaton; **Writing original draft**: Supalak Mithong, and Corentin Bochaton; **Writing – review & editing**: All the authors.

